# Aperiodic neural dynamics define a novel signature of glioma-induced excitation-inhibition dysregulation

**DOI:** 10.1101/2025.05.23.655626

**Authors:** Youssef E. Sibih, Abraham O. Dada, Emily Cunningham, Niels Olshausen, Jasleen Kaur, Velmurugan Jayabal, Sena Oten, Sanjeev Herr, Cesar Nava Gonzales, Andy Daniel, Saritha Krishna, Vardhaan S. Ambati, Alexander A. Aabedi, Gray Umbach, Kanish Mirchia, Poortata Lalwani, Edward F. Chang, David R. Raleigh, Srikantan Nagarajan, David Brang, Shawn L. Hervey-Jumper

## Abstract

Diffuse gliomas remodel neuronal circuits with prognostic and therapeutic significance for patients. Electrophysiologic measures of cortical excitability hold promise for monitoring disease progression and evaluating therapeutic responses. The power law exponent (aperiodic slope) reflects the balance between excitatory and inhibitory activity within neuronal networks, a critical aspect of normal brain function often disrupted in neurological conditions. Despite its potential, the significance of the aperiodic slope in glioma-infiltrated tissue and its underlying cellular processes has not been fully investigated. Here, we integrate multi-modal electrophysiological analysis with transcriptomic profiling to analyze the aperiodic slope in both normal and glioma-infiltrated cortex. We determine that glioma infiltration induces a flattening of the aperiodic slope, indicating a shift toward excitation dominance that varies according to tumor subtype and correlates with impairments in semantic naming. Single-nucleus RNA sequencing revealed that cortical regions with flat aperiodic slope exhibit transcriptional programs enriched in glutamatergic signaling, membrane depolarization, and excitatory synaptic transmission. The aperiodic slope responds to pharmacologically induced changes in cortical inhibition during propofol administration, a GABA_A_ agonist. Our results establish the aperiodic slope as a robust biomarker of glioma-associated excitation-inhibition imbalance, with potential applications in tumor classification and treatment monitoring.

## INTRODUCTION

The balance between excitation and inhibition (E/I) within cortical circuits is fundamental to maintaining neural homeostasis and supporting essential functions such as information processing, sensory integration, and motor control^1–4^. Disruptions to this balance have been implicated in a range of neurological and neuropsychiatric conditions, including epilepsy, autism spectrum disorder, and schizophrenia, highlighting its importance in both physiological and pathological states^5–11^. Recently, the role of E/I dysregulation in the initiation, proliferation, invasion, and treatment resistance of diffuse glioma has garnered increasing attention^12–14^. Gliomas are the most common primary brain malignancies and exhibit a unique capacity to remodel the neuronal microenvironment through direct synaptic integration, the release of excitatory paracrine factors, and the loss of inhibitory neuronal populations^12,15,16^. Glioma-induced neuronal remodeling leads to circuit-level hyperexcitability, which manifests clinically as seizures and impaired cognitive function, and has been linked to regional immunosuppression and shortened overall survival^12,17–19^. However, our ability to characterize the extent and nature of glioma-driven neuronal network dysfunction in the human cortex remains limited by the absence of a physiologically grounded, quantitative measure of circuit-level excitability.

Recent studies have implicated multiple mechanisms governing glioma-induced local disruption of E/I dynamics, including aberrant glutamatergic transmission, altered GABAergic signaling, shifts in chloride homeostasis, and fluctuations in extracellular ion gradients^20–22^. Although a more excitation- or inhibition-dominant state may reflect disease biology, quantifying cumulative E/I balance in vivo remains challenging, where direct measurement of synaptic activity is limited. A promising proxy for local E/I dynamics lies in the aperiodic exponent of the power spectral density (PSD) of the electrophysiological signal^23–26^. The aperiodic (non-oscillatory) component of the electrophysiological signal exhibits a characteristic 1/f-like form, in which power declines with increasing frequency. The rate of this decline (the ‘aperiodic exponent’, equivalent to the absolute value of the slope of the log-transformed PSD) is reported to reflect shifts in synaptic excitation and inhibition in both animal models and human cortical recordings^23,27–29^. In this framework, larger exponents (steeper negative slopes) are associated with increased inhibitory tone, while smaller exponents (flatter negative slopes) suggest elevated excitatory tone. The aperiodic exponent can be extracted from subdural array local field potentials in both resting-state and task-based electrocorticography (ECoG) using specparam (formerly FOOOF, or “Fitting Oscillations and One Over F” algorithm)^25^ and similar programs^30^. Specparam decomposes the power spectral density (PSD) into periodic and aperiodic components, facilitating the characterization of the underlying neural state. Despite its growing use, the biological validity of the aperiodic exponent as a measure of E/I balance remains unresolved. Recent evidence challenges the interpretability of the aperiodic slope as a direct readout of the excitation-inhibition (E/I) relationship, reporting that spectral slope did not consistently track the ratio of excitatory to inhibitory synaptic conductance^31^. In glioma, where synaptic remodeling profoundly disrupts excitatory-inhibitory (E/I) dynamics, electrophysiological measures of cortical excitation in the human brain may reflect the behavior of malignant cells. We therefore test the hypothesis that the aperiodic exponent in glioma-infiltrated tissue is pharmacologically responsive, tumor subtype-specific, and reflects excitatory transcriptional programs.

In this research, we utilize high-resolution subdural array recordings from patients diagnosed with World Health Organization (WHO) grades 2-4 diffuse gliomas, including oligodendroglioma, astrocytoma, and glioblastoma, to investigate the cortical aperiodic exponent in both glioma-affected and unaffected regions. Recordings were obtained intraoperatively from awake, behaving patients during resting states and task-evoked activities. Given that the aperiodic exponent is thought to reflect the excitation-inhibition balance at the neural circuit level, we explored its cellular and molecular underpinnings. By conducting molecular profiling of resected cortical tissue and correlating these data with electrode locations, we linked spectral changes to specific gene expression patterns and pathway activity modules. This approach provides a comprehensive, multiscale framework for understanding glioma-induced disruptions in neural circuitry, integrating electrophysiological, behavioral, and molecular analyses. Our objectives were to determine whether the aperiodic exponent differentiates between glioma-infiltrated and normal tissue, and whether it varies across different molecular glioma subtypes. We also evaluated whether the exponent responds to pharmacologically induced modifications in cortical inhibition, supporting its utility in tumor classification. Additionally, we investigated whether task-related fluctuations in the exponent can predict language performance on a trial-by-trial basis across tumor types. Lastly, using single-nucleus RNA sequencing and immunofluorescence on spatially matched cortical samples, we identified transcriptomic correlates of the exponent, linking spectral flattening to excitatory remodeling in both non-neoplastic and glioma-affected tissue. Collectively, these findings position the aperiodic exponent as a physiologically and molecularly relevant biomarker of neuronal network dysfunction associated with gliomas.

## RESULTS

### Experimental Framework for Electrophysiologic Analysis

We aimed to determine whether the aperiodic exponent—an electrophysiologic measure of complex brain electrical activity—may be computed in glioma-infiltrated cortex and serve as a reliable measure of excitatory-inhibitory (E/I) imbalance. We analyzed subdural array local field potentials obtained using intraoperative ECoG recordings from 13 patients (38.4% female) with newly diagnosed WHO 2-4 gliomas, focusing on the power spectrum’s aperiodic (1/f) exponent across glioma-infiltrated and normal-appearing cortical regions. The cohort included patients diagnosed with primary oligodendroglioma, IDH-mutant, 1p/19q codeleted, WHO grade 2 (n=3), astrocytoma, IDH-mutant, WHO grade 2 and 3 (n=5), and glioblastoma, IDH-wildtype, WHO grade 4 (n=5) (**Fig. 1A, Extended Data Fig. 1**, **Table 1**). The mean age across all patients was 47.3 years (range, 27.7 to 68.4 years). Low-density or high-density subdural microarrays (ranging from 4×5 to 8×12 configurations, respectively) were placed directly on the cortical surface, over normal-appearing and glioma-infiltrated regions, during both resting-state and language tasks. Tumor locations included the frontal lobe (n=4), temporal lobe (n=2), parietal lobe (n=3), and insular language/sensory-associated regions (n=4). Across the cohort, 660 subdural electrodes were included for analysis. Each electrode was anatomically co-registered to T1 post-gadolinium and T2 FLAIR MRI, and glioma-infiltration status was identified based on electrode spatial overlap with hyperintense signal on FLAIR imaging and the absence of central contrast enhancement on T1-post-gadolinium sequences^32^ (**Fig. 1B**). 518 electrodes overlaid normal-appearing cortex, and 142 overlaid glioma-infiltrated cortex (**Extended Data Fig. 2**).

**Fig. 1.**
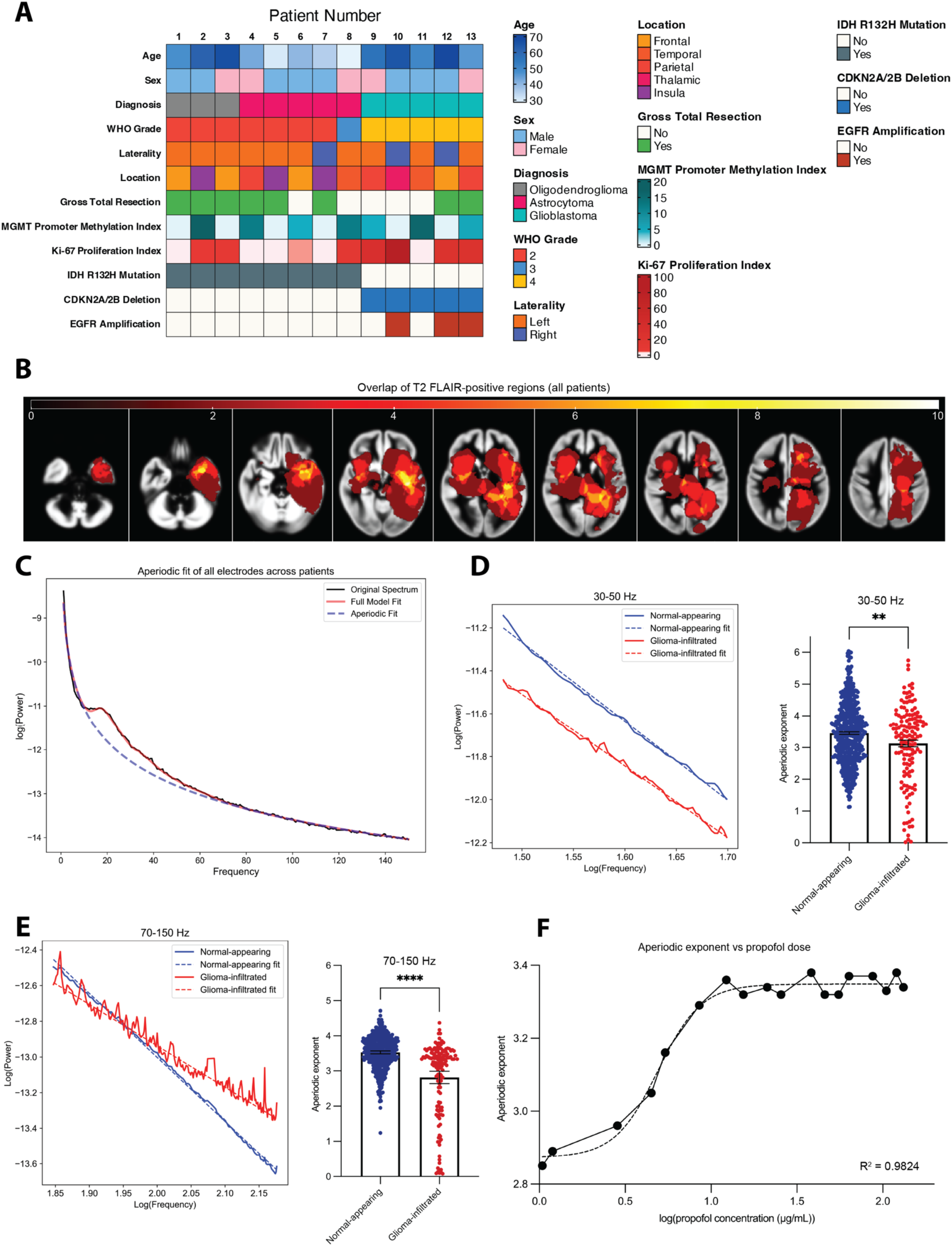
Glioma-associated Alterations in Cortical Aperiodic Activity and Molecular Features. The study cohort consists of 13 patients diagnosed with oligodendroglioma, IDH-mutant and 1p/19q-codeleted, CNS WHO grade 2 (n=3, 23.0%), astrocytoma, IDH-mutant, CNS WHO grade 2 and 3 (n=5, 38.5%), and glioblastoma, IDH-wildtype, CNS WHO grade 4 (n=5, 38.5%). Spectral analysis was performed on 660 electrodes, comprising of both normal-appearing (n=518, 78.5%) and glioma-infiltrated (n=142, 21.5%) sites. **A**, Oncoprint displaying the clinical, histopathological, and genetic features of patients in cohort. Clinical variables include age, sex, WHO grade, glioma subtype, and extent of resection. Histopathological variables include proliferation index (Ki-67) and MGMT promoter methylation index. **B,** Overlay of FLAIR-positive regions on preoperative MRI T2 FLAIR across all patients. The heatmap represents the normalized spatial overlap of FLAIR-positive areas indicative of glioma-infiltrated cortex, with warmer colors signifying regions of higher overlap. **C,** Spectral analysis and aperiodic exponent fit of all electrodes across all patients analyzed in this study during resting state. The black line illustrates the original neural power spectrum, and the red line displays the full model fit. The aperiodic fit is represented by the blue dashed line. **D,** Left: Log-log aperiodic fits analyzed in the gamma frequency range (30-50 Hz) for all glioma-infiltrated (red) and normal-appearing (blue) electrodes during resting state. Right: Bar plot of average aperiodic exponent magnitudes comparing normal-appearing and glioma-infiltrated electrodes at 30-50 Hz. **E**, Left: Log-log aperiodic fits analyzed in the high gamma frequency range (70-150 Hz) for all glioma-infiltrated and normal-appearing electrodes during resting state. Right: Bar plot of average aperiodic exponent magnitudes comparing normal-appearing 2.813 (range 1.24-4.71) and glioma-infiltrated 3.525 (range 0.8-4.37) electrodes. Glioma-infiltrated electrodes have significantly flatter aperiodic exponents than normal-appearing electrodes. Each dot represents an electrode; error bars represent the 95% confidence interval. Comparisons were conducted with linear mixed effects models incorporating a patient-level random effect **p≤0.01, ****p≤0.0001. **F**, Sigmoidal regression modeling of the relationship between aperiodic exponent and average propofol dose (log(μg/mL)) in a patient’s body during surgery at a given time-point. Dots represent average aperiodic exponent across all patients computed in their respective concentration bin. Dashed line represents regression fit. A significant positive correlation (sigmoidal fit R^2^ = 0.9824, 95% CI Top: 3.33-3.37, Bottom: 2.68-2.90) suggests that a steeper aperiodic exponent correlates with a higher propofol dose. Abbreviations: WHO = World Health Organization, MGMT = O6-methylguanine-DNA methyltransferase, O = oligodendroglioma, A = astrocytoma, GBM = glioblastoma multiforme

**Table 1.** Cohort Characteristics. Summary of patient demographics, tumor pathology, and electrode classifications for the study cohort (n = 13 patients, n = 660 cortical electrodes). *PN = picture naming, AN = auditory naming. MGMT = O6-methylguanine-DNA methyltransferase*.

| Variable | Study Cohort (n = 13 patients, n = 660 Electrodes) |
| --- | --- |
| **Age at diagnosis (years), mean (range)** | 47.3 (27.7 – 68.4) |
| **Sex, n (%)** |  |
| Female | 5 (38.4%) |
| Male | 8 (61.6%) |
| **Language Task Distribution, n (%)** |  |
| Patients with ≥ 1 Incorrect PN Trial(s) | 9 (69.2%) |
| Patients with 0 Incorrect PN Trials | 4 (30.8%) |
| Patients with ≥ 1 Incorrect AN Trial(s) | 9 (75.0%) |
| Patients with 0 Incorrect AN Trials | 3 (25.0%) |
| **Tumor Location, n (%)** |  |
| Laterality | Left: 10 (76.9%), Right: 3 (23.1%) |
| Temporal | 2 (15.4%) |
| Frontal | 4 (30.8%) |
| Parietal | 3 (23.1%) |
| Insula | 3 (23.1%) |
| Thalamic | 1 (7.7%) |
| **Gross Total Resection, n (%)** | 8 (61.5%) |
| **Tumor Diagnosis, n (%)** |  |
| Oligodendroglioma, IDH-Mt and 1p/19q-codeleted | 3 (23.0%) |
| Astrocytoma, IDH-Mt, WHO 2-3 | 5 (38.5%) |
| Glioblastoma, IDH-Wt, WHO 4 | 5 (38.5%) |
| **CNS WHO Grade, n (%)** |  |
| 2 | 7 (53.8%) |
| 3 | 1 (7.8%) |
| 4 | 5 (38.4%) |
| **Ki-67 Proliferation Index %, mean (range)** | 15.7% (2.0 – 85.0) |
| MGMT Promoter Methylation Index, mean (range) | 5.0 (0.0 – 17.0) |
| **Total Electrodes of Cohort (n = 660)** |  |
| Glioma-infiltrated (%) | 142 (21.5%) |
| Normal-appearing (%) | 518 (78.5%) |
| Oligodendroglioma Electrodes (%) | 104 (15.8%) |
| Astrocytoma Electrodes (%) | 285 (43.2%) |
| Glioblastoma Electrodes (%) | 271 (41.0%) |
| **Normal-appearing Electrodes (n = 518)** |  |
| Oligodendroglioma (%) | 82 (15.8%) |
| Astrocytoma (%) | 200 (38.6%) |
| Glioblastoma (%) | 236 (45.6%) |
| **Glioma-infiltrated Electrodes (n = 142)** |  |
| Oligodendroglioma (%) | 22 (15.5%) |
| Astrocytoma (%) | 80 (56.3%) |
| Glioblastoma (%) | 40 (28.2%) |

### Spectral Decomposition and Selection of Frequency Bands for Aperiodic Analysis

To establish the spectral features of cortical activity across the study cohort, we first computed the average power spectral density (PSD) for all electrodes and patients in the 0-150 Hz frequency range (**Fig. 1C**) during resting-state ECoG using the specparam algorithm. ^25^ The full-range decomposition revealed the expected broadband aperiodic decay in semi-log space, confirming the validity of using specparam to extract the aperiodic exponent in these data. Following this, we extracted aperiodic exponent values from two frequency bands: gamma (30-50 Hz) and high-gamma (70-150 Hz; **Extended Data Fig. 3**). Both of these are ranges that have been used for quantification of aperiodic exponent in the past^23,27,33^. When measuring aperiodic exponent, the goal is to capture changes in slope over a uniform range while avoiding contamination from low-frequency oscillatory activity, as well as any ‘knee’ or bend in the signal. The 30-50 Hz range is often applied in scalp EEG contexts and has been used in recent publications to demonstrate the aperiodic exponent as a proxy for E/I balance in human and animal computational models^23^. The 70-150 Hz range has been used in ECoG contexts to measure the aperiodic exponent above a knee in the spectrum identified around 70-80 Hz^33^. Beyond this range, the spectrum typically exhibits scale-free behavior (the aperiodic exponent does not change as a function of measurement range). Although the 30-50 Hz range provides continuity with recent investigations of the relationship between the aperiodic exponent and E/I balance, the 70-150 Hz range may provide a more accurate measurement of the aperiodic component of the signal. Accordingly, we focus the majority of our analysis on measurements drawn from the higher-frequency band.

### Glioma Infiltration Is Associated with Frequency-Independent Flattening of Aperiodic Activity

We next compared aperiodic exponent magnitudes between normal-appearing and glioma-infiltrated electrodes at both the 30-50 Hz and 70-150 Hz frequency range (**Fig. 1D-E**). Across the cohort, electrodes overlying glioma-infiltrated cortex exhibited significantly flatter aperiodic exponents compared to those over normal-appearing cortex in both frequency ranges. In the 30-50 Hz band (**Fig. 1D**), the mean exponent for glioma-infiltrated electrodes was 3.124 (range 0.63-5.75) compared to 3.448 (range 1.13-6.04) in normal-appearing cortex. Similarly, in the 70-150 Hz band (**Fig. 1E**), glioma-infiltrated cortex exhibited a mean exponent of 2.813 (range 0.8-4.37) versus 3.525 (range 1.24-4.71) in normal-appearing cortex. The differences in each band were statistically significant in both frequency ranges, based on linear mixed-effects models with the Satterthwaite approximation, which included patient ID as a random intercept to account for within-subject correlation due to repeated electrode sampling. For the 30-50 Hz band, glioma-infiltrated cortex was associated with a significantly flatter aperiodic exponent compared to normal-appearing cortex (β = −0.21, 95% CI: −0.36 to −0.05, SE = 0.078, t(654) = −2.65, p = 0.008). The model accounted for between-subject variability with a random intercept for patient ID (standard deviation = 0.84), and residual variance had a standard deviation of 0.70. For the 70-150 Hz band, glioma-infiltrated cortex showed a significantly flatter aperiodic exponent compared to normal-appearing cortex (β = −0.51, 95% CI: −0.60 to −0.42, SE = 0.046, t(653) = −11.13, p < 0.0001). The random intercept for patient ID captured between-subject variability (SD = 0.54), and the residual standard deviation was 0.41. These results demonstrate that glioma infiltration is associated with a robust flattening of the aperiodic exponent across both frequency ranges, consistent with a shift toward excitation dominance in the affected cortex.

### Spectral Steepness Reflects Pharmacologically Induced Shifts in E/I Balance

To assess whether the aperiodic exponent reflects pharmacologically driven shifts in cortical inhibition, we examined its relationship to propofol concentration. Propofol, a GABA_A_ agonist, enhances inhibitory tone^34,35^ and has been shown in computational and primate models to steepen the aperiodic exponent with increasing doses^36–39^. Additionally, studies in healthy young adults have reported that the aperiodic exponent increases with propofol-induced unresponsiveness^27^. However, this relationship has not been tested in older adults and patients with glioma, where the cortical environment is fundamentally altered. In our cohort, we estimated instantaneous propofol concentration using a one-compartment pharmacokinetic decay model aligned to anesthetic discontinuation during a patient’s emergence from anesthesia. We then binned log-transformed propofol concentrations and computed the corresponding mean aperiodic exponent across the cohort for each bin. A clear sigmoidal relationship was observed, with the exponent increasing (i.e., the steepening of the negative slope of the PSD) as the modeled propofol concentration increased (**Fig. 1F, Extended Data Fig. 4**). Nonlinear regression yielded an excellent fit (R² = 0.9824), with asymptotes of 2.68–2.90 (lower) and 3.33–3.37 (upper). Notably, no such relationship was observed during dexmedetomidine administration, suggesting that the aperiodic exponent specifically reflects GABAergic modulation rather than general sedation or anesthetic depth (**Extended Data Fig. 4C**). These findings extend prior work by demonstrating that the aperiodic exponent remains sensitive to pharmacologically induced shifts in E/I balance even in the context of glioma-related network remodeling.

### Aperiodic Exponent Differentiates Molecularly Defined Glioma Subtypes

Having established that glioma-infiltrated cortex is associated with flattening the aperiodic exponent, suggestive of an excitation-dominant state, we next set out to determine how much this feature varies across diffuse glioma subtypes. Tissue samples were separated according to WHO 2021 diagnostic criteria^40^ into: (1) oligodendroglioma, IDH-mutant, 1p/19q-codeleted, (2) Astrocytoma, IDH-mutant, and (3) glioblastoma, IDH-wildtype. Embedded within neuronal networks, low- and high-grade gliomas promote neuronal hyperexcitability through activity-dependent paracrine signaling, as well as direct synaptic inputs to neurons and inhibitory neuron loss. However, the relative balance of excitatory and inhibitory inputs within and between molecular subgroups remains poorly understood. Violin plots of electrode-level aperiodic exponents in the high gamma frequency band (70-150 Hz) were generated to reveal glioma subtype-specific differences, with a progressive flattening of the slope from oligodendroglioma to astrocytoma to glioblastoma (**Fig. 2A**). Combined electrodes from oligodendroglioma and astrocytoma samples exhibited the largest exponents (steepest negative slopes), while glioblastoma electrodes displayed markedly flatter values. The glioma subtype distribution was evident across subtypes, as indicated by the median and interquartile ranges. To quantify subtype-specific differences in aperiodic activity, we calculated the mean aperiodic exponent for each electrode in the high-gamma band, extracted from ECoG data. Oligodendroglioma electrodes had a total average aperiodic exponent of 3.68, astrocytoma of 3.55, and glioblastoma of 3.01. We fit a linear mixed-effects model with glioma subtype as a fixed effect and patient ID as a random intercept, using oligodendroglioma as the reference group. Glioblastoma was associated with significantly flatter aperiodic slopes than oligodendroglioma (β = –0.73, 95% CI: –1.37 to –0.10, p < 0.0001). Additionally, astrocytoma had significantly flatter exponents from oligodendroglioma (−0.90, 95% CI: −0.73 to –0.54, p < 0.01). The model accounted for inter-patient variability with a random intercept (SD = 0.43), and the residual standard deviation was 0.45. These findings indicate that increasing tumor grade is associated with an incremental flattening of the aperiodic exponent, with glioblastoma exhibiting the flattest exponents, suggesting a greater excitation-dominant state when compared with IDH mutant gliomas (**Fig. 2B, left**).

**Fig. 2.**
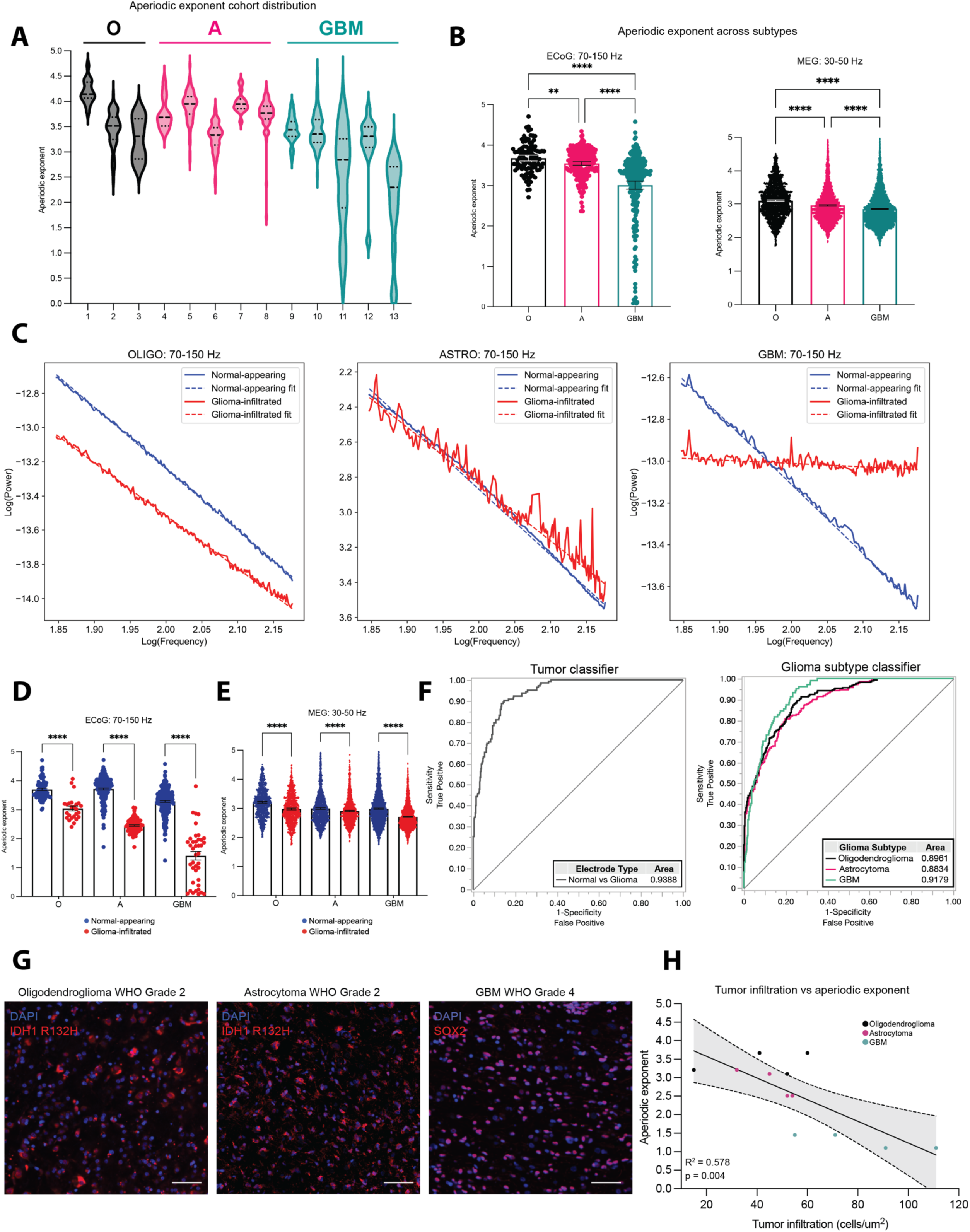
Aperiodic exponent differences across glioma subtypes and classification performance. **A,** Violin plots of ECoG aperiodic exponent values in the 70-150 Hz frequency range grouped by glioma subtype across patients: Dashed lines within violins represent median, first quartile, and third quartile. **B,** (Left) Bar plot comparisons of ECoG aperiodic exponent magnitudes across glioma subtypes at 70-150 Hz, (Right) Bar plot comparisons of aperiodic exponent magnitudes within patient-matched magnetoencephalography (MEG) validation set at 30-50 Hz. Individual dots represent an electrode. Linear mixed effects modeling of ECoG data revealed GBM, and astrocytoma had a significantly flatter average aperiodic exponent compared to oligodendroglioma (GBM 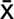=3.01 vs oligodendroglioma 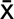=3.68, ****p≤0.0001; astrocytoma 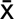=3.55 vs oligodendroglioma 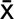=3.68, p=0.0052). Further, GBM had a significantly flatter average aperiodic exponent compared to astrocytoma (GBM 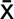=3.01 vs astrocytoma 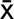=3.55; ****p≤0.0001). **C,** Log-log aperiodic fits for normal-appearing (blue) and glioma-infiltrated (red) electrodes across glioma subtypes. **D,** Comparison of ECoG average aperiodic exponent between normal-appearing (blue) and glioma-infiltrated (red) electrodes for each glioma subtype at 70-150 Hz. Across all glioma subtypes in this study, glioma-infiltrated electrodes were found to have significantly flatter mean aperiodic exponents compared to normal-appearing electrodes (O: 3.04 vs 3.71, A: 2.46 vs 3.70, GBM: 1.40 vs 3.29; p≤0.0001). Bars represent mean aperiodic exponent magnitudes, and error bars represent 95% confidence intervals. Linear mixed effects modeling, ****p≤0.0001. **E,** Comparison of average aperiodic exponent between normal-appearing and glioma-infiltrated regions within MEG analysis at 30-50 Hz. Across all glioma subtypes in this study, glioma-infiltrated electrodes were found to have significantly flatter mean aperiodic exponents compared to normal-appearing electrodes (O: 2.98 vs 3.21, A: 2.91 vs 3.00, GBM: 2.99 vs 2.71; p≤0.0001). Bars represent mean aperiodic exponent magnitudes, and error bars represent 95% confidence intervals. Linear mixed effects modeling, ****p≤0.0001. **F,** ROC curves for bootstrap random forest classifiers predicting electrode tissue type (left), glioma subtype (right: O vs A vs GBM) using aperiodic exponent values. A “leave one patient out” cross-validation approach was utilized for each classifier. **G,** Representative immunofluorescence images from oligodendroglioma WHO 2 (left), astrocytoma WHO 2 (middle), and GBM (right). Tumor cells were labeled by IDH1 R132H or SOX2 immunostaining (red), with DAPI counterstaining (blue). For each sample, tumor cell density (cells/μm²) was quantified from a 20x field of view and normalized to tissue area, scale bar 50 μm. **H,** Aperiodic exponents were assigned to each sample based on the nearest electrode using a Euclidean distance-matching approach. Tumor cell density correlated inversely with the aperiodic exponent (n = 12 samples, R² = 0.578, p = 0.004). This relationship was consistent across glioma subtypes. Abbreviations: O = oligodendroglioma, A = astrocytoma, GBM = glioblastoma multiforme

To validate subtype-specific spectral variations in the ECoG subdural recordings, we subsequently analyzed a magnetoencephalography (MEG) dataset from the same patients included in the ECoG analysis. We created tumor volume masks and stratified MEG voxels based on their spatial overlap with glioma-infiltrated cortex. We used voxels in the contralateral, non-affected hemisphere to define normal appearing. While ECoG analyses focus on the 70-150 Hz high gamma band, this frequency range is not reliably captured in MEG due to biophysical limitations, such as a reduced signal-to-noise ratio and cortical signal attenuation at higher frequencies ^41–43^. As a result, we performed MEG decomposition in the 30-50 Hz band, a frequency range commonly used in noninvasive EEG/MEG studies and previously employed to illustrate shifts in the excitation-inhibition balance. ^23,25^ To assess global differences in the aperiodic exponent across glioma subtypes using MEG, we computed the mean aperiodic exponent for each voxel and grouped them by glioma subtype. The average aperiodic exponent was 3.19 for oligodendroglioma, 2.92 for astrocytoma, and 2.85 for glioblastoma (**Fig. 2B, right**). We fit a linear mixed-effects model with glioma subtype as a fixed effect and subject ID as a random intercept, using oligodendroglioma as the reference group. Both astrocytoma (β = –0.27, 95% CI: –0.69 to 0.15, p = 0.0023) and glioblastoma (β = –0.34, 95% CI: –0.76 to 0.08, p = 0.0011) exhibited significantly smaller aperiodic exponents (flatter slopes) compared to oligodendroglioma. The model accounted for inter-subject variability with a random intercept (SD = 0.27), and the residual standard deviation was 0.45.

To determine whether the observed glioma subtype-dependent differences in the aperiodic exponent were confined to glioma-infiltrated cortex or extended more broadly, impacting normal-appearing cortical regions, we designated each electrode as normal-appearing or glioma-infiltrated and stratified the aperiodic exponent by both glioma subtype. This comparison was motivated by recent work demonstrating that low- and high-grade gliomas can promote long-range changes in synaptic structure, functional connectivity, and neuronal excitability^15,44^. Given that gliomas integrate into cortical networks through synaptic coupling with neurons^20^ and that glioma grade correlates with the degree of excitatory modeling^21,22,45,46^, we reasoned that glioma subtype might modulate both the power and spatial representations of spectral flattening. Across all gliomas, electrodes overlying glioma-infiltrated cortex exhibited a consistently flatter aperiodic exponent at high gamma relative to normal-appearing cortex, with the magnitude of flattening varying by tumor type (**Fig. 2C**). The difference was most pronounced in glioblastoma, where glioma-infiltrated electrodes exhibited markedly flatter aperiodic exponents compared to normal-appearing cortex and lower-grade gliomas. The same comparisons were performed between 0-150 and 30-50 Hz to facilitate comparison with prior slope-fitting studies^23,33^ (**Extended Data Fig. 5**). To quantify the magnitude and consistency of differences in aperiodic exponent between glioma-infiltrated and normal-appearing cortex within each tumor subtype, we compared exponent values stratified by both tissue classification and tumor type (**Fig. 2D**). In oligodendroglioma, glioma-infiltrated electrodes exhibited a mean aperiodic exponent of 3.04 versus 3.71 in normal-appearing cortex. Astrocytoma demonstrated a larger difference, with glioma-infiltrated electrodes averaging 2.46 versus 3.70 in normal tissue. Glioblastoma showed the most pronounced flattening, with glioma-infiltrated regions exhibiting a mean exponent of 1.40 compared to 3.29 in normal-appearing electrodes. Linear mixed-effects modeling with patient ID as the random effect and tissue cortex type and glioma subtype as fixed effects revealed a significant interaction between cortex tissue type (glioma-infiltrated vs. normal-appearing) and glioma subtype (p < 0.001). Specifically, glioma-infiltrated cortex exhibited a significantly flatter aperiodic exponent compared to normal-appearing cortex in glioblastoma (β = −1.37, 95% CI: −1.56 to −1.18, p < 0.0001), oligodendroglioma (β = −0.61, 95% CI: −0.80 to −0.42, p < 0.0001), and astrocytoma (β = −0.30, 95% CI: −0.41 to −0.19, p < 0.0001). These findings demonstrate that the aperiodic exponent is tumor subtype-specific and influenced by tumor infiltration, with glioblastoma-infiltrated cortical electrodes exhibiting the smallest exponent (flattest slope) compared to normal-appearing regions.

Following the discovery of ECoG glioma subtype-specific shifts in the aperiodic exponent (**Fig. 2B, right**), we set out to validate tissue subtype and tissue-specific exponent using MEG. To assess whether glioma subtype modulates the effect of glioma infiltration on aperiodic activity, we fit a linear mixed-effects model with cortex type (glioma-infiltrated vs. normal-appearing), glioma subtype, and their interaction as fixed effects, and subject ID as a random intercept (**Fig. 2E**). In oligodendroglioma, the mean aperiodic exponent was 3.21 in normal-appearing and 2.98 in glioma-infiltrated cortex. Glioma-infiltrated voxels exhibited significantly smaller aperiodic exponents (flatter slopes) than normal-appearing cortex (β = –0.23, 95% CI: –0.27 to –0.20, p < 0.0001). In astrocytoma, the mean exponent was 3.00 in normal-appearing and 2.91 in glioma-infiltrated cortex. The glioma-related difference was significantly attenuated in astrocytoma (β = 0.15, 95% CI: 0.10 to 0.20, p < 0.0001). In GBM, the mean exponent was 2.99 in normal-appearing and 2.71 in glioma-infiltrated cortex. GBM showed the most pronounced difference between the cortex types (β = –0.06, 95% CI: –0.10 to –0.01, p = 0.0097). These findings demonstrate a consistent reduction in the aperiodic exponent across glioma-infiltrated regions, with the smallest exponent (i.e., the flattest slope) observed in glioblastoma. The parallel pattern of tissue-specific slope flattening across both the ECoG subdural array and MEG supports the reliability of the aperiodic exponent as a marker for glioma cortical infiltration. To assess the spatial distribution and localization of glioma-associated changes in aperiodic exponent values, we performed z-statistical mapping of MEG-derived exponent values at 30-50 Hz for each patient (**Extended Data Fig. 6A-C**). These z-maps revealed spatially localized reductions and gradients of the aperiodic exponent. Oligodendroglioma and astrocytoma exhibited smaller and spatially restricted deviations compared with glioblastoma. The z-score normalization approach controlled for patient-specific baseline variability and global spectra shifts, allowing for the identification of focal spectral abnormalities attributable to tumor presence. These findings demonstrate that MEG and ECoG-derived aperiodic exponent flattening is spatially localized to glioma-infiltrated cortex.

### Aperiodic Exponent Classifiers Accurately Predict Tumor Infiltration and Glioma Subtype

Given the discovery that adult-type diffuse glioma has a distinct aperiodic exponent, we next sought to determine the extent to which aperiodic exponent predicts tumor subtype. We trained a random forest classifier using the spectral exponent extracted at 70-150 Hz. Two classification tasks were tested: (1) distinguishing glioma-infiltrated from normal-appearing cortex at the electrode level; and (2) classifying glioma subtype from the aperiodic slope. In the first model, we used the combined cohort’s electrode-level aperiodic exponent values (n = 660) as the sole predictor to distinguish between glioma-infiltrated and normal-appearing cortex. A bootstrap random forest classifier was trained under a leave-one-patient-out cross-validation (LOPO-CV) framework to prevent overfitting and ensure generalizability. This classifier achieved an area under the receiver operating characteristic (ROC) curve (AUC) of 0.9388 with a misclassification rate of 14.39%, sensitivity of 62.7%, and specificity of 98.8% (**Fig. 2F, left**). Next, we determined whether the aperiodic exponent could be used to classify glioma subtype based solely on spectral physiology. Under LOPO-CV, the classifier demonstrated robust performance across subtypes, with AUC values of 0.8961 for oligodendroglioma, 0.8834 for astrocytoma, and 0.9179 for GBM, resulting in a misclassification rate of 29.09%. The classifier demonstrated high sensitivity for astrocytoma (85.4%) and glioblastoma multiforme (GBM, 80.1%), but lower sensitivity for oligodendroglioma (57.7%). Specificity was highest for Oligodendroglioma (99.6%), followed by glioblastoma (80.2%) and Astrocytoma (70.0%) (**Fig. 2F, right**).

We next trained random forest classifiers using aperiodic exponent values derived from MEG voxel data (n = 11,618 total voxels; n = 5,809 normal; n = 5,809 glioma-infiltrated) to test whether tumor infiltration and glioma subtype could be identified non-invasively. Using a LOPO-CV framework, the normal-appearing vs. glioma-infiltrated classifier achieved an AUC of 0.78, with a sensitivity of 72.7%, specificity of 68.6%, and a misclassification rate of 29.3% (**Extended Data Fig. 6D**). For glioma subtype classification, the MEG-based model achieved an AUC of 0.8119 for oligodendroglioma, 0.778 for astrocytoma, and 0.762 for GBM, with a misclassification rate of 38.2% (**Extended Data Fig. 6E**). The classifier exhibited high sensitivity for GBM (91.8%), but lower sensitivity for astrocytoma (27.9%) and oligodendroglioma (23.0%). Specificity was highest for oligodendroglioma (96.6%), followed by astrocytoma (85.6%) and glioblastoma (68.1%). These findings demonstrate that aperiodic spectral features derived from MEG carry the discriminative signal for both glioma presence and subtype, though performance is reduced relative to ECoG-based models. These findings demonstrate that the aperiodic exponent enables robust classification of glioma subtype based on the aperiodic exponent, suggesting that the electrophysiological state of the tumor-bearing cortex reflects its molecular features.

### Tumor Infiltration Correlates with Flatter Aperiodic Exponents

Glioma subtype accounts for broad differences in the aperiodic exponent; however, we hypothesize that regional variation in malignant cell infiltration may contribute to the flattening of the aperiodic slope. To test this, we quantified malignant cell burden in intraoperatively collected tissue samples from glioma-infiltrated cortex using SOX2 (for IDH-wildtype glioblastoma) or IDH1 R132H (for IDH-mutant gliomas) immunolabeling (**Fig. 2G**). Each sample was assigned the aperiodic exponent from its nearest electrode using a Euclidean distance-based mapping approach. Across samples, we observed a significant inverse correlation between tumor burden and aperiodic exponent (slope = –0.029, 95% CI: –0.047 to –0.012, p = 0.004, R² = 0.578; **Fig. 2H**), indicating that regions with higher malignant cell infiltration exhibit greater spectral flattening. This relationship was consistent across glioma subtypes, suggesting that the aperiodic exponent reflects not only tumor subtype but also the extent of malignant cell cortical infiltration.

### Aperiodic Exponent Tracks Trial Accuracy in Glioma-infiltrated Cortex During Language Tasks

Although the aperiodic exponent may offer insights into tumor subtype, its connection to cognition and behavior remains poorly understood. Connections have been drawn between aperiodic exponent dynamic changes and performance in various cognitive contexts ^47–50^. We therefore next sought to understand the extent to which gliomas-induced neuronal excitability disrupts domain-specific cognitive processes such as language or executive function^51–54^. Trial-by-trial interrogation of the aperiodic exponent may offer a dynamic reflection of the influence of glioma infiltrations on cognitive engagement.

To evaluate whether behavioral performance modulates aperiodic dynamics across glioma-infiltrated and normal-appearing cortex, patients underwent two semantic naming assessments: picture naming (PN) and auditory naming (AN). Both tasks involved structured trials (PN=48 trials; AN=32 trials) with stimulus presentation followed by a predefined response window (**Fig. 3A**). Accuracy was determined on a per-response basis, enabling stratification of neural data by performance outcome at the patient level^55^. To contextualize task-related changes, resting-state epochs were sampled from the same patients and matched in duration and number to the respective language task trial. Trial-matched rest segments served as a within-subject baseline for comparing condition-specific shifts in the aperiodic exponent. During picture naming, glioma-infiltrated cortex exhibited a significant flattening of the aperiodic exponent during incorrect trials compared to correct trials (**Fig. 3B, right**). In the normal-appearing cortex, this effect was not as evident, and the exponent did not substantially influence accuracy. (**Fig. 3B, left**). Using linear mixed-effects models, we modeled the aperiodic exponent as a function of resting-state exponent, trial correctness, and cortex tissue type. In the simplest model (containing resting-state exponent and accuracy as predictors), task-related aperiodic exponents were significantly predicted by resting-state values (β = 0.56, 95% CI: 0.44 to 0.69, p < 0.0001), and incorrect trials were associated with flatter exponents (β = –0.11, 95% CI: –0.22 to –0.0003, p = 0.0438). Adding tissue type improved model fit (p < 0.001), revealing that glioma-infiltrated cortex exhibited significantly flatter task-related exponents compared to normal-appearing cortex (β = –0.59, 95% CI: –0.82 to –0.35, p < 0.0001). The data were then stratified based on glioma subtype, which further improved the model fit (p = 0.0016); task-related exponents were flatter in glioblastoma (β = –0.40, 95% CI: –0.54 to –0.25, p < 0.001) compared to astrocytoma and oligodendroglioma. An interaction model revealed that incorrect trials in glioma-infiltrated cortex showed the greatest flattening (interaction β = – 0.82, 95% CI: –1.19 to –0.46, p < 0.0001). A similar pattern was identified during auditory naming: incorrect trials were associated with significantly flatter exponents in glioma-infiltrated regions but not in normal-appearing cortex (**Fig. 3C**).

**Fig. 3.**
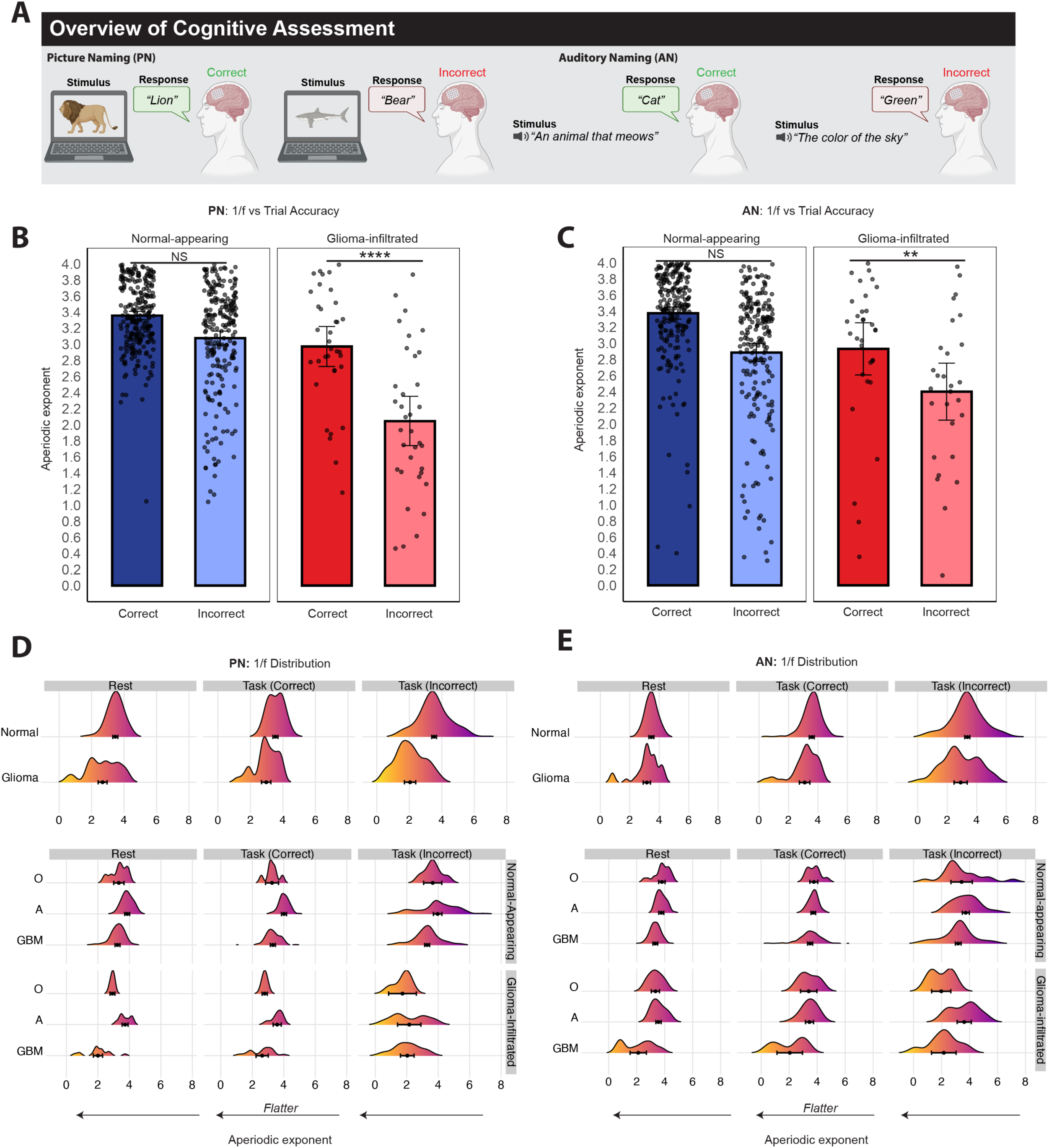
Relationship between aperiodic exponent and cognitive performance during picture naming (PN) and auditory naming (AN) language tasks. **A,** Overview of cognitive assessment language tasks and the difference between correct and incorrect trials. Left: PN task, where participants name the object shown on a laptop screen (e.g. “lion”) at stimulus onset. Periods of interest include the baseline period (pre-stimulus) and the post-stimulus onset window (500 ms). Right: AN task, where participants name an object (e.g., “Cat”) following a sentence prompt (“What animal meows?”). Periods of interest include the baseline period (pre-sentence) and the post-stimulus window (500 ms). **B,** Average aperiodic exponent extracted at 500 ms post-stimulus windows during PN trials at 70-150 Hz for normal-appearing (blue) and glioma-infiltrated (red) electrodes comparing correct (dark) and incorrect (light) responses. Linear mixed effects modeling showcased glioma-infiltrated electrodes having significantly lower aperiodic exponents for incorrect responses compared to correct responses (****p≤0.0001), while normal-appearing electrodes showed no significant differences across trial response (ns, p=0.575). **C,** Average aperiodic exponent during AN trials, comparing correct and incorrect responses across glioma-infiltrated and normal-appearing electrodes. Linear mixed effects modeling yielded similar results as PN where the aperiodic exponent was significantly lower in incorrect trials compared to correct trials within glioma-infiltrated electrodes (**p≤0.001). There was no significant difference in aperiodic exponent between trial response within normal-appearing electrodes (ns, p=0.480). Error bars represent the 95% confidence interval. **D,** Ridgeline density plots showing the distribution of the aperiodic exponent values during resting state, correct trials, and incorrect trials across glioma-infiltrated and normal-appearing electrodes during the PN language task. Distributions of the aperiodic exponent are further stratified by glioma subtype (O, A, GBM). Glioma-infiltrated electrodes consistently exhibit a shift toward flatter exponents during task-related errors compared to resting state and correct trials indicating a change in the evoked response to stimulation. **E,** Density plots showing the distribution of the aperiodic exponent values during resting state, correct trials, and incorrect trials within the AN language task. Similar to PN, when stratified by subtype, glioma-infiltrated regions show shifts toward shallower exponents, particularly during task-related errors. This shift is most prominent in GBM. Dot represents the mean, error bars represent the 95% confidence interval (*p≤0.01). Abbreviations: PN = picture-naming, AN = auditory naming, ns = no significance, O = oligodendroglioma, A = astrocytoma, GBM = glioblastoma multiforme

Picture naming reflects semantic language processing; however, we performed the same analysis during auditory naming to understand the generalizability of the aperiodic slope during speech processing. Task-specific aperiodic exponents again shifted positive relative to resting-state exponent (β = 0.67, 95% CI: 0.52 to 0.82, p < 0.0001), and incorrect responses were associated with flatter exponents compared to correct trials (β = –0.19, 95% CI: –0.33 to –0.04, p = 0.012). Stratifying glioma-infiltrated versus normal-appearing electrodes into the model revealed that glioma-infiltrated cortex exhibited significantly flatter task exponents relative to normal-appearing cortex (β = –0.28, 95% CI: –0.51 to –0.04, p = 0.026). Model fit was further improved by including glioma subtype (p = 0.013), with glioblastoma (β = –0.23, 95% CI: –0.42 to –0.04, p = 0.017) demonstrating flatter task-related exponents compared to astrocytoma and oligodendroglioma. These effects persisted when restricting the analysis to correct trials (resting-state β = 0.76, 95% CI: 0.62 to 0.90, p < 0.0001; tissue-type β = –0.23, 95% CI: –0.45 to –0.02, p = 0.035), and trends in the same direction were observed during incorrect trials (resting-state β = 0.50, 95% CI: 0.23 to 0.76, p < 0.001; tissue-type β = –0.37, 95% CI: –0.79 to 0.05, p = 0.085). Together, these findings suggest that glioma infiltration and subtype influence the dynamic modulation of cortical excitability during both picture and auditory naming. Ridgeline plots displaying kernel density estimates of exponent values across resting-state, correct, and incorrect trials confirmed the trends identified in Fig. 3B and C. Infiltrated cortex consistently showed leftward shifts (towards flatter values) in the exponent distribution of incorrect trials relative to baseline and correct task performance (**Fig. 3D-E, top)**. When stratified by glioma subtype, task-induced flattening of the exponent was evident across oligodendroglioma, astrocytoma, and glioblastoma, but was most exaggerated in glioblastoma-infiltrated cortex (**Fig. 3D-E, bottom**). Tumor-specific flattening of aperiodic exponent during incorrect trials suggests increased tumor infiltration may amplify circuit-level dysregulation during task failure.

### Transcriptional Architecture of Normal-Appearing Cortex Reflects Local Aperiodic Exponent and Excitatory-Inhibitory Balance of Neuronal Circuits

Dynamic transcriptomic programs are essential for cognitive processing^56^, yet the molecular correlates of aperiodic exponent, including neuronal expression programs of excitation and inhibition dominance, remain unknown. We performed snRNA-seq from cortical regions annotated for aperiodic exponent to identify the molecular features within normal-appearing and glioma-infiltrated cortex. Subdural high-density arrays were used to compute the aperiodic exponent at each electrode during resting-state activity. Using Euclidean distance, spatially co-localized tissue samples were aligned between electrode coordinates and sample centroids. Patient derived tissue samples from WHO grade 2 oligodendroglioma, WHO grade 2 astrocytoma, and grade 4 glioblastoma reflecting the highest and lowest aperiodic exponent values were selected for sequencing, ensuring within-patient comparison of steep and flat aperiodic exponent regions (n=14 cortex samples from 7 patients; **Extended Data Fig. 7**). We sequenced a total of 508,899 nuclei and performed dimensionality reduction and cell-type classification based on canonical expression programs malignant and non-malignant cell populations including neuronal subpopulations (**Fig. 4A, Extended Data Fig. 8**). Uniform manifold approximation and projection (UMAP) projection revealed discrete clustering of key excitatory (glutamatergic) and inhibitory (GABAergic) neuronal populations (**Fig. 4B**). Assignment of transcriptionally defined neuron subtypes was consistent across normal-appearing and glioma-infiltrated cortex. Copy number variation (CNV) analysis verified chromosomal alterations consistent with glioma infiltration in all tumor-infiltrated samples (n=12), and genomic stability in the normal-appearing cortex samples (n=2), validating the separation of neoplastic and non-neoplastic tissue (**Extended Data Fig.7B-F**). Within the neuronal population, hierarchical classification of cells into excitatory glutamatergic and inhibitory GABAergic neuron populations was performed. Both glutamatergic and GABAergic neurons were further divided into nine main subtypes based on their biological significance. The distributions of neuronal subtypes did not differ between steep (high exponent; n = 7) and flat 1/f samples (low exponent; n = 7), indicating that wide neuron diversity was preserved in both cohorts (**Fig. 4C**). Neuronal subtypes exhibited distinct morphological, electrophysiological, and transcriptomic properties. Similar distributions of neuronal populations between steep and flat samples allowed us to attribute subsequent analysis to transcriptional expression programs rather than the abundance of neuronal subtypes. To define molecular drivers of excitability in normal-appearing cortex, we calculated an excitatory score for each neuronal nucleus based on the aggregate expression of 42 canonical glutamatergic genes for their role in neuronal depolarizations and hyperexcitability^57–60^. Neurons within the flat aperiodic exponent sample exhibited significantly higher excitatory module scores compared with those from the steep aperiodic exponent samples (t(8954) = 70.73, p < 0.0001, Mann-Whitney U test; **Fig. 4D)**. The mean excitatory score was 0.2345 in flat aperiodic exponent regions and 0.1360 in steep aperiodic exponent regions, with a mean difference of 0.09855 ± 0.001393 (95% CI: 0.096 to 0.101; **Fig. 4D**). This difference was consistent across most neuron subpopulations in flat aperiodic exponent samples (**Fig. 4E**), indicating that variation in the aperiodic exponent reflects both global and subtype-specific differences in glutamatergic gene expression. The largest shifts were observed in lamina 2/3 and lamina 6 intratelencephalic (IT) neurons–subtypes involved in intracortical and long-range projection^61^, which may be associated with coordinated reductions in both local and output excitatory signaling in cortical regions characterized by flatter aperiodic exponent values. Next, we performed differential gene expression analysis between neurons from steep and flat aperiodic exponent samples to identify molecular drivers of aperiodic exponent. Flat aperiodic exponent regions were enriched for genes involved in excitatory synaptic function, including glutamate receptor subunits (*GRIN2A, GRM7*), vesicular glutamate transporters (*SLC17A7*), and synaptic scaffolding proteins (*HOMER1*). In contrast, steep exponent regions displayed increased expression of genes associated with inhibitory tone (*GABBR2, SL6A12*) (**Fig. 4F**). Gene ontology enrichment revealed that steep exponent cortex was enriched for processes related to actin organization, RNA splicing, and supramolecular fiber organization (**Fig. 4G**), while flat exponent cortex was enriched in processes involving regulation of the membrane potential, synapse organization, potassium ion transport and glutamatergic synaptic transmission (**Fig. 4H**). These findings support the interpretation that a flatter aperiodic exponent in normal-appearing cortex reflects a transcriptional state of excitation dominance when compared to a steep 1/f slope. We validated these findings at the protein level by performing immunofluorescence on formalin-fixed, paraffin-embedded patient-derived tissue sections reflecting flat and steep exponent (n = 2 samples from 1 patient). Consistent with transcriptomic analyses, flat-exponent normal-appearing cortex contained a significantly higher proportion of Homer1⁺/NeuN⁺ cells compared to steep-exponent samples (t(22) = 4.040, p = 0.0005, unpaired t-test). The mean proportion of Homer1⁺/NeuN⁺ cells was 49.09% in flat-exponent tissue and 19.72% in steep-exponent tissue, yielding a mean difference of 29.37 ± 7.27 (95% CI: 14.30 to 44.45; **Fig. 4I–J**). Detection of the glioma stem cell marker SOX2 was absent from both steep and flat exponent normal-cortex samples, confirming the lack of tumor infiltration (**Fig. 4I**). Together, these results demonstrate that even within structurally intact, non-neoplastic cortex, the aperiodic exponent reflects excitatory tone. Flat exponent regions are characterized by elevated glutamatergic gene expression and synaptic signaling pathways, whereas steep exponent regions exhibit a shift toward less excitation.

**Figure 4.**
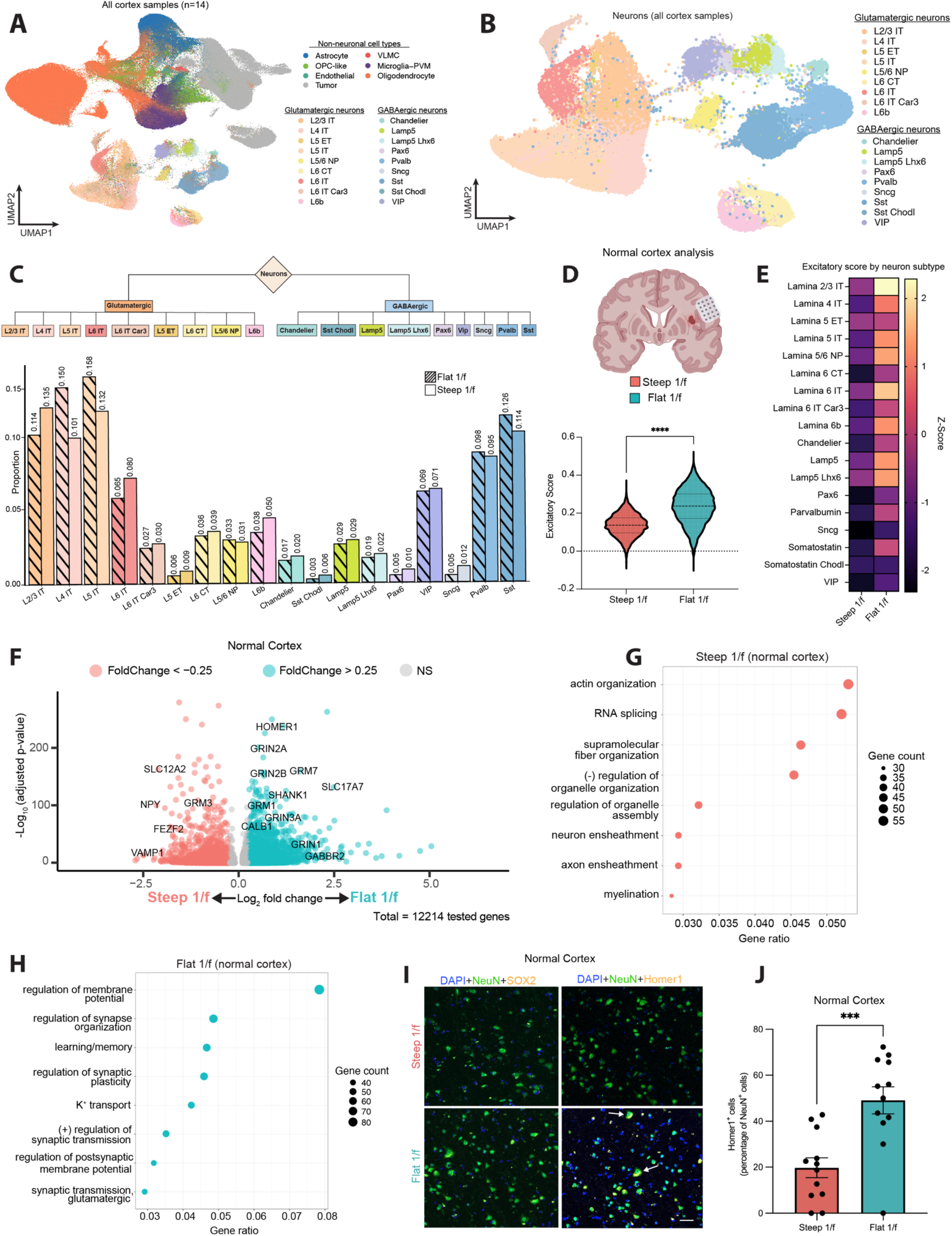
Transcriptional analysis reveal excitation dominant tone in flat exponent region neurons of normal-appearing cortex. **A**, Uniform manifold approximation and projection (UMAP) of all nuclei from both glioma-infiltrated and normal-appearing cortex tissue, annotated by broad cell type categories, including neurons (purple), glial cells, and tumor-infiltrating populations. **B**, UMAP embedding of neuronal nuclei, colored by transcriptionally defined neuronal subclasses, illustrating the diversity of excitatory (glutamatergic) and inhibitory (GABAergic) neurons. Extended data figure 8 displays feature plots for each UMAP. **C**, Hierarchical classification of neuronal subtypes into excitatory (Glutamatergic, left) and inhibitory (GABAergic, right) classes. Neuronal subclass composition differs between cortical regions preselected for steep and flat exponent samples. Bar plots display relative proportions of neuronal subclasses in cortical regions with steep and flat exponent assignments. **D**, Top: Coronal gross anatomical brain slice illustration depicting an insular tumor and the overlying electrode grid positioned on adjacent normal-appearing cortex. Subsequent panels reflect single-nucleus RNA sequencing (snRNA-seq) data derived from steep and flat exponent samples collected from normal-appearing cortex. Bottom: violin plot showing a significant difference in excitatory marker expression between steep and flat exponent samples (p < 1 × 10⁻¹⁰, two-tailed Mann Whitney U Test), suggesting an association between excitation-dominant transcriptional signatures and flatter exponents. **E,** Heatmap illustrating the z-score normalized difference in excitatory module scores between steep and flat exponent normal-appearing cortex samples (x-axis), displayed across identified neuronal subtypes (y-axis). Higher z-scores indicate relative enrichment within that cell type. **F**, Volcano plot showing log₂ fold-change of gene expression between steep and flat exponent normal-appearing cortex samples (two-sided Wilcoxon rank-sum test). Glutamatergic receptor subunits (e.g. *GRIN2A, GRM7, GRIA2*), inhibitory synaptic genes (e.g. *GABBR2*), and synaptic plasticity regulators (e.g. *HOMER1*) are significantly differentially expressed. **G-H,** Gene ontology analysis of biological processes enriched in steep (G) versus flat exponent normal-appearing cortex (H). Top enriched pathways in each condition are displayed, with point size indicating the gene count. **I,** Representative immunofluorescence images of formalin-fixed and paraffin embedded normal-appearing cortex samples showing DAPI+ nuclei and NeuN+ mature neurons. Left: SOX2 was not detected across both steep and flat exponent samples, consistent with the absence of glioma stem-like or infiltrating tumor cells in this region. Right: Images showing the distribution of Homer1+ and NeuN+ positive cells between steep and flat exponent samples. Scale bar, 50 μm. **J,** Quantification of Homer1+ and NeuN+ cells as a percentage of NeuN+ cells. Bars represent mean ± SEM, and statistical comparisons were performed using two-tailed T-test (p < 0.001).

### Transcriptional Architecture of Glioma-Infiltrated Cortex Reflects Local Aperiodic Exponent

Building on the observation that the aperiodic exponent in normal-appearing cortex reflects local excitatory neuronal tone, we next assessed whether the aperiodic exponent within glioma-infiltrated cortex similarly corresponds to transcriptionally defined excitatory state. Single-nucleus RNA sequencing was performed on 12 spatially localized glioma-infiltrated cortical samples from six patients based on aperiodic slope computed from subdural array data. Steep (total n=6) and flat exponent samples (total n=6) were selected from cortical tissues acquired from each patient in Figure 2A based on the highest and lowest slope values. The sample cohort included two patient samples with oligodendroglioma, astrocytoma, and glioblastoma, allowing comparisons across glioma subtypes used for electrophysiology analysis (**Extended Data Fig. 7A**). Differential gene expression analysis revealed widespread transcriptional divergence between steep and flat exponent glioma-infiltrated cortex (**Fig. 5A, right**). Flat exponent samples showed increased expression of excitatory synapse-related genes, including *HOMER1*, *GRIN3A*, *SNAP25*, *GRM1*, and *GRM8,* while steep exponent samples exhibited upregulation of *GABRA3*, *GABRA5*, *SLC12A2*, and *SHANK1*–genes implicated in inhibitory signaling, calcium buffering, and stress responses ^62–65^. Gene ontology analysis supported this dissociation: steep exponent samples were enriched for pathways related to ribosome biogenesis, oxidative phosphorylation, and apoptotic signaling (**Fig. 5B**), whereas flat exponent samples were enrichment in synaptic assembly, membrane potential regulation, and chloride transport (**Fig. 5C**). Gene set enrichment analysis using curated validated glutamatergic gene modules revealed greater enrichment in flat versus steep exponent samples (NES = 2.54, adjusted p = 4.42×10^−5^; **Fig. 5D**). Pathway enrichment focused on neuronal biophysical processes further highlighted increased representation of glutamatergic signaling, neurotransmitter receptor activity, and postsynaptic potential regulation in flat exponent regions, whereas steep regions were enriched in developmental processes (**Fig. 5E**). To test whether this pattern of excitation was conserved across glioma subtypes, we stratified excitatory module scores by tumor subtype and compared flat versus steep aperiodic exponent samples (**Fig. 5F**). In oligodendroglioma, astrocytoma, and glioblastoma, flat exponent cortex exhibited higher excitatory scores compared with steep exponent glioma cortical samples. In oligodendroglioma, the mean excitatory score was 0.1414 in flat cortex versus 0.1185 in steep cortex (difference = 0.0229 ± 0.0010, t(10128) = 23.16, p < 0.0001, 95% CI: 0.0210 to 0.0249], Mann-Whitney U test). In astrocytoma, the difference was smaller but still significant (flat: 0.1425 vs. steep: 0.1390; difference = 0.0035 ± 0.0007, t(31417) = 5.28, p < 0.0001, 95% CI: 0.0022 to 0.0048, Mann-Whitney U Test). Glioblastoma exhibited the most significant difference, with flat cortex showing a mean excitatory score of 0.1848 versus 0.1005 in steep cortex (difference = 0.0844 ± 0.0015, t(8446) = 56.59, p < 0.0001, 95% CI: 0.0814 to 0.0873, Mann-Whitney U test). Z-scored excitatory scores across glutamatergic and GABAergic neuronal subtypes showed a broad upregulation of excitation in flat exponent samples in all excitatory neuronal populations, with no single subtype solely responsible (**Fig. 5G**). This effect was again most prominent in glioblastoma compared to astrocytoma and oligodendroglioma. We next sought to validate our differential gene expression analysis using patient-derived electrophysiologically annotated samples. We performed immunofluorescence staining and confocal microscopy for Homer1 and NeuN on FFPE patient- and sample-matched glioma-infiltrated cortex tissue sections (**Fig. 5H**). Homer1 was selected for validation because it was among the most consistently upregulated excitatory synaptic genes in flat exponent regions across all three glioma subtypes (**Extended Data Fig. 9**). Quantification revealed a greater proportion of Homer1⁺/NeuN⁺ excitatory neurons in flat compared to steep exponent regions (mean = 42.01% in flat vs. 30.09% in steep, difference = 11.92 ± 5.05, 95% CI = 1.86 to 21.98, p = 0.0209, unpaired t-test; **Fig. 5I, left**). When stratified by glioma subtype a trend towards significance was identified in all samples however, differences in Homer1 labeling was significant in glioblastoma, where flat exponent regions showed a higher proportion of Homer1⁺/NeuN⁺ excitatory neurons compared to steep regions (mean = 53.00% in flat vs. 27.98% in steep; mean difference = –25.02 ± 7.95, 95% CI: –45.65 to –4.40, p = 0.0149, Dunnett’s T3 post hoc following Brown-Forsythe and Welch ANOVA). In contrast, no significant difference was observed in oligodendroglioma (mean = 42.50% in flat vs. 35.53% in steep; mean difference = –6.97 ± 8.15, 95% CI: –27.94 to 14.00, p = 0.7768) or astrocytoma (mean = 30.51% in flat vs. 26.76% in steep; mean difference = –3.76 ± 9.29, 95% CI: –27.75 to 20.23, p = 0.9682; **Fig. 5I, right**). These findings suggest that excitatory remodeling associated with flatter aperiodic exponents may be most pronounced in glioblastoma-infiltrated cortex.

**Figure 5.**
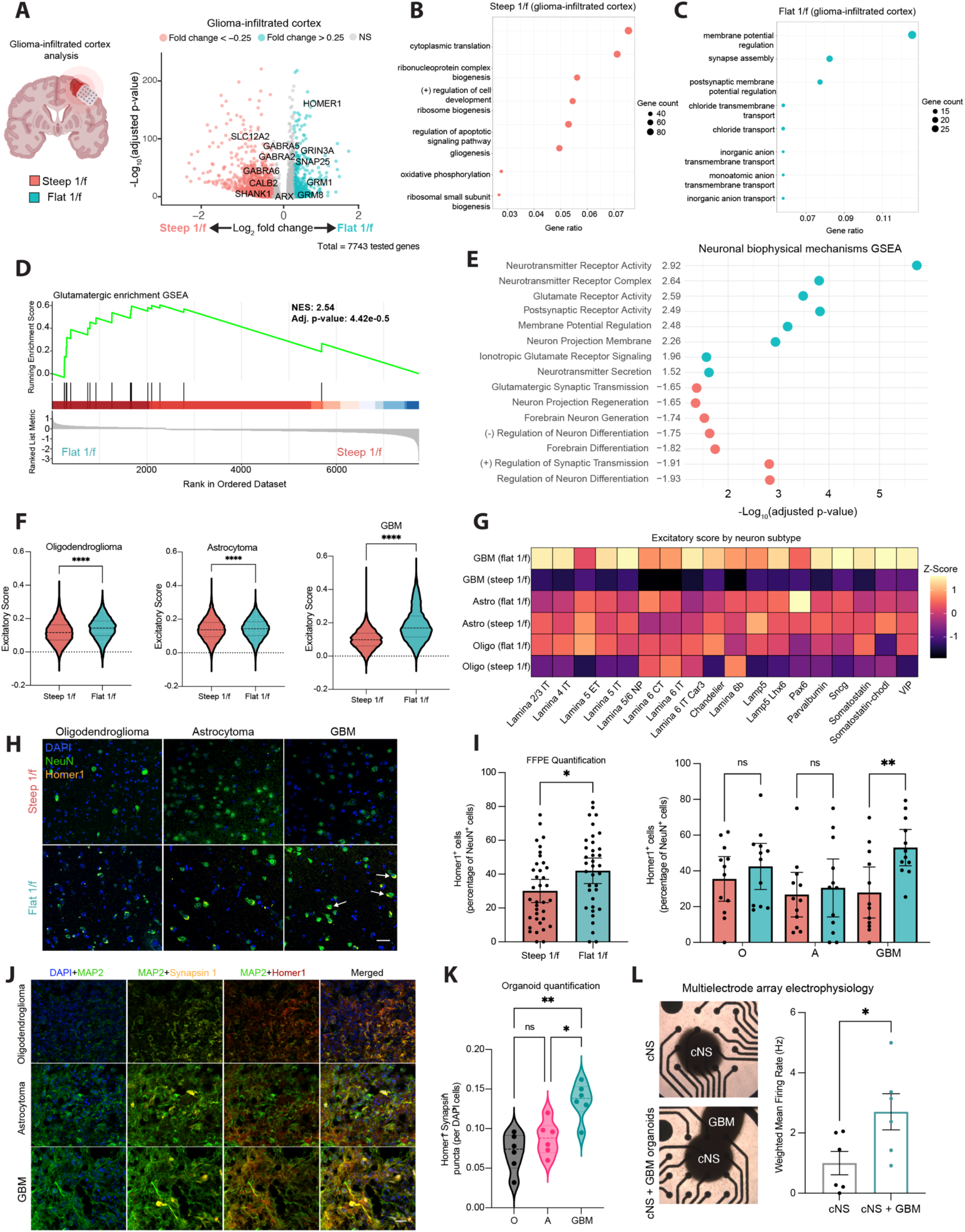
Transcriptional analysis reveals excitation dominance in flat exponent cortical region neurons within glioma-infiltrated cortex. **A,** Left: Coronal gross anatomical brain slice illustration depicting a cortically-projecting tumor and the overlying electrode grid positioned on adjacent glioma-infiltrated cortex. This panel provides the anatomical context for subsequent single-nucleus RNA sequencing (snRNA-seq) analyses in this figure. Right: volcano plot showing log₂ fold-change of gene expression between steep and flat exponent glioma-infiltrated cortex samples (two-sided Wilcoxon rank-sum test). **B-C,** Gene ontology analysis of biological processes enriched in steep (B) versus flat exponent glioma-infiltrated cortex (C). Top enriched pathways in each condition are displayed, with point size indicating the gene count. **D,** Gene set enrichment analysis (GSEA) of glioma-infiltrated cortex steep (n=6 samples) and flat exponent samples (n=6 samples), showing enrichment of glutamatergic synaptic pathways in flat versus steep exponent glioma-infiltrated cortex samples. Normalized enrichment scores (NES) and adjusted P values (false discovery rate correction) are indicated. **E,** Summary of neuronal biophysical mechanisms gene set enrichment analysis associated with steep versus flat exponent samples, highlighting key glutamatergic and inhibitory pathways. **F,** Violin plots of excitatory module scores in oligodendroglioma, astrocytoma, and glioblastoma, stratified by steep versus flat exponent samples. Center values represent median, dashed lines represent upper and lower quartile, and statistical significance was determined using a Mann-Whitney U test, ****p < 0.0001. **G,** Heatmap illustrating the z-score normalized difference in excitatory module scores between steep and flat exponent glioma-infiltrated cortex samples by glioma subtype (x-axis), displayed across identified neuronal subtypes (y-axis). Higher z-scores indicate relative enrichment within that cell type. H, Representative immunofluorescence images of formalin fixed and paraffin embedded glioma-infiltrated cortex samples showing the distribution of Homer1+ and NeuN+ positive cells between steep and flat exponent samples. Scale bar, 50 μm. **I,** Quantification of Homer1+ and NeuN+ cells as a percentage of NeuN+ cells of all samples combined by status (steep vs flat; left) and stratified by glioma subtype (right). Bars represent mean ± SEM, individual dots represent a snapshot used for quantification, and statistical comparisons were performed using two-tailed Mann-Whitney U test (*p < 0.05, **p < 0.001). **J,** Representative confocal microscopy images of organoid slices stained for MAP2 (neuronal marker), synapsin-1 (presynaptic marker), and Homer1 (postsynaptic marker) across glioma subtypes. Scale bar, 20 μm. **K,** Quantification of glioma-infiltrated organoids. Colocalization of Homer1 and Synapsin-1 is quantified per DAPI+ nucleus across glioma subtypes. Bars represent mean ± SEM, individual dots represent a snapshot used for quantification, and statistical comparisons were performed using one-way ANOVA (*p < 0.05, **p < 0.001). **L,** Multielectrode array analysis of glioma-infiltrated GBM organoids co-cultured with neurons, showing weighted mean firing rate (Hz) across conditions. Bars represent mean ± SEM. Statistical significance was determined using a two-tailed Mann-Whitney U test (*p < 0.05).

The excitation dominance of gene expression programs of flat exponent glioma-infiltrated cortex, while most striking within glioblastoma-infiltrated cortex, was less evident in IDH mutant oligodendrogliomas and astrocytoma samples. To validate this finding and investigate the electrophysiological and functional interactions between glioma cells and neurons, we developed a 3-dimensional fusion model comprising patient-derived oligodendroglioma, astrocytoma, or glioblastoma organoids in co-culture with mouse cortical neurospheres (cNS)^12,66,67^. Organoids were derived from 7 patients with WHO grade 2 or grade 3 oligodendroglioma or grade 2 or grade 3, or glioblastoma to enable comparisons across histologic grades. Multielectrode array (MEA) recordings of the central nervous system (cNS) with or without fusion to glioma organoids were used to assess the effect of glioma infiltration on hyperexcitability. Organoids co-cultured with glioblastoma exhibited increased Homer1⁺/Synapsin1⁺ puncta colocalized with MAP2⁺ neurons compared with those exposed to astrocytoma or oligodendroglioma cells (**Fig. 5J**). Quantification revealed that GBM-exposed organoids had a significantly higher density of Homer1⁺/Synapsin1⁺ puncta per DAPI-labeled cell (mean = 0.1355) compared to organoids exposed to oligodendroglioma (mean = 0.0703; mean difference = –0.0652 ± 0.0134, 95% CI: –0.1031 to – 0.0272, p = 0.0019) or astrocytoma (mean = 0.0878; mean difference = –0.0477 ± 0.0128, 95% CI: –0.0838 to –0.0116, p = 0.0112), based on Dunnett’s T3 post hoc test following Brown-Forsythe and Welch ANOVA (**Fig. 5K)**. No significant difference was observed between organoids exposed to oligodendroglioma versus astrocytoma cells (mean difference = –0.0175 ± 0.0130, 95% CI: –0.0542 to 0.0192, p = 0.4770). Next, using multielectrode arrays, we determined neuronal hyperexcitability by the weighted mean firing rate from co-organoid cultures. Organoids co-cultured with patient-derived glioblastoma cells exhibited higher weighted mean firing rates (WMFR; Hz) compared to control organoids (mean = 2.704 in cNS+ glioblastoma vs. 1.000 in cNS; mean difference = 1.704 ± 0.7145, 95% CI: 0.1124 to 3.296, p = 0.0383, unpaired t-test; **Fig. 5L**). These findings indicate that glioma exposure is sufficient to enhance spontaneous neuronal network activity in vitro. The relationship between flat exponent and excitatory state in glioma-infiltrated cortex was consistent across gene expression, protein labeling, and network activity, suggesting that local aperiodic exponent reflects excitability dominance within normal and glioma-infiltrated cortex, with greatest significance in glioblastoma-infiltrated cortex.

## DISCUSSION

Neural circuits, the fundamental units of computation in the brain, are crucial for glioblastoma progression and are therefore essential to understanding glioma behavior and treatment response. Few measures exist to interrogate the dominance of excitation or inhibition in neuronal populations resulting from malignant infiltration into the human brain. Here, we integrate electrophysiological, genomic, and behavioral analyses of normal-appearing and glioma-infiltrated samples, as well as patient-derived co-cultures, to demonstrate that the aperiodic exponent derived from the power spectral density reflects glioma-induced excitation-inhibition states both globally and regionally. We find that the glioma-infiltrated cortex displays a comparatively flat aperiodic exponent, reflecting a shift toward an excitation-dominant state, which is glioma subtype and patient-specific. This finding aligns with previous evidence suggesting significant shifts toward an excitation-dominant state^12,17,20^, and confirms that gliomas infiltration remodels the spectral profile of normal cortex. The distinct electrophysiological signature of glioma infiltration emerges not only during resting-state activity but is also associated with cognitive performance, with aperiodic exponent correlating with semantic naming accuracy across multiple language tasks^53^. This is consistent with prior observations that aperiodic spectral parameters vary systematically across brain structures, reflecting both cytoarchitectural properties and disease severity^68^. By connecting behavioral analysis with molecular features, we observe that cortical regions characterized by flatter aperiodic exponents, across both normal-appearing and glioma-infiltrated cortex, exhibit elevated neuronal excitatory expression programs and transcriptional signatures consistent with heightened excitatory signaling.

Employing a multi-modal approach that integrates electrophysiology, behavior, and single-nucleus transcriptomics, we establish the aperiodic exponent as a vital metric for delineating glioma-driven neural dysregulation. This comprehensive framework elevates our understanding of the biological underpinnings of tumor-induced remodeling of neuronal circuits. Importantly, our findings align with recent investigations using ECoG to differentiate tumor-affected cortex from peritumoral regions by analyzing spectral properties. This consistency strengthens the reliability of the aperiodic exponent as a malignancy biomarker. Our study extends prior work by demonstrating, for the first time, that electrodes overlying glioma-infiltrated cortex with flattened aperiodic exponents correspond to regions enriched for transcriptionally defined excitatory neuronal populations, characterized by significantly higher excitatory module scores, suggesting a link between aperiodic flattening and hyperexcitable states. Thus, the relationship between the aperiodic exponent and E/I imbalance within glioma pathology becomes compellingly evident ^69–71^; yet, it has not been fully explored in the human brain. Observing stronger effects in the high-frequency range, where dynamics are more likely to follow a simple power law structure, suggests that this is indeed an aperiodic effect and not a narrowband artifact ^72^. Furthermore, the aperiodic exponent’s dynamic state, reflected by its sensitivity to pharmacologically induced GABA_A_ agonist E/I shifts, emphasizes its potential as a physiological marker of cortical excitability in older adult populations with glioma, as demonstrated by its steepening with increasing propofol infusion dose. Such findings extend prior work derived from computational modeling, EEG, and nonhuman primate electrophysiology, underscoring the conserved nature of this spectral feature across species and cortical states^23,39^. Stratification of the aperiodic exponent according to molecularly defined glioma subtypes reveals a significant gradient, where glioblastomas exhibit the flattest exponents, deemed indicative of a heightened E/I imbalance proportional to the malignant cell burden, consistent with tumor-induced hyperexcitability seen in malignant gliomas ^21,73,74^. This finding suggests that tumor grade affects the degree of E/I disruption. Our findings are consistent with MEG data, further providing cross-modal validation of these results, albeit acknowledging the limitations inherent in magnetoencephalography at higher frequencies.^75,76^ The pronounced flattening of the aperiodic exponent within glioma-affected regions suggests extensive repercussions on neural circuit function, potentially contributing to the cognitive impairments often observed in patients with gliomas^77–79^.

An additional compelling aspect of our analysis is the capacity of electrophysiological glioma subtype classifiers to successfully differentiate between glioma-infiltrated and normal-appearing cortical regions, as well as classify glioma subtypes based solely on the aperiodic exponent, similar to studies that have used electrophysiological or genomic data for classification^53,80–83^. Additionally, we show that tumor infiltration correlates quantitatively with the aperiodic exponent, establishing a direct relationship between histological burden and electrophysiological remodeling. This provides histological validation of the circuit-level alterations measured electrophysiologically and extends prior work demonstrating reduced aperiodic exponents in glioma-infiltrated cortex^69^. By quantifying tumor cell density and spatially matching histological samples to subdural electrodes, we link cellular-level tumor invasion to macroscale aperiodic dynamics. This relationship was consistent across glioma subtypes, suggesting a shared electrophysiological signature of infiltration.

Our findings during semantic naming substantiate the known relationship between E/I imbalance and cognitive task performance. Glioma-induced hyperexcitability adversely affects cognitive processing. Further supporting the role of the aperiodic exponent in cognition, recent work has demonstrated that variations in aperiodic slope during resting-state recordings from EEG were associated with broad cognitive abilities in a healthy cohort^48^. The authors’ findings report that participants who performed better across multiple cognitive tasks had steeper aperiodic exponents. Other recent studies have found that flatter aperiodic exponents vary with task engagement and during higher cognitive demand^47,84^. This aligns with our findings that glioma-driven flattening of the aperiodic exponent, reflecting an excitation-dominant state, is associated with semantic naming errors. These data suggest a shared mechanistic link between neural excitability and cognition. The trial-by-trial modulation of the aperiodic exponent provides a dynamic reflection of the influence of glioma infiltrations on maintaining the E/I balance during cognitive engagement. This contribution merits further exploration to understand whether the findings in this study are persistent across cognitive domains, such as language or executive function, ^51,52^, and to identify the cellular and molecular drivers of such impairments. Additionally, it is essential to determine whether function may be restored through the restoration of E/I towards an established norm.

This study’s DNA and snRNA sequencing approach elucidates the molecular underpinnings of aperiodic slope. In normal-appearing cortex, flatter aperiodic exponents correspond to greater expression of an excitatory genomic signature. In glioma-infiltrated areas, the transcriptional landscape highlights hyperexcitability, characterized by the upregulation of genes associated with excitatory synapses and the inhibition of those involved in synaptic assembly and membrane potential regulation. This balanced interplay of transcriptional activity suggests both compensatory mechanisms and possibly dysfunction within the neuronal microenvironment, warranting deeper investigation into the specific molecular pathways involved, particularly those governing synaptic signaling, ion homeostasis, and glioma-neuron interactions ^85–87^. Our findings highlight the importance of excitatory neuron-enriched genes such as *HOMER1, SNAP25*, and *SLC17A7*, as well as pathways involving glutamatergic transmission and potassium-chloride transport, implicating these as key regulators of local E/I balance within glioma-infiltrated cortex.

### LIMITATIONS

Our study has several significant limitations. First, the spatial sampling of glioma specimens and their associated electrophysiological recordings was dictated by clinical necessity, resulting in heterogeneous electrode coverage across patients and potential under-sampling of functionally relevant peritumoral regions. Anatomical classification of electrodes as “normal-appearing” or “glioma-infiltrated” was based on magnetic resonance imaging and visual inspection, which may fail to detect microscopic invasion or functional disruption. The snRNA-seq analysis was limited to tissue samples from the extremes of the aperiodic exponent distribution within each selected patient. It may not capture the full transcriptomic gradient across the E/I spectrum. Finally, the cross-sectional nature of this dataset limits the ability to make causal inferences. Longitudinal, mechanistic, and interventional studies are needed to validate the aperiodic exponent further as a reliable biomarker of network dysregulation in glioma and to determine its potential utility in clinical decision making.

## CONCLUSION

This work highlights the significance of the aperiodic exponent in characterizing glioma-induced hyperexcitability in the human brain. The integrated analysis of electrophysiology, transcriptomics, and behavior provides a comprehensive approach to understand better how gliomas disrupt the cortical balance of excitation and inhibition. The aperiodic exponent holds considerable promise as a novel biomarker with the potential to refine glioma classification, track the course of disease, and pave the way for more effective treatments to restore neuronal circuit homeostasis.

## Supporting information

Sibih_Supplementary Figures

## Acknowledgements

We thank Anny Shai and the staff of the UCSF Brain Tumor Center Biorepository and Pathology Core, Annie Poon, Catherine Shu, and the staff of the UCSF Genomics CoLab and UCSF Center for Advanced Technology (CAT). Figures 3A, 4D, 5A and extended data figures 1A and 7A were made with BioRender.com. Lastly, we are deeply grateful to the patients, whose participation made this research possible.

## Data Availability Statement

Raw single-nucleus RNA sequencing data from IDH-mutant and IDH-wildtype glioma cortex samples (n=14) will be made available on Gene Expression Omnibus (GEO) prior to publication. All other data are available in the article, source data, or from the corresponding author upon reasonable request.

## Funding

This study was supported by grants from the NIH: National Institute of Neurological Disorders and Stroke (NINDS) R01-NS137850 (S.L.H.J, D.B), NINDS R25-NS143071-01 (S.L.H.J), Tom Paquin Brain Cancer Research Fund (S.L.H.J), and Oligo Nation (S.L.H.J).

## Author Contributions Statement

All authors made substantial contributions to the conception or design of the study; the acquisition, analysis, or interpretation of data; or drafting or revising the manuscript. All authors approved the manuscript. All authors agree to be personally accountable for individual contributions and to ensure that questions related to the accuracy or integrity of any part of the work are appropriately investigated and resolved and the resolution documented in the literature. Y.E.S conceived and designed the study with supervision from S.L.H.J. Human electrophysiology data was preprocessed by J.K and analyzed by Y.E.S and N.O with supervision from E.C, A.A.A, P.L, D.B, and S.L.H.J. Subdural grid placement was performed by Y.E.S, N.O, S.H, and V.A with supervision from D.B and S.L.H.J. Single-nucleus RNA sequencing data were preprocessed and analyzed by A.O.D and Y.E.S with guidance from A.D, S.K, K.M, and D.R.R. MEG data were compiled and analyzed by V.J and Y.E.S with supervision from S.N. Immunofluorescence staining and imaging of human FFPE tissue was done by Y.E.S and C.N.G. Co-culture and organoid experiments were performed by S.O. Pharmacologic modulation analysis was conducted by Y.E.S and A.O.D with guidance from G.U. The study was supervised by K.M, P.L, E.F.C, D.R.R, S.N, D.B, and S.L.H.J. The Manuscript was prepared and written by Y.E.S and S.L.H.J with input from all authors.

## Competing Interests

The authors declare no competing interests.

## Code Availability

All open-source software, analysis tools, and packages used in this study are referenced in the Methods and include MNE-Python (v1.3.1), EEGLAB (v2024.2.1), MATLAB (v2024b), specparam (formerly FOOOF; v1.1.0), SciPy (v1.15.1), NumPy (v2.2.2), pandas (v2.2.3), Python (v3.13), R (v4.3.2), RStudio (v2024.09.1+394), Seurat (v5.1.0), MAST (v.3.21), lme4 (v1.1.35.5), lmerTest (v3.1.3), tidyverse (v2.0.0), future (v1.34.0), clusterProfiler (v3.2), fgsea (v1.32.2), InferCNV (v3.20), EnhancedVolcano (v1.24.0), performance (v0.12.4), merTools (v0.6.2), emmeans (v1.10.6), ggridges (v0.5.6), GraphPad Prism (v10.2.3), 10x Cell Ranger (v8.0), QuPath (v0.4.3), ImageJ/FIJI (v1.54p), SynBot (v1.1.3), PsychToolbox 3 (v3.0.18), BrainNet Viewer (v1.7), JMP Pro 18 (v18.1), Nikon NIS-Elements (v5.42.05), SPM12 (v25.01), ANTsR (v0.5.7.6), FreeSurfer (v7.4.1), BrainTRACE (v1.0), Neural Metric Tool (v4.1), MapMyCells (RRID:SCR_024672), Adobe Illustrator (v29.3), Adobe Photoshop (v26.6), and PyCharm – Professional Edition (v2024.1.4). Custom code for electrophysiology analysis is available upon request or hosted on GitHub at https://github.com/sibihy/oneoverf_glioma_e-i.

## METHODS

This research complies with all relevant ethical regulations and was approved by the University of California, San Francisco (UCSF) institutional review board (IRB) for human research (UCSF CHR 17-23215) and performed in accordance with the Declaration of Helsinki. All participants provided written informed consent to participate in this study and to contribute de-identified brain tissue for research.

### Participant characteristics

The study cohort included 13 patients undergoing surgical resection of supratentorial gliomas. World Health Organization (WHO) grade 2-4 gliomas were included and stratified into three subtypes: oligodendroglioma (WHO 2), astrocytoma (WHO 2-3), and glioblastoma (WHO 4). All study participants were individuals seeking care for presumed diffuse glioma at the University of California, San Francisco. Each participant in this study was recruited from a prospective registry of adults aged 18-85 with newly diagnosed, frontal, temporal, parietal, and insular glioma. Inclusion criteria was patients with suspected brain tumor on magnetic resonance imaging (MRI) and spatially mapped collected cortical tissue. Exclusion criteria was any vulnerable population: pediatric patients (age < 18 years old). Intraoperative electrocorticography (ECoG) recordings were obtained during awake craniotomy while at rest, and during visual confrontation naming (picture naming) and auditory naming cognitive tasks. Demographic and neuropathological details, including tumor molecular classification are summarized in Figure 1 and Table 1,. Tumor molecular characteristics were classified per the 2021 WHO Classification of Central Nervous System Tumors. IDH-1 R132H mutation status was assessed via immunohistochemistry, while 1p/19q codeletion was determined via fluorescence in situ hybridization. Spatially annotated normal-appearing and glioma-infiltrated cortex tissue were collected prospectively and split into components to process in liquid nitrogen and for formalin-fixation and paraffin-embedding (FFPE).

### Electrophysiologic signal acquisition

Participants with infiltrative gliomas affecting the language network underwent awake craniotomy using standard neurosurgical techniques. The craniotomy approach was determined based solely on clinical necessity to ensure adequate exposure of the tumor and surrounding cortical and subcortical structures at risk during resection. For patient comfort, a continuous infusion of rapidly metabolized anesthetic agents such as propofol and/or dexmedetomidine was administered during the initial stages of the procedure. All anesthetics were discontinued at least 15 minutes prior to neurophysiological testing to allow for pharmacologic washout. Following this period, a brief wakefulness task was performed and compared to preoperative baseline assessments to confirm adequate alertness and task engagement before data collection. To record intracranial electrophysiological activity, subdural electrocorticography (ECoG) grids were temporarily placed over the exposed cortical surface. Depending on the clinical indication and anatomical constraints, low (4×5 electrodes; 10mm spacing) and high-density (8×12 electrodes; 5mm spacing) grid arrays (Ad-Tech, Oak Creek, WI) were used.

### Intraoperative language testing and protocol

On the day of the procedure, a structured anesthesia washout period of at least 15 minutes was enforced, followed by an extensive wakefulness assessment to confirm adequate arousal for intraoperative language testing. To ensure optimal signal quality, all nonessential auditory and mechanical disturbances were minimized, including muting alarms, pausing surgical suction, and restricting verbal communication among operating room personnel. Stimuli were presented on a 15-inch laptop (60 Hz refresh rate) positioned 30 cm from the participant, running a MATLAB-based script using PsychToolbox 3^88^. The task consisted of a single block of 48 unique stimuli, each depicting a common object or animal as a colored line drawing. Images occupied 75% of the screen and were centrally presented. Participants were instructed to verbally name each item upon presentation. A total of 28 correct responses were monosyllabic, while 20 were polysyllabic. Stimuli were manually advanced by the clinician either upon the participant’s response or after 6 seconds in the absence of a response. A synchronization trigger was sent to the biosignal acquisition system (g.tec) at the onset of each stimulus to facilitate speech latency analysis. An identical approach was utilized during the auditory naming task. The task consisted of a single block of 32 unique stimuli, each depicting an auditory stimulus describing an object. For example, “an animal that barks”. To ensure familiarity with the task, participants completed a training session two days before surgery.

### Grid registration and identification of normal-appearing versus glioma-infiltrated electrodes

To determine whether an electrophysiologic signal originated from glioma-infiltrated or normal-appearing cortex, we employed a multimodal localization approach integrating intraoperative photography, stereotactic neuro-navigation, and structural MRI. For each patient, high-resolution intraoperative images were obtained with and without subdural electrocorticography (ECoG) grids to facilitate precise anatomical registration. These images were aligned to individual pial surface reconstructions and vascular landmarks to extract three-dimensional electrode coordinates. Electrode positions were then co-registered to postgadolinium T1-weighted and T2-FLAIR MRI, and each site was classified as “normal-appearing” or “glioma-infiltrated” by an investigator blinded to the electrophysiologic data. Glioma-infiltrated regions were defined based on established radiographic criteria, characterized by areas of abnormally increased signal on T2-FLAIR imaging without central contrast enhancement on T1-postgadolinium sequences. This classification was further corroborated by intraoperative visual inspection, assessing abnormal vascular patterns and cortical surface morphology indicative of tumor involvement. To ensure precision, electrode classifications were reviewed by a senior investigator and validated against intraoperative stereotactic neuro-navigation records (Brainlab, Munich, Germany). Enantiomorphic normalization was applied to correct for potential brain shift due to edema or mass effect. All spatial transformations, including MRI-to-template alignment, were conducted using SPM12 and ANTsR^89^, while cortical surface reconstructions were generated with FreeSurfer^90^. Electrodes were ultimately mapped to a standardized MNI template for intersubject comparisons, and anatomical regions of interest were assigned according to the Automated Anatomical Labeling atlas. Accuracy of final electrode localization was independently verified by a research technician and a board-certified neurosurgeon, cross-referencing sulcal and gyral anatomy with intraoperative images. This method provided a robust framework for defining glioma-infiltrated versus normal-appearing cortical regions, allowing for systematic analysis of tumor-induced electrophysiologic alterations.

### Electrocorticographic Signal Processing and Spectral Decomposition

During recordings, patients remained awake with their eyes closed to minimize visual input while ensuring wakeful rest. Neural signals were acquired at an initial sampling rate of 4800 Hz, amplified, and imported into Python for preprocessing. Data were down sampled to 1200 Hz, and artifact removal procedures were applied using MNE-Python^91^ (v1.3.1, https://mne.tools/stable/index.html). Raw ECoG recording channels were visually inspected using EEGLAB^92^ (https://eeglab.org/), and excessively noisy recordings were manually rejected. Remaining channels were re-referenced to the common average to reduce global noise contributions. For language-task data, epochs were aligned to stimulus presentation and extracted for 500ms post-stimulus windows during picture-naming and auditory-naming tasks. Electrodes overlying tumor-infiltrated and normal-appearing cortex were analyzed separately as well as by task performance. Preprocessed ECoG recordings were imported in EEGLAB (.set) format using MNE’s read_raw_eeglab function with data preloaded into memory to allow for direct manipulation. To minimize power-line noise contamination, a multi-band notch filter was applied at the fundamental frequency (60 Hz) and its harmonics (120, 180, and 240 Hz) using a zero-phase finite impulse response (FIR) filter implemented in MNE’s notch_filter function. This selective notch filtering approach was chosen to preserve neural signal integrity while attenuating power line interference. Channels were also high-pass filtered at 0.01 Hz to preserve slow dynamics while eliminating drift. Power spectral density (PSD) estimates were computed using Welch’s periodogram method, which reduces noise in the estimated power spectra by averaging periodograms from overlapping segments. For each electrode channel, the time series was segmented into 3-second epochs with 50% overlap between consecutive segments. The segmentation parameters were selected to balance frequency resolution and estimation variance. For each segment, a periodogram was calculated after applying a Hanning window to minimize spectral leakage. PSDs for each electrode and condition of interest (in the case of behavioral analyses) were forwarded to analyses using the specparam toolbox for parameterization of neural power spectra.

### Parameterization of Aperiodic Neural Activity in High Gamma Frequency Range Using specparam

Neural power spectral densities were analyzed to characterize the aperiodic components of neural oscillations using the specparam (formerly FOOOF, or “Fitting Oscillations & One-Over F”) toolbox^25^ (v1.1.0) implemented in Python 3.13. For each electrode, a targeted interpolation approach was employed to address potential artifacts from the 120 Hz harmonic of line noise, which persists as an artifact in the log spectrum after notch filtering (i.e., data points at exactly 120 Hz were linearly interpolated to prevent distortion of the model fit, using SciPy’s interp1d function with the ‘linear’ method and extrapolation enabled for boundary frequencies). Spectral parameterization was performed separately over the ranges [30-50 Hz] and [70-150 Hz], with the following parameter constraints in both cases: peak width limits were set to 0.5-12.0 Hz to capture physiologically plausible oscillatory components; minimum peak height threshold was established at 0.05 (logarithmic units) with a peak detection threshold of 2.0 standard deviations above the aperiodic component; the aperiodic mode was fixed to a ‘fixed’ approach rather than ‘knee’ to optimize the fitting for our frequency range of interests where the power spectrum typically exhibits a linear decay in log-log space. Following model fitting, the aperiodic parameters were extracted, specifically the spectral exponent (slope) and offset (intercept) estimates.

### Event-related spectral analysis of post-stimulus ECoG during language processing tasks

Event-related spectral analyses were performed on ECoG data collected during picture naming (PN) and auditory naming (AN) language tasks to evaluate neural responses in the 500 ms post-stimulus period. Raw ECoG recordings were imported from EEGLAB (.set) files using the MNE-Python package. To minimize electrical line noise contamination, a zero-phase finite impulse response (FIR) notch filter was applied at the fundamental frequency of 60 Hz and its harmonics (120, 180, and 240 Hz) using the ‘firwin’ filter design method. Task-related event markers were extracted from the ECoG recordings using MNE’s events_from_annotations function, with the first event marker in each recording excluded to eliminate potenti‘l init’alization artifacts. Each neural response was segmented into 500 ms post-stimulus epochs, time-locked to the presentation of either visual (PN task; 48 trials) or auditory (AN task; 32 trials) stimuli, with precise sample-level alignment. For each epoch, PSD estimates were computed using the periodogram method with a Hann window to minimize spectral leakage at frequency boundaries. The periodogram approach was selected over Welch’s method to maximize frequency resolution within the short 500 ms epochs, as this duration precluded effective signal averaging across multiple segments. Spectral decomposition was performed with scaling set to ‘spectrum’ to focus on the power distribution across frequencies rather than absolute power values. Each channel’s spectral response was computed independently to preserve spatial specificity, and trials were categorized based on behavioral performance (correct vs. incorrect responses) using a binary encoding scheme (1 for correct, 0 for incorrect). The individual trial PSDs were separately averaged within performance categories for each electrode channel to generate condition-specific spectral representations. This approach allowed for direct comparison of spectral features between successful and unsuccessful language processing attempts. For validation purposes, an alternative spectral estimation approach was implemented using the fast Fourier transform (FFT) with Hann windowing, mean removal, and appropriate scaling. This method computed power as the squared magnitude of the complex Fourier coefficients, with adjustment for one-sided spectral representation by doubling the power of non-DC components except at the Nyquist frequency in even-length signals. The frequency vectors and channel-specific power spectral densities were exported as separate NumPy^93^ (.npy) binary files to facilitate subsequent analysis of task-specific spectral signatures across different frequency bands. The resulting spectral data were organized in a structured directory hierarchy according to subject identifier, task type, and performance condition to enable systematic cross-subject and cross-condition analyses.

### Parameterization of aperiodic neural activity in language task-related high gamma responses

Power spectral density estimates from 500 ms post-stimulus epochs during language tasks were further analyzed to parameterize the aperiodic components of neural activity using the specparam toolbox. For computational efficiency, frequency vectors were loaded once from a shared file (‘frequencies_psd.npy’) and validated to ensure absence of non-numeric values (NaN), infinite values, or empty arrays that could compromise model fitting. Individual channel power spectra were then sequentially processed with rigorous validation at each step. Trials were categorized using a binary coding scheme (0 for correct responses, 1 for incorrect responses) embedded in the filename structure, enabling direct comparison of spectral characteristics between successful and unsuccessful language processing attempts. To mitigate potential residual effects from line noise, data points at exactly 120 Hz were systematically excluded from all power spectra. This targeted filtering approach was implemented to prevent distortion of the aperiodic model fit by persistent harmonic artifacts that could have remained despite prior notch filtering. Linear interpolation was subsequently applied using SciPy’s interp1d function with extrapolation enabled at boundary conditions to ensure uniform frequency spacing across the filtered spectrum. Interpolated frequency vectors were constructed to span from the minimum to maximum of the filtered range while preserving the original frequency resolution. The parameterization of neural power spectra was performed using a fixed aperiodic mode in FOOOF with stringent parameter constraints: peak width limits were set to 0.5-12.0 Hz to capture physiologically plausible oscillatory components; minimum peak height threshold was established at 0.05 (logarithmic units); and peak detection threshold was set to 2.0 standard deviations above the aperiodic component to identify significant oscillatory features. The model fitting was restricted to the high gamma frequency range (70-150 Hz). For each electrode channel, the spectral exponent (|1/f slope|) and offset parameters were extracted from the aperiodic component of the FOOOF model. These parameters quantify the 1/f-like decay of the power spectrum and the overall power magnitude, respectively, which have been theoretically linked to excitatory-inhibitory balance in cortical circuits. In instances where power spectra contained invalid values, placeholder “empty” markers were inserted to maintain the data structure integrity while excluding compromised data points from subsequent statistical analyses. The extracted parameters were organized by electrode identifier and trial type, with additional computation of average slope and offset values for each response category (correct vs. incorrect).

### Epoch-matched spectral analysis of resting state ECoG as neurophysiological baseline

To establish a neurophysiological baseline for comparison with task-related neural activity, resting state ECoG recordings were processed using a temporal segmentation approach matched to the task-related analysis pipeline. Raw resting state ECoG data were imported from EEGLAB (.set) format using MNE-Python and subjected to identical preprocessing as the task data, including notch filtering at line noise frequencies (60, 120, 180, and 240 Hz) using a zero-phase finite impulse response filter with ‘firwin’ design. The continuous resting state recordings were then systematically segmented into multiple non-overlapping epochs of 500 ms duration to match the post-stimulus analysis window used in the language tasks. For picture naming control analysis, 48 epochs were extracted, while 32 epochs were used for auditory naming control analysis, maintaining consistency with the respective numbers of trials in each task. These epochs were distributed equidistantly throughout the resting state recording to ensure comprehensive sampling across the entire recording period, with epoch start points calculated using linear spacing between the beginning and end of the recording while preserving sufficient margin to accommodate complete 500 ms windows. For each epoch, power spectral density estimates were computed using the periodogram method with Hann window tapering to minimize spectral leakage, matching the spectral estimation approach applied to task data. The computational implementation used SciPy’s periodogram function with scaling set to ‘spectrum’ to maintain methodological consistency with the task-related analysis. Spectral computations were performed independently for each electrode channel to preserve spatial specificity of the neural signals. The resulting power spectra from all epochs were averaged for each channel to generate a robust estimate of baseline spectral properties that minimized the influence of transient fluctuations in the resting state. Unlike the task-related analysis, which categorized trials based on behavioral performance, the resting state epochs were treated as a homogeneous condition representing baseline neural activity in the absence of task demands. Channel-specific frequency vectors and corresponding power spectral density estimates were stored as separate NumPy (.npy) binary files to facilitate direct comparison with task-related spectral features using identical analytical approaches. The directory structure for data storage followed the same organizational scheme as the task data, with naming conventions that explicitly identified the data as resting state control recordings. This methodologically matched approach to spectral analysis between task and rest conditions enabled direct statistical comparison of neural spectral properties across cognitive states while controlling for subject-specific anatomical and physiological factors.

### Machine-learning models

To determine the most effective classification algorithm, we utilized JMP 18 Pro’s Model Screening feature, which allows for systematic evaluation of multiple machine learning approaches on a given dataset. Initial model comparisons included random forest, logistic regression, boosted trees, support vector machines (SVM), and neural networks. Each model was assessed based on misclassification rate, area under the receiver operating characteristic curve (AUC), and cross-validation performance. The random forest algorithm consistently demonstrated the highest classification accuracy across all tasks, particularly in handling nonlinear relationships between aperiodic exponent values and glioma classification outcomes, while also being robust to overfitting due to its ensemble learning approach. The model’s ability to handle high-dimensional data, capture complex interactions, and provide interpretable feature importance rankings made it the optimal choice for distinguishing normal vs. glioma-infiltrated electrodes and glioma subtype classifications.

For classification of glioma-infiltrated versus normal-appearing cortex and glioma subtype prediction, we implemented random forest classifiers utilizing aperiodic exponent values. These classifiers were designed to assess whether electrophysiological features alone could accurately differentiate between normal and tumor-infiltrated cortex and classify glioma subtypes. The first classifier was developed to distinguish glioma-infiltrated electrodes from normal-appearing electrodes based solely on the aperiodic exponent’s magnitudes. Electrodes were labeled according to preoperative T2-FLAIR hyperintensity, with glioma-infiltrated electrodes classified as glioma-infiltrated and electrodes overlying structurally normal cortex classified as normal-appearing. The second classifier predicted glioma subtype (oligodendroglioma, astrocytoma, GBM) using the aperiodic exponent as the primary predictive feature to determine whether electrophysiological markers alone could differentiate tumor subtypes.

All models were built and optimized in JMP 18 Pro, which provided an interactive framework for random forest modeling, feature selection, and hyperparameter tuning. Model training and validation were performed using leave-one-patient-out cross-validation (LOPO-CV) to minimize overfitting and account for inter-patient variability. Hyperparameter tuning was conducted to optimize the number of trees, maximum depth, and minimum node size based on cross-validation performance. Model evaluation metrics included receiver operating characteristic (ROC) curves, area under the curve (AUC) scores, precision-recall metrics, and misclassification rates.

### Time-off anesthesia analysis

To quantify the pharmacokinetics of anesthetic washout following infusion cessation, we implemented a first-order exponential decay model to estimate residual concentrations of both propofol and dexmedetomidine in glioma patients undergoing intraoperative electrocorticography (ECoG). Drug clearance was modeled using a single-compartment elimination framework with medication-specific rate constants: k = 0.119 min⁻¹ for propofol and k = 0.006 min⁻¹ for dexmedetomidine, based on published pharmacokinetic data^94,95^. At any time t after infusion cessation, plasma concentration was defined as C(t) = C₀e⁻ᵏᵗ, where C₀ represents the estimated steady-state concentration. Volume of distribution (Vd) was also adjusted for sex (propofol: 75 L for males, 65 L for females; dexmedetomidine: 109 L for males, 94 L for females).

Initial concentrations were calculated per patient using documented infusion rates and patient-specific factors (sex and weight), with steady-state verified as ≥4 half-lives of infusion duration. Full intraoperative infusion records for each medication were manually curated and converted from military time to absolute minutes post-midnight. Events were classified as bolus or continuous infusion, and infusion rates were standardized to mcg/kg/min. For each patient and medication, we computed the dynamic decay in estimated plasma concentration across the full intraoperative wakefulness period, including the timing of a wake-up task, a 3-minute resting-state ECoG baseline, and two functional language tasks: (1) Picture Naming (PN; 48 trials) and (2) Auditory Naming (AN; 32 trials). For patients receiving both agents prior to infusion cessation (n = 2 patients), propofol and dexmedetomidine concentrations were modeled independently. The estimated concentration at each minute post-infusion was computed for every patient and aligned with task timing. Visualization of concentration decay curves was performed for each drug individually, enabling assessment of residual anesthetic burden (Extended Data Fig. 4). To quantify neural recovery, aperiodic exponent values were computed from ECoG recordings during rest, PN, and AN tasks, and correlated with modeled anesthetic levels across time. This approach enabled patient-specific mapping of excitation-inhibition balance dynamics as sedation waned.

### Resting-state MEG acquisition, source localization, and spectral exponent quantification in patient-matched validation cohort

Resting-state magnetoencephalography (MEG) was recorded from each participant for 10 minutes in the supine position, eyes closed, and awake, using a 275-channel whole-head CTF Omega 2000 system (CTF MEG International Services LP, Coquitlam, BC, Canada) at the UCSF Biomagnetic Imaging Laboratory. Recordings were sampled at 600 Hz as part of a standardized protocol for functional mapping in patients with brain tumors. Head position relative to the sensor array was determined using three fiducial coils (nasion and bilateral preauricular points), which were subsequently co-registered to each subject’s high-resolution MRI to enable precise anatomical localization. To ensure signal quality, head motion was restricted to <5 mm during acquisition. From each subject’s recording, a continuous 60-second artifact-free segment was selected and bandpass-filtered between 1–50 Hz. An 8 mm isotropic voxel grid was generated within a standard MRI template space (∼11,000 voxels) and spatially transformed to individual native MRI space. Forward modeling was conducted using a single-shell approximation of the head model^96^, and subject-specific magnetic lead field vectors were calculated for each voxel.

Tumor-infiltrated regions were manually delineated based on FLAIR hyperintensities, co-registered with structural post-contrast T1- and T2-FLAIR weighted images. Contralateral, anatomically homologous regions lacking radiographic abnormalities were defined as control (healthy) tissue for within-subject comparisons of electrophysiological activity. Source-level time series were reconstructed using scalar synthetic aperture magnetometry (SAM) beamforming^97^. The data covariance matrix was computed across the full 60-second segment, and optimal source orientation was estimated by solving a generalized eigenvalue problem^98^. Lead fields were normalized to mitigate central localization bias. All beamformer computations were performed using the CTF SAM source reconstruction pipeline.

To quantify voxel-specific neural excitability, we analyzed the aperiodic spectral exponent derived from the 30– 50 Hz frequency band. This range was selected based on prior evidence suggesting that spectral flattening in higher frequencies reflects increased excitability of underlying cortical microcircuits. For each voxel, the power spectrum was computed from the reconstructed time series, and the aperiodic component—representing the 1/f-like structure of the spectrum—was isolated using a robust three-step least-squares fitting approach. The spectral exponent (α) was defined as the slope of the linear regression on the log–log transformed spectral power versus frequency plot.

Voxel-wise Z-scores were calculated for each participant by comparing tumor-infiltrated regions to their contralateral homologues, using a threshold of |Z| > 1.96 (p < 0.05; Extended Data Fig. 6). For group-level analyses stratified by molecular tumor subtype, individual Z-maps were transformed into Montreal Neurological Institute (MNI) space. For consistent hemispheric visualization, right-hemispheric tumors (Subjects 7, 10, and 12) were mirrored to the left. Three-dimensional renderings of both subject-specific and group-level aperiodic slope distributions were visualized using the BrainNet Viewer toolbox^99^, overlaid on the Brainnetome atlas for cortical and subcortical parcellation.

### Tissue collection and processing

Cortex tissue samples (∼2-8 pieces per surgery) resected from patients with a known or suspected oligodendroglioma, astrocytoma, and glioblastoma was transferred from the operating room to the laboratory on ice with a target cold ischemia time of 30 minutes. All tissue samples collected had stereotactic coordinates of their location logged and were retrieved from the Brainlab curves. Normal-appearing cortex samples were collected from patients undergoing insular tumor resection. Glioma-infiltrated cortex samples were collected from patients who had T2 flair-positive sites as previously defined. Tissue samples were processed immediately upon arrival to the laboratory. Samples were split into two halves. One half was flash frozen in liquid nitrogen for storage and the other half was placed into a tissue processing and embedding cassette designed to securely hold specimens. The cassettes were placed into a 4oz specimen cup filled with 10% formalin. The specimen cup was then transferred to a 4C fridge for overnight incubation. Formalin was replaced with 70% ethanol after 24 hours had passed.

### Assigning tissue samples nearest electrode 1/f value

Electrode coordinates were extracted following intraoperative electrode grid registration using BrainTRACE (Brain Tumor Registration and Cortical Electrocorticography)^100^, a software platform that enables precise mapping of subdural electrodes onto a patient’s cortical surface based on intraoperative photography and preoperative MRI. The patient-specific MRI registered coordinates (x, y, z) of all implanted electrodes were exported into a CSV file, preserving their anatomical positions relative to the brain surface for each patient. Tissue sample three-dimensional coordinates were recorded intraoperatively using Brainlab neuro-navigation, which enabled real-time tracking of biopsy locations relative to preoperative imaging. These coordinates were saved in the operating room at the time of tissue collection and later used for spatial mapping. To establish a spatial correspondence between the sampled tissue and the nearest intracranial electrodes, a nearest-neighbor analysis was performed by computing the Euclidean distance between each tissue sample and every electrode in the corresponding patient’s grid. The Euclidean distance was calculated using √(x_1_-x_2_)^2^+(y_1_-y_2_)^2^+(z_1_-z_2_)^2^, where (x_1_, y_1_, z_1_) represent the tissue sample coordinates and (x_1_, y_2_, z_2_) correspond to the electrode coordinates. This calculation was applied iteratively for all electrodes within a patient’s grid, and the four closest electrodes to each tissue sample were identified and ranked by proximity. The electrode grid ID corresponding to each of the four nearest electrodes, along with the respective Euclidean distances, was recorded for each sample to ensure accurate alignment between electrophysiological recordings and molecular tissue analyses. This spatial correspondence enabled direct integration of intracranial ECoG recordings with single-nucleus RNA sequencing (snRNA-seq) data, supporting a multimodal investigation of glioma-induced alterations in excitation-inhibition balance. All computations were performed in Python v3.13 using the pandas and numpy packages, with results stored in a structured data frame for subsequent statistical analysis and visualization.

To assign an electrophysiological signature to each tissue sample, the aperiodic exponent was computed for all electrodes and mapped onto the corresponding tissue coordinates. For each tissue sample, the 1/f slope values from the four nearest electrodes were averaged, ensuring that the assigned value reflected local cortical activity rather than isolated electrode variability. This averaged aperiodic exponent was then assigned to the corresponding tissue sample, allowing for direct integration of electrophysiological and transcriptomic data. To identify biologically meaningful differences in cortical excitation-inhibition balance, we selected the two most extreme tissue samples per patient, defined as the sample with the lowest and highest 1/f slope within that patient’s dataset. These samples were categorized as “flat 1/f” or “steep 1/f” relative to the patient’s own cortical landscape, establishing an intra-patient reference rather than an absolute classification. To systematically examine glioma-induced alterations in cortical activity, we analyzed a total of 12 glioma-infiltrated tissue samples from six patients (two patients per glioma subtype: oligodendroglioma WHO 2, astrocytoma WHO 2, and glioblastoma). Additionally, we included two normal-appearing cortex samples from a single patient who was undergoing tumor resection of an insular glioblastoma, providing a reference for normal electrophysiological and transcriptomic profiles. This approach enabled the comparison of excitation-inhibition dynamics between different glioma subtypes and allowed for the identification of molecular signatures associated with cortical hyperexcitability in glioma-infiltrated versus normal cortical regions. By integrating electrophysiological and transcriptomic measures at a spatially resolved level, this framework provided a robust strategy for investigating the neurobiological underpinnings of glioma-associated network dysfunction.

### Nuclei isolation from frozen tissue and single-nucleus RNA sequencing pre-processing

For the single-nucleus RNA sequencing (snRNA-seq) analysis, we included two biologically independent patients per glioma subtype for our glioma-infiltrated samples (n = 2 oligodendroglioma, n = 2 astrocytoma, n = 2 glioblastoma) and one patient undergoing tumor resection for an insular glioblastoma for our normal-appearing cortex samples. A total of 14 cortex specimens spanning three major molecular subtypes were sequenced. Glioma infiltrated cortex samples: oligodendroglioma (n=4), astrocytoma (n=4), glioblastoma (n=4); normal-appearing cortex samples (n=2). Specimens were classified as steep 1/f (n=7) or flat 1/f (n=7) based on previous 1/f signal analysis, with each 1/f category containing samples from all cortex and glioma subtypes. Nuclei from frozen tissue were extracted from normal-appearing and glioma-infiltrated cortex samples from patients included in the electrophysiologic analysis using the 10x Genomics Chromium Nuclei Isolation Kit (10x Genomics, PN-1000493). Nuclei viability and counts were assessed using AOPI dye and GFP labeling. Nuclei that stained green were considered live according to the chromium nuclei isolation kit protocol and counts were performed using the Invitrogen Countess 3FL automated cell counter (Invitrogen, A49893). Libraries were prepared with the assistance of the UCSF Genomics CoLab, and sequencing was performed at the UCSF CAT, supported by UCSF PBBR, RRP IMIA, and NIH 1S10OD028511-01 grants on an Illumina NovaSeq 6000 platform with a target depth of 50,000 reads per nucleus.

### Single-nucleus RNA sequencing data integrating, dimensionality reduction, clustering, and cell-type identification

Raw sequence data was processed using Cell Ranger (10x Genomics) to generate filtered gene-barcode matrices in HDF5 format. To efficiently process the high-dimensional data, computing resources were optimized using the ‘future’ R package with multicore processing across 32 CPU cores and 300GB of RAM allocated (plan(“multicore”, workers = 32); options(future.globals.maxSize = 300 * 1024^3)). Individual filtered gene-barcode matrices were imported into Rstudio (v2024.09.1+394) using the Read10X_h5 function from Seurat^101^ v5.1.0 For each sample, a Seurat object was created with the following quality control parameters: minimum of 3 cells per gene, minimum of 200 and maximum of 10,000 features (genes) per cell, and maximum mitochondrial gene expression of 10%. Mitochondrial gene content was calculated using the PercentageFeatureSet function with the pattern “^MT-“ to identify human mitochondrial genes. QC parameters were visualized using violin plots and feature scatter plots to confirm appropriate thresholding. Each Seurat object underwent standard log-normalization (NormalizeData function), data scaling (ScaleData function), and variance stabilizing transformation using SCTransform with the v2 algorithm (vst.flavor = “v2”). Sample metadata was appended to each object including experimental identifier, glioma molecular subtype (oligodendroglioma, astrocytoma, or glioblastoma), and 1/f status (steep1overf or flat1overf). To correct for batch effects and enable joint analysis across all samples, we implemented a reference-based integration approach using reciprocal PCA (rPCA). The integration workflow consisted of selecting 3,000 highly variable genes as anchor features (SelectIntegrationFeatures function with nfeatures = 3000), preparing the RNA assay for integration (PrepSCTIntegration function), running PCA on each object using the selected feature set, and finding integration anchors using dimensionality reduction (FindIntegrationAnchors function with normalization.method = “SCT”, dims = 1:30, reduction = “rpca”, and k.anchor = 20). Copy number variation analysis was performed with the InferCNV R package (https://www.bioconductor.org/packages/release/bioc/html/infercnv.html).

A total of 508,899 cells were identified and subsequently assigned to 34 distinct clusters. Clusters were identified using the FindAllMarkers function in Seurat with adjusted P < 0.05 (Wilcoxon rank-sum test with Bonferroni correction), log fold-change threshold of 0.5, and analysis limited to genes expressed in at least 50% of cells (min.pct). Canonical cell population identities were verified through literature review. Neuronal subpopulations were further classified using Allen Brain Institute’s MapMyCells graphical user interface leveraging the 10x Human MTG SEA_AD (CCN20230505) reference taxonomy and the SEA_AD Hierarchical Mapping algorithm (MapMyCells; RRID:SCR_024672). A final subset of 84,825 neurons (52,763 steep 1/f and 32,062 flat 1/f) were included in primary analysis.

### Excitatory module score calculation

Excitatory genes included^102,103^: *SLC17A6, SLC17A7, SLC17A8, GRIA1, GRIA2, GRIA3, GRIA4, GRIN1, GRIN2A, GRIN2B, GRIN2C, GRIN2D, GRIN3A, GRIN3B, GRM1, GRM2, GRM3, GRM4, GRM5, GRM6, GRM7, GRM8, TBR1, FEZF2, NEUROD1, CTIP2, SATB2, CAMK2A, CAMK2B, NRGN, SYT1, PSD95, DLG4, SHANK1, SHANK2, SHANK3, HOMER1, HOMER2, HOMER3, VAMP1, VAMP2*.

Excitatory scores were computed using the AddModuleScore from the Seurat package. This function calculates the average expression levels of the gene set subtracted by the aggregated expression of 100 randomly chosen control gene sets, where the control gene sets are chosen from matching 24 expression bins corresponding to the tested gene set expression. Statistical comparisons of the average excitatory scores between steep 1/f and flat 1/f neurons across pathologies were performed using Mann-Whitney U Test and one-way ANOVA for neuronal subtype comparisons in Prism.

### Differential gene expression analysis

Differential gene expression between steep 1/f and flat 1/f neurons was conducted using Seurat’s FindMarkers function with default parameters, stratified by pathology. Genes from the excitatory gene set identified as differentially expressed with a log-fold change > 0.25 were specifically reported. Volcano plots illustrating these findings were generated using EnhancedVolcano (10.18129/B9.bioc.EnhancedVolcano) (v1.24.0).

### Gene ontology and gene set enrichment analysis (GSEA)

Gene ontology (GO) analysis was performed using the clusterProfiler R package (v3.2). Differentially expressed genes (log-fold change > 0.25, adjusted P < 0.05) were analyzed using the using the enrichGO function applying thresholds of P-value cutoff = 0.01 and q-value cutoff = 0.05. All GO domains were queried including Biophysical Processes (BP), Cellular Component (CC), Molecular Function (MF), were queried for enrichment.

GSEA was carried out using fgsea (v1.32.2). Differentially expressed genes were interrogated enrichment in “GOCC_EXCITATORY_SYNAPSE.” Additionally, the entire biophysical processes ontology was queried, and results were filtered to include pathways related to neuronal signaling identified by search terms, “NEUROTRANSMITTER”, “GLUTAMATE”, “GABA”, “EXCITATORY”, “INHIBITORY”, “SYNAPSE”, “NEURON”.

### Immunofluorescent staining of human cortex FFPE tissue

Immunofluorescent staining was performed on 5μm formalin-fixed paraffin-embedded (FFPE) human cortex tissue sections from patient cohort using a modified two-day protocol optimized for detection of neuronal markers. On the day of staining, slides were equilibrated to room temperature (RT) for 10 minutes prior to deparaffinization. Slides were processed under a chemical fume hood. Sections were deparaffinized in three changes of xylene (3 minutes each), followed by sequential immersion in 100%, 95%, 80%, and 70% ethanol (2–3 minutes per step), and rinsed in deionized water for 5 minutes. Sections were then transferred to 1× PBS for an additional 5-minute wash. Antigen retrieval was performed using a vertical Coplin jar filled with 1× citrate-based antigen retrieval buffer (pH 6.0; Vector Laboratories), heated in a microwave at high power for 2.5 minutes followed by 10 minutes at low power (10% power). After retrieval, slides were cooled at RT for 20–30 minutes and rinsed in distilled water for 5 minutes. Following antigen retrieval, tissue sections were encircled with a hydrophobic barrier pen and incubated in 3% hydrogen peroxide for 15 minutes to quench endogenous peroxidase activity. Slides were washed twice with distilled water (5 minutes each), then permeabilized with PBS containing 1% animal serum (goat) and 0.4% Triton X-100 for 2 × 10 minutes. Blocking of nonspecific antibody binding was performed with 5% goat serum in PBS-T for 1 hour at RT in a humidified chamber. Slides were incubated for 1 hour at RT with primary antibodies diluted in PBS-T containing 1% goat serum. The following primary antibodies were used: mouse anti-NeuN (1:100, clone A60, Millipore Sigma, MAB377), rabbit anti-IDH1 R132H (1:200, Invitrogen, MA5-44585), rabbit anti-SOX2 (1:300, Cell Signaling Technology, 2748S), and rabbit anti-Homer1 (1:250, Invitrogen, PA5-21487). Negative control sections were incubated with antibody diluent lacking primary antibody. Parafilm was applied to prevent evaporation, and slides were stored overnight at 4 °C in a humidified, light-protected chamber. The next day, slides were washed twice with PBS-T containing 1% serum (10 minutes each) and once with PBS (10 minutes). Sections were incubated at RT for 1–2 hours with the following fluorophore-conjugated secondary antibodies, each diluted 1:500 in PBS-T containing 1% goat serum: Alexa Fluor 488 goat anti-mouse IgG (H+L) (Invitrogen, A-11001) and Alexa Fluor 568 goat anti-rabbit IgG (H+L) (Invitrogen, A11036). After secondary incubation, slides were washed three times in PBS (5 minutes each), then counterstained with DAPI nuclear dye (1 μg/mL; 1:1000 dilution in PBS, Thermo Fisher, 62248) for 10 minutes, followed by a final 5-minute PBS wash. Slides were mounted using Fluoromount-G mounting medium (SouthernBiotech, 0100-01) and imaged.

### Microscopy and quantification of human cortex FFPE tissue

Confocal imaging was performed on immunostained FFPE human cortical sections using an Eclipse Ti2-E inverted microscope (Nikon Instruments Inc.). For each section, six high-resolution fields of view (snapshots) were acquired at 20x magnification (2048 × 2048 pixels) across three anatomically matched tissue sections per slide. Imaging parameters were optimized to preserve tissue integrity and minimize photobleaching. A laser dwell time of 2 μs per pixel was used, with single line averaging applied to enhance signal-to-noise ratio. All imaging conditions, including laser power, gain, and scan speed, were held constant across all sections and samples. For quantification, immunopositive cell counts were performed using QuPath (v0.4.3), an open-source digital pathology analysis platform. Within each image, the number of NeuN+ nuclei and Homer1+/NeuN+ double-positive cells were quantified using threshold-based detection parameters and manually validated segmentation. The proportion of Homer1-expressing neurons was calculated as the percentage of Homer1+/NeuN+ cells relative to total NeuN+ cells within each image. All quantification was performed independently across the six fields of view per section, and values were averaged to yield a per-section estimate of synaptic marker expression. In respect to the Tumor infiltration vs aperiodic exponent analysis, IDH+ and SOX2+ cells were manually identified and counted using the Cell Counter plugin in FIJI, based on clear nuclear or perinuclear immunoreactivity. The total area of the imaged field (μm²) was determined using the freehand selection tool in ImageJ, allowing precise delineation of the imaged region for each sample. Cell counts were normalized to tissue area and expressed as tumor cell density (cells/μm²). All quantifications were performed blinded to electrophysiological data and glioma subtype.

### Tumor organoid development from patient derived glioma tissue

Patient-derived tumor organoids were generated using freshly resected tumor tissues, transported to the laboratory (<30-minute cold ischemia), and maintained for up to 6 hours in Hibernate A medium (Gibco, A1247501). To develop organoids, tissue fragments (∼2-3 mm³) underwent incubation with RBC lysis buffer (Invitrogen, 00-4333-57) for 10 minutes under constant agitation at room temperature, followed by one wash in ice cold Hibernate A supplemented with 1:100 Glutamax (Gibco, A1286001) and 1:100 Anti-Anti (Gibco, 15240062). Tumors were then transitioned into ice cold culture medium comprising 1:1 mixture of Neurobasal Medium (Gibco, 21103049) and DMEM/F12 (Gibco, 11320033), supplemented with 1:50 B-27 without Vitamin A (Gibco, 12587001), 1:100 N-2 Supplement (Gibco, 17502001), 1:100 Glutamax, 1:100 Penicillin-Streptomycin (Gibco, 15140122), low-glutamate non-essential amino acids mixture (Gly, L-Ala, L-Asn, L-Asp, L-Pro, L-Ser = 100μM and L-Glutamic acid = 300nM), 0.05 mM 2-mercaptoethanol (Sigma-Aldrich, 63689), and 2.5 μg/mL insulin solution (Sigma-Aldrich, I9278). Uniform 1 mm fragments were created using biopsy punches and subsequently cultured under hypoxic conditions (37°C, with 5% CO₂, and 5% O₂) on an orbital shaker at 120 RPM for at least 4 weeks. Regular mycoplasma screening was performed using the MycoAlert kit (Lonza, LT07-318).

### Cortical neurosphere (cNS) development from murine cortical tissue

Murine cNS were derived using C57BL/6J E15.5-18.5 embryos from timed-pregnant dams that were sacrificed in accordance with UCSF Institutional Animal Care and Use Committee (IACUC) guidelines. Neuron isolation was performed as previously described^12^. In brief, dissected cortical tissues were finely chopped into approximately 1 mm² fragments and enzymatically dissociated using the Worthington Papain Dissociation System (Worthington Biochemical Corporation, LK003150). Here, tissue fragments underwent incubation with papain solution at 37°C with constant agitation for 7 minutes, after which enzyme activity was halted by aspirating papain and adding the inhibitor solution for 3 minutes at 37°C. Following inhibition, the solution was aspirated, and enzymatic digestion was completed by adding 5 mL of Neurobasal medium containing DNase (Worthington, LK003172; 0.5 mL of 10 mg/mL). The suspension was then centrifuged at 1400 RPM at 4°C for 6 minutes, and the supernatant was discarded and replaced with fresh Neurobasal medium. Subsequently, the cell suspension was filtered through a 40 µm nylon mesh into a fresh 50 mL conical tube to remove residual tissue fragments. The filtrate was centrifuged again at 1400 RPM for 6 minutes at 4°C, and the resulting pellet was resuspended in pre-warmed neuronal culture medium consisting of Neurobasal medium supplemented with N-2 (1:100), Anti-Anti (1:100), B-27 Supplement (1:50), and sodium pyruvate (Gibco, 11360070; 1:100). Cell viability and concentration were assessed using a 1:1 Trypan blue to cell suspension mixture with a hemocytometer. Cells were seeded at a density of 500,000 cells per well in 24-well ultra-low attachment plates containing 1 mL neuronal culture medium. Plates were incubated under hypoxic conditions (37°C, 5% CO₂, and 5% O₂) on an orbital shaker set at 120 RPM, with half-media changes performed every other day.

### Tumor organoid + cNS co-cultures

For the creation of tumor organoid + cNS co-cultures, patient-derived glioma organoids and DIV4 cNS were combined within individual wells containing 1 mL neuronal culture medium. Plates were incubated at a 45° angle under hypoxic conditions to facilitate physical fusion and attachment, typically achieved after approximately 48 hours. Following stable attachment, the plates were returned to orbital shaking at 120 RPM, and all subsequent experimental assays were performed 21 days after initial co-culture establishment.

### Immunofluorescent staining, microscopy, and quantification of organoid co-cultures

For immunofluorescent analysis of tumor organoid + cortical neurosphere (cNS) co-cultures, samples underwent fixation in 4% paraformaldehyde (PFA) at 4°C for 45 minutes with gentle agitation, followed by two hours of phosphate-buffered saline (PBS) washes. Samples were subsequently incubated overnight at 4°C in 30% sucrose solution until fully submerged. The samples were embedded in cryomolds and cryosectioned at a thickness of 12 µm. Before staining, sections were rinsed in PBS for 15 minutes. Nonspecific antibody binding was blocked using 10% normal goat serum (NGS) in PBS at room temperature for one hour, followed by incubation in a permeabilization buffer containing 5% NGS, 0.25% Triton X-100, and primary antibodies in PBS. Primary antibodies used included chicken anti-MAP2 (1:500, EnCor, CPCA-MAP2), rabbit anti-Homer1 (1:250, Invitrogen, PA5-21487), and mouse anti-Synapsin I (1:200, Invitrogen, MA5-31919). Sections were incubated overnight at 4°C. Post-primary incubation, sections were rinsed thrice with PBS (15 minutes each) and incubated with secondary antibody solution containing 1% NGS, 0.25% Triton X-100, Alexa Fluor 488 goat anti-chicken IgY (H+L) (1:500, Invitrogen, A11039), Alexa Fluor 568 goat anti-mouse IgG (H+L) (1:500, Invitrogen, A11004), and Alexa Fluor 647 goat anti-rabbit IgG (H+L) (1:500, Invitrogen, A21245) in PBS for one hour at room temperature. Following secondary incubation, slides underwent three PBS washes (5 minutes each), counterstaining with DAPI (Thermo-Fisher, 62248; 1:1000), and were mounted using Fluoromount-G (SouthernBiotech, 0100-01). Imaging was performed using a Nikon Eclipse Ti2-E inverted microscope for confocal microscopy and image processing was conducted in the Nikon NIS-Elements software (v5.42.05). Imaging conditions included a 40x objective lens with image resolution at 2048 × 2048 pixels. To optimize signal-to-noise ratio, laser dwell time was set at 2 µs per pixel with two rounds of line averaging depending on the assay performed. Synaptic marker analyses (Homer1 and Synapsin1) were performed by quantifying colocalized puncta using SynBot (https://github.com/Eroglu-Lab/Syn_Bot), a FIJI macro optimized for automated detection and analysis of synaptic marker colocalization.

### Organoid electrophysiology

Multielectrode array (MEA) recordings were acquired using 24-well CytoView MEA plates (Axion Biosystems, M384-tMEA-24W, Serial # 11-01.03.00387) with a 4 × 4 electrode array (16 electrodes per well, spaced 350 µm apart). Two days prior to organoid plating, each MEA well was prepared by coating with 100 µL Poly-D-Lysine (PDL; Gibco, A3890401) overnight at 37°C under humidified conditions with 5% CO₂. The PDL solution was then aspirated, and wells were rinsed twice thoroughly using ultrapure distilled water. One day before plating organoids, wells were coated with 100 µL of laminin (mouse protein; Corning, CB-40232) at a concentration of 10 µg/mL in PBS, and incubated overnight at 37°C, 5% CO₂, under humidified conditions.

On the day of plating, the laminin solution was carefully removed without rinsing. Individual organoids (either cNS alone or tumor organoid + cNS co-cultures) were transferred to MEA wells containing 1 mL fresh neuronal culture medium per well. Plates were incubated undisturbed for 48 hours in a humidified incubator at 37°C and 5% CO₂ to facilitate organoid attachment.

Extracellular electrophysiological activity was recorded using a Maestro Edge system (Axion Biosystems). During recordings, MEA plates were maintained at 37°C on a heated stage with continuous ventilation (5% CO₂). Spontaneous neuronal firing was recorded over a 15-minute interval, utilizing a sampling rate of 12.5 kHz per channel, 1000× gain, and a bandpass filter of 200–3000 Hz. Spikes were detected by an adaptive threshold method, defined as 5 standard deviations above the root mean square noise level on each electrode. Electrodes exhibiting a spike rate of ≥5 spikes per minute were categorized as active, and wells were considered active if at least four electrodes displayed activity above this threshold. Data analysis was conducted using the Neural Metric Tool software (Axion Biosystems), quantifying the weighted mean firing rate, defined as the average firing frequency of active electrodes (Hz).

### Statistics and reproducibility

All statistical analyses were conducted using Python (v3.13), GraphPad Prism (v10.2.3), and R (v4.3.2) with RStudio and the tidyverse suite^104^(v2.0.0) for data cleaning, wrangling, and visualization. Linear mixed-effects models (LME) were performed using the lmerTest and lme4 packages in R^105^ to account for repeated measures across electrodes within subjects. In all LME models, patient ID was included as a random intercept to account for intra-subject correlation. Model outputs include fixed effect estimates (β), standard error, degrees of freedom (df), t-statistics, 95% confidence intervals (CI), and p-values computed using Satterthwaite approximation. Where appropriate, Welch’s t-tests were used for unequal variance comparisons, Mann– Whitney U tests were employed for non-normally distributed group comparisons, and one-way ANOVA with Šídák’s or Tukey’s post hoc tests were used for multi-group comparisons. Model fits were evaluated using AIC and BIC values. For imaging-based z-statistical mapping of MEG aperiodic slope, z-scores were computed voxel-wise within patients to localize spectral deviations relative to contralateral control cortex.

All n-values refer to biologically independent samples (i.e., patients, electrodes, tissue regions) unless otherwise specified. No statistical methods were used to predefine sample size; however, cohort size is consistent with previous intraoperative electrophysiology and human brain tumor studies. Quantification in immunofluorescence experiments was performed across six fields of view from three adjacent FFPE sections per sample. All quantification was conducted blinded to sample identity using QuPath and ImageJ (v1.54p). For single-nucleus RNA-seq analyses, cell-type assignments were based on canonical markers, and differential expression was performed across neuronal populations. Excitatory module scores were compared using Mann-Whitney U test, and gene set enrichment was performed using clusterProfiler and fgsea R packages. All tests were two-tailed, and statistical significance was set at p < 0.05, unless otherwise noted. Full statistical results including model coefficients, p-values, and confidence intervals are reported in the text, and figure legends.

---

**Extended Data Fig. 1.**
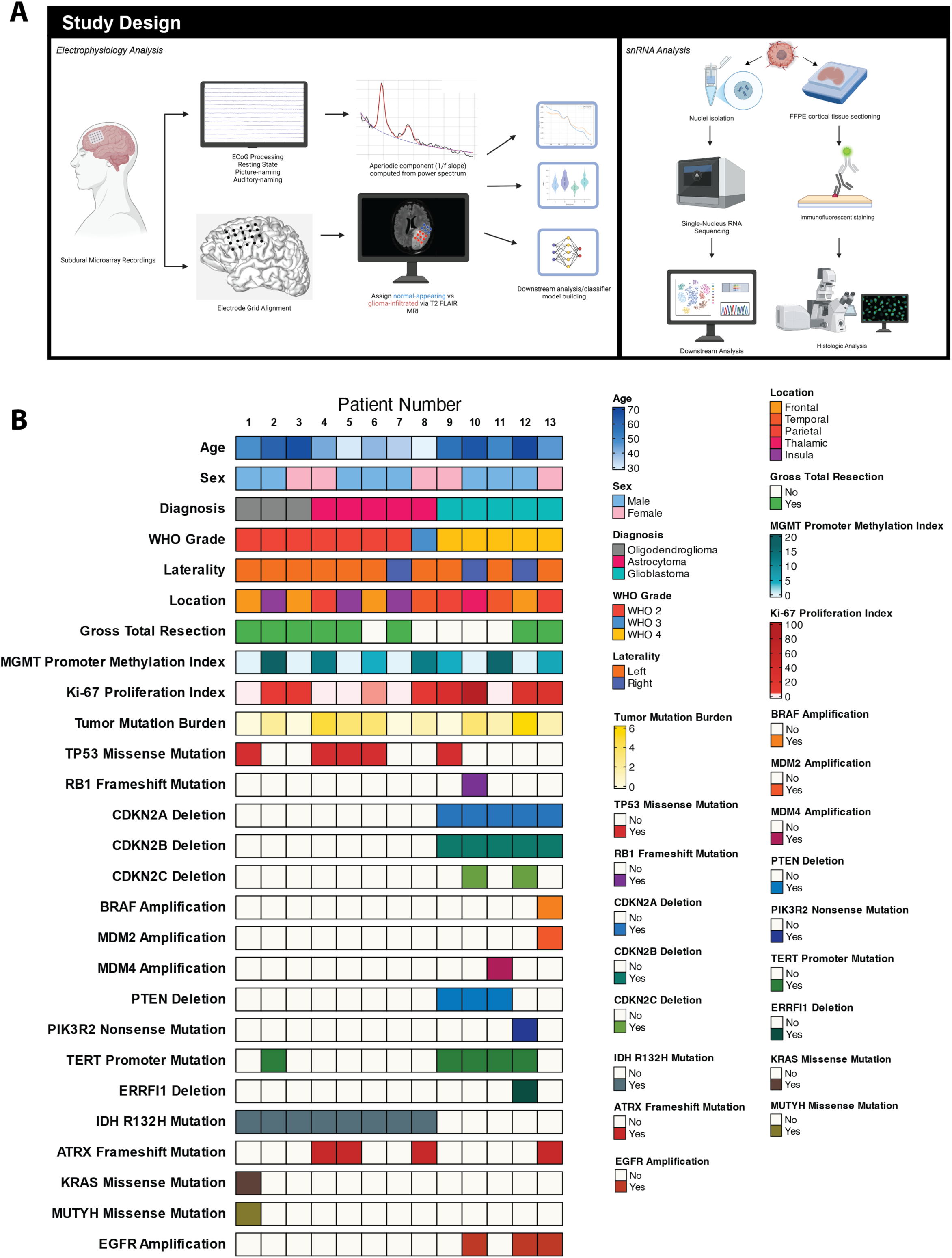
Over of study design and patient histologic, molecular, and clinical features with complete list of next-generation sequencing targets. **A,** Left: Electrophysiology analysis workflow. Right: Single-nucleus RNA sequencing analysis pipeline. **B,** Extended oncoprint displaying full list of targeted next-generation sequencing genes identified with targeted next-generation DNA sequencing.

**Extended Data Fig. 2.**
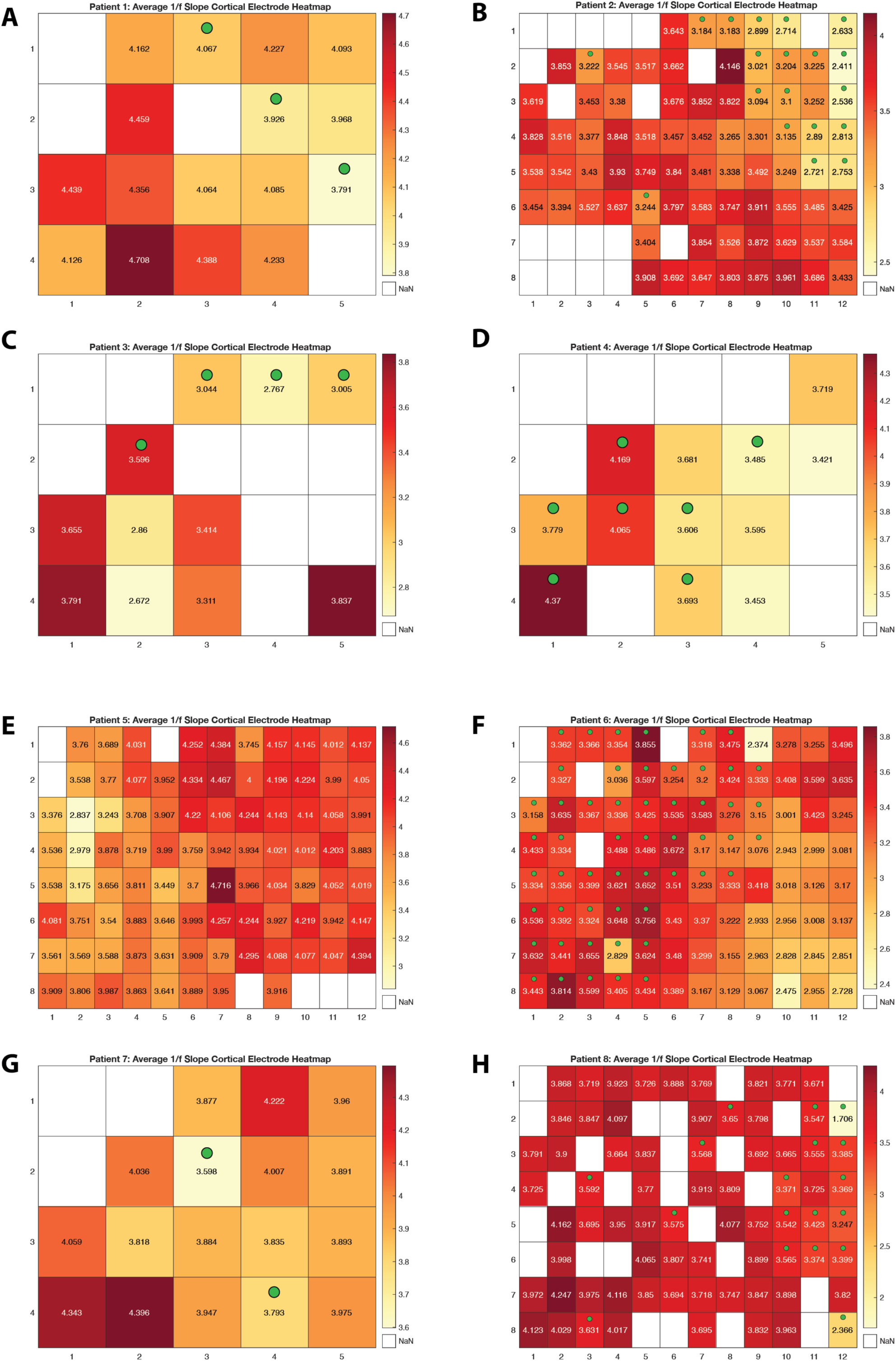
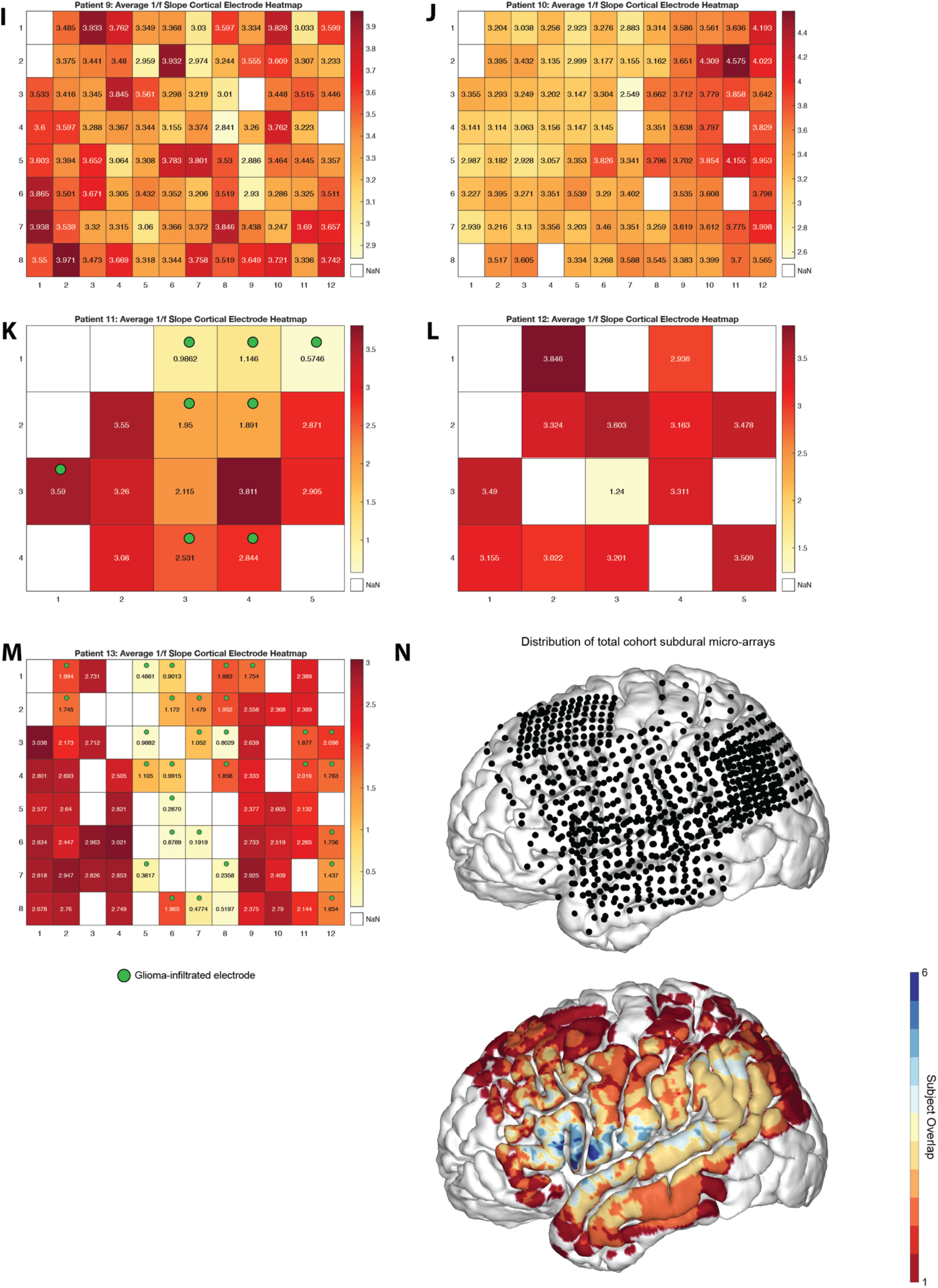
Heatmaps of average aperiodic exponent across cortical electrodes for individual patients. **A-M,** Electrode-wise average aperiodic exponent heatmaps for individual patients with high-density (8×12) and low-density (4×5) grids. Each panel represents the spatial distribution of average aperiodic exponent values across cortical electrodes for different patients, labeled by patient ID. The color scale represents the magnitude of the aperiodic exponent, with higher values shown in darker red and lower values in yellow. White squares indicate missing or excluded electrodes. The arrangement and variability of aperiodic exponents across electrodes reflect differences in spectral aperiodicity across individual patients. Green circle indicates values assigned to glioma-infiltrated electrodes. **N,** Electrode spatial locations mapped on an MNI brain. Top: Dots represent electrodes spatial distribution, Bottom: Heatmap of electrode spatial overlap across all patients.

**Extended Data Fig. 3.**
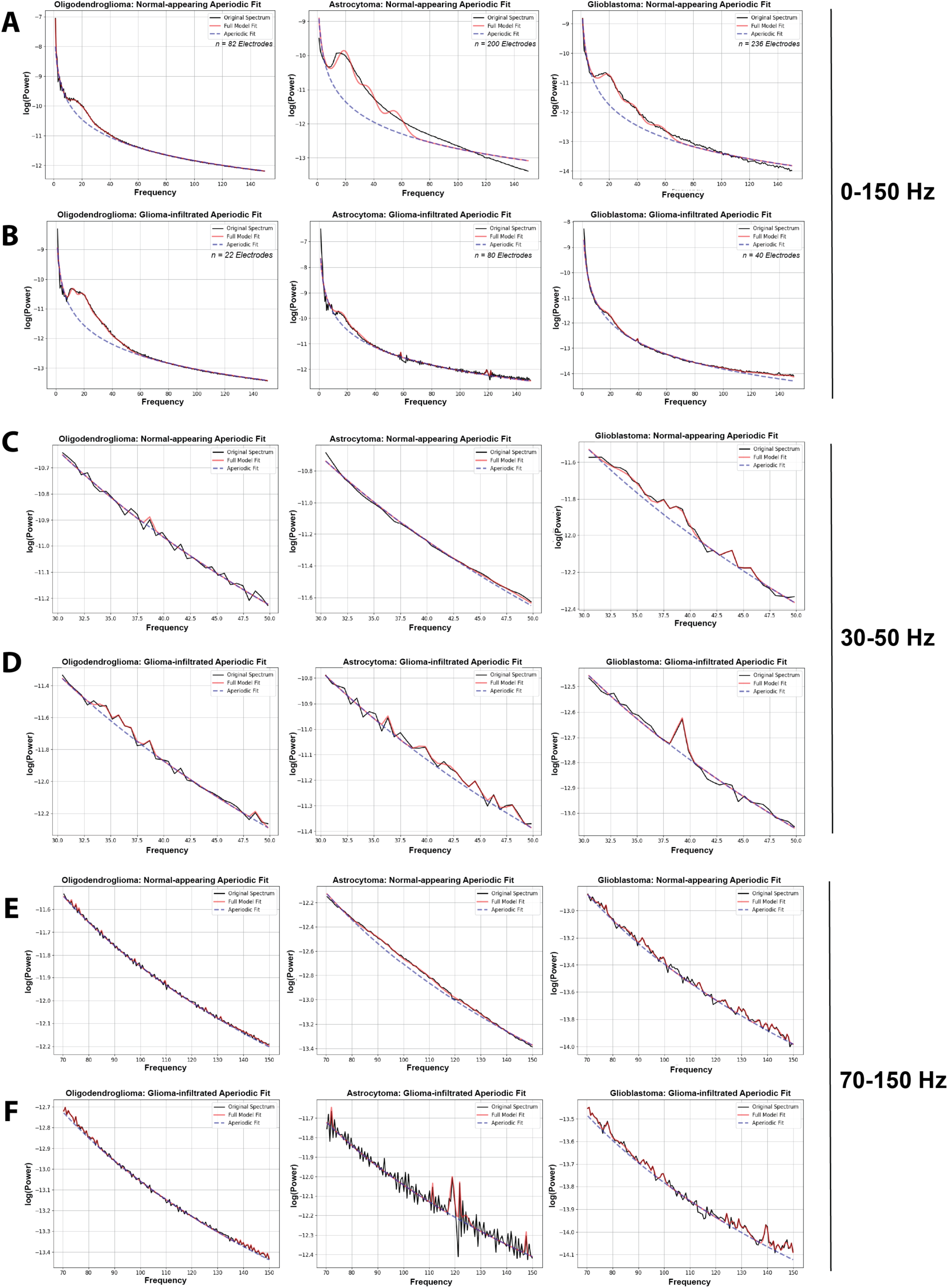
Aperiodic exponent differences and FOOOF-fit across glioma subtypes at 0-150 Hz, 30-50 Hz, and 70-150 Hz. **A-B**, Spectral analysis of normal-appearing (A) and glioma-infiltrated (B) electrode’s subdural ECoG data across glioma subtypes within the broadband (0-150 Hz) range. Left: oligodendroglioma (n=82 electrodes) middle: astrocytoma (n=200 electrodes), right: GBM (n=236 electrodes). The black line illustrates the original neural power spectrum, and the red line displays the full model fit. The aperiodic slope (or exponent) is represented by the blue dashed line. **C-D**, Aperiodic fit of normal-appearing (C) and glioma-infiltrated electrodes across glioma subtypes (Left to right: oligodendroglioma, astrocytoma, GBM) within the 30-50 Hz frequency range. **E-F**, Aperiodic fit of normal-appearing (E) and glioma-infiltrated (F) electrodes across glioma subtypes (Left to right: oligodendroglioma, astrocytoma, GBM) within 70-150 Hz.

**Extended Data Fig. 4.**
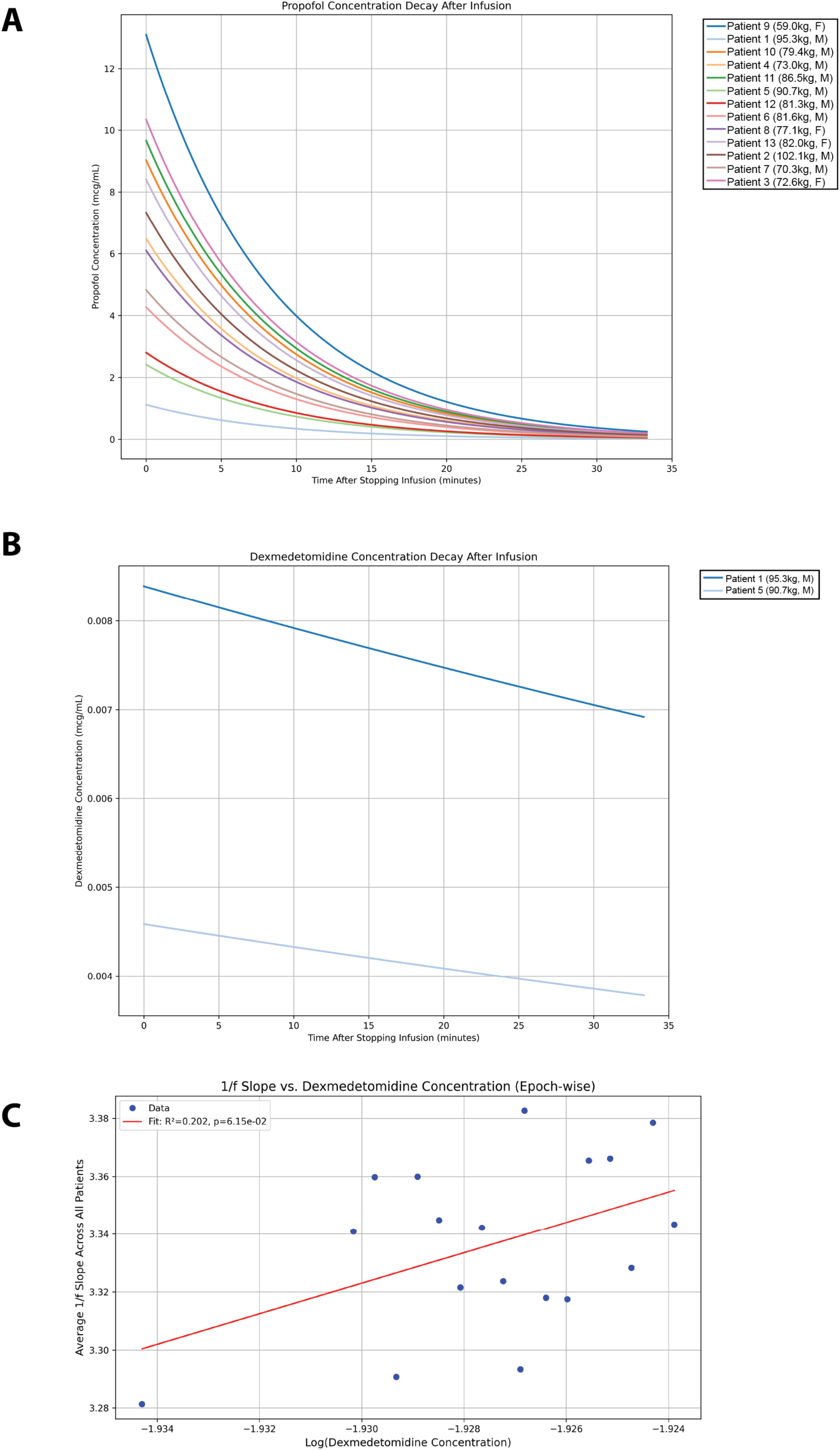
Pharmacokinetic profiles of propofol and dexmedetomidine following intravenous infusion. **A,** Time course of propofol plasma concentration decay (ng/ml) after cessation of intravenous infusion across 13 subjects. Each colored line represents an individual subject (subject ID with weight in kg and sex indicated in parentheses; M, male; F, female). **B,** Time course of dexmedetomidine plasma concentration decay (ng/ml) after cessation of intravenous infusion in two male subjects (patient 1, 95.3kg; patient 5, 90.7kg). **C,** Aperiodic exponent versus Dexmedetomidine concentration. Blue dots represent average aperiodic exponent across all patients computed in their respective concentration bin. Red line represents linear regression fit with an R² of 0.2017, p-value = 0.0615, and 95% CI = −0.2861 to 10.8029.

**Extended Data Fig. 5.**
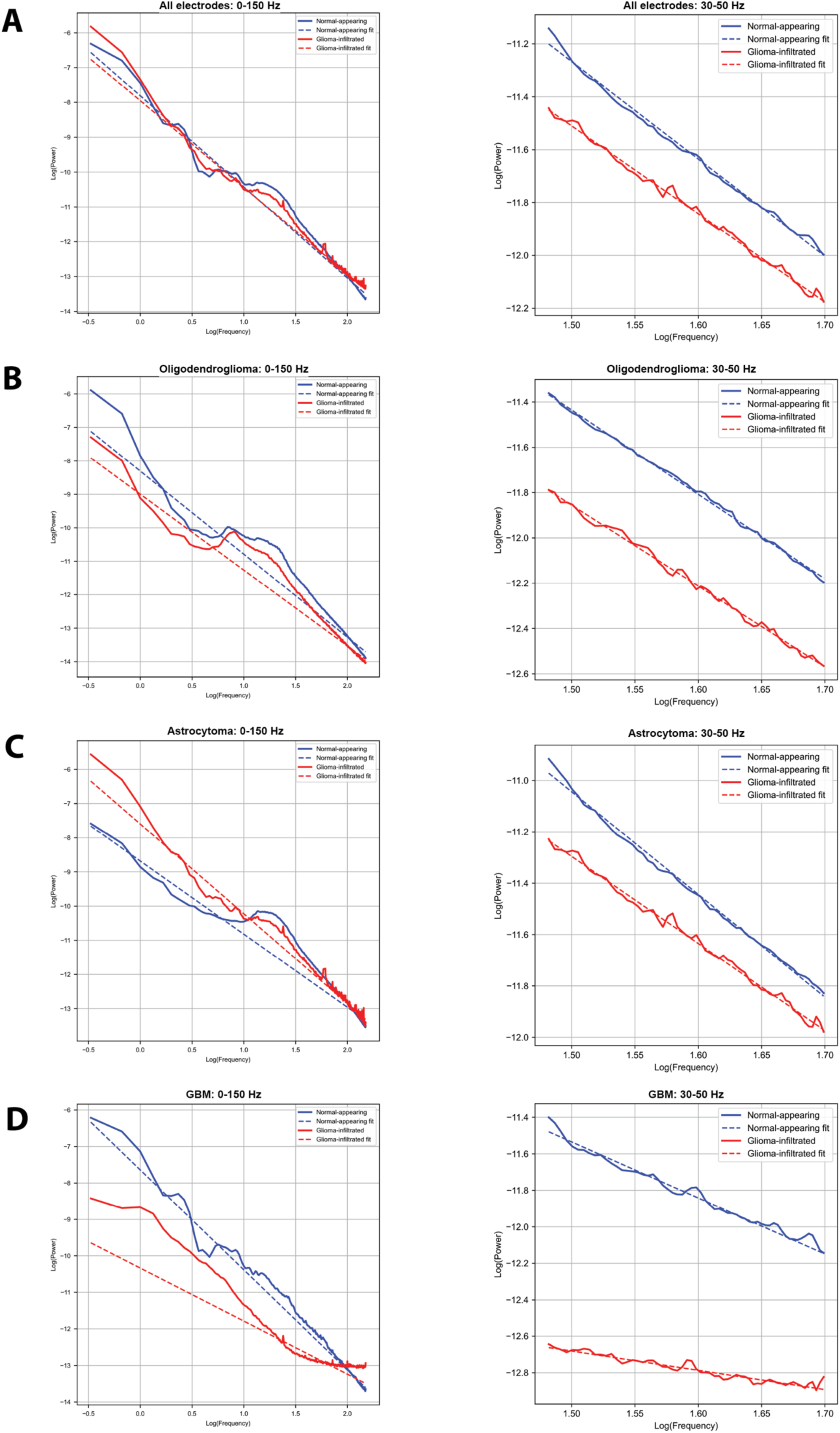
Power spectral density (PSD) comparisons and aperiodic fits between across all electrodes and glioma subtypes at 0-150 Hz and 30-50 Hz. **A,** Log-log aperiodic fits for all normal-appearing (blue) and all glioma-infiltrated (red) electrodes at 0-150 Hz (left) and 30-50 Hz (right). **B,** Oligodendroglioma log-log aperiodic fits for normal-appearing and glioma-infiltrated electrodes at 0-150 Hz (left) and 30-50 Hz (right). **C,** Astrocytoma log-log aperiodic fits for normal-appearing and glioma-infiltrated electrodes at 0-150 Hz (left) and 30-50 Hz (right). **D,** Glioblastoma log-log aperiodic fits for normal-appearing and glioma-infiltrated electrodes at 0-150 Hz (left) and 30-50 Hz (right).

**Extended Data Fig. 6.**
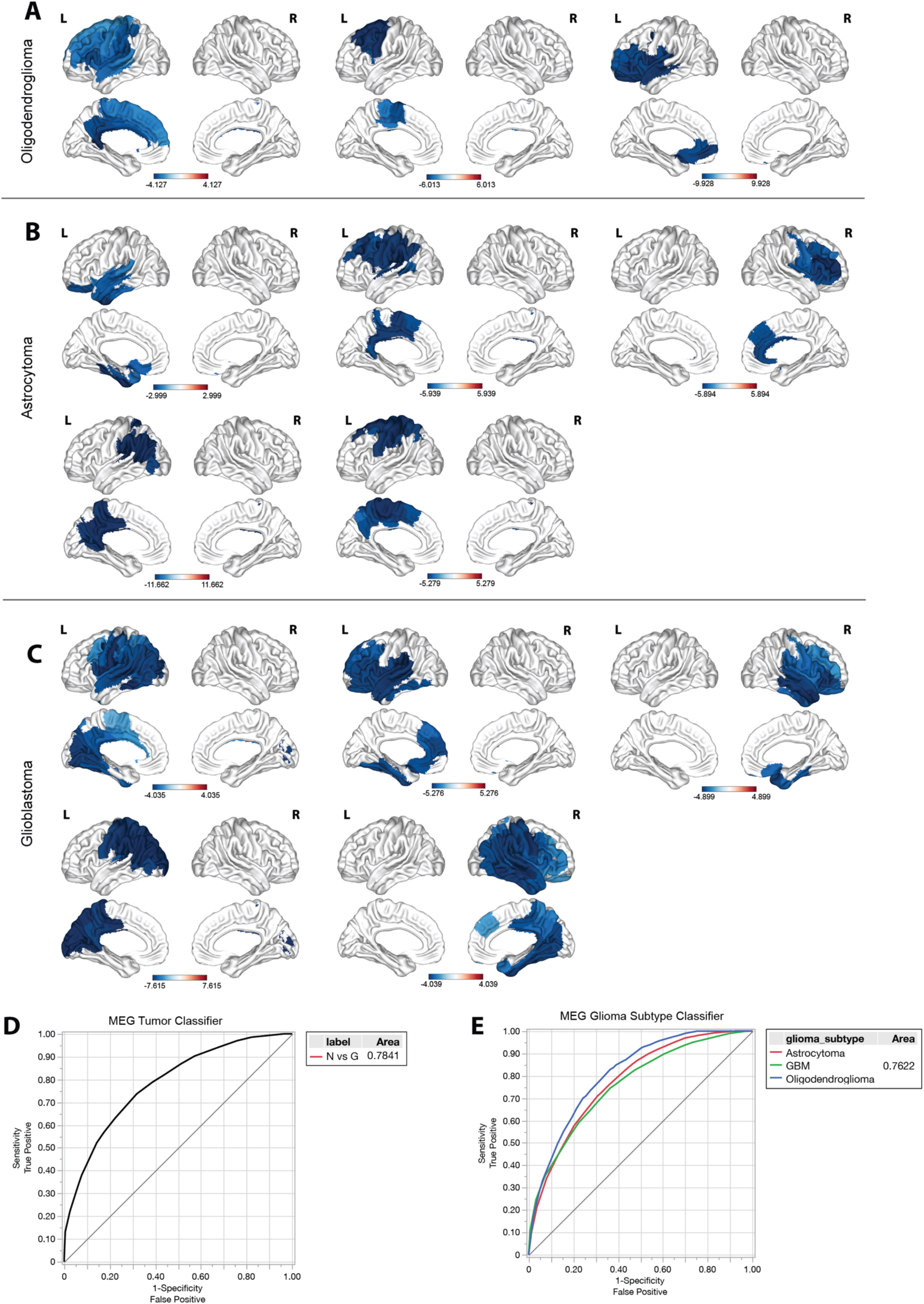
Individual z-statistical maps of MEG-derived aperiodic slope deviations across all patients and glioma subtypes. **A,** oligodendroglioma; **B,** astrocytoma; **C,** GBM. For each patient, aperiodic exponent values (30–50 Hz) were calculated at the voxel level and z-scored relative to anatomically homologous voxels in the contralateral hemisphere to account for individual baseline variability. Voxels with negative z-scores indicate local flattening of the aperiodic exponent in the tumor hemisphere relative to the non-tumor hemisphere, while positive z-scores reflect relative steepening. **D,** ROC curve for bootstrap random forest classifier predicting electrode tissue type. **E,** ROC curve for random forest classifier predicting glioma subtype using aperiodic exponent values. A “leave one patient out” cross-validation approach was utilized for each classifier.

**Extended Data Fig. 7.**
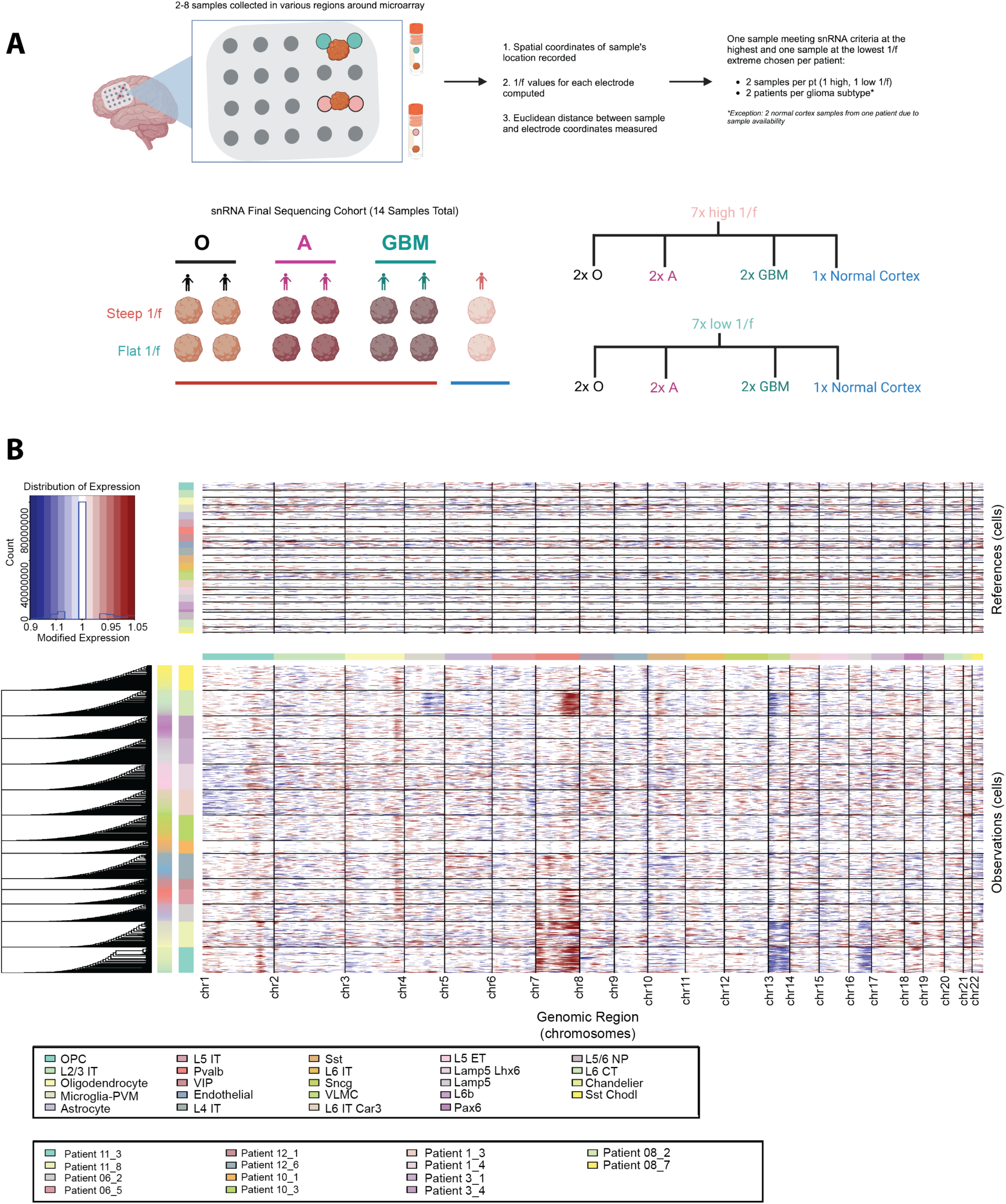
Tissue selection and snRNA-seq analysis of normal-appearing and glioma-infiltrated cortex. **A,** Schematic depicting the selection process for single-nucleus RNA sequencing (snRNA-seq) samples from glioma-infiltrated and normal cortex. Spatial coordinates of each sample were recorded, and aperiodic exponent values were computed for nearby electrodes. Euclidean distance mapping between samples and electrodes was measured to select one steep and one flat aperiodic exponent sample (at each extreme) per patient, ensuring equal representation across glioma subtypes. **B,** Heatmap of copy number variation (CNV) clusters across all 14 cortex samples (n=12 glioma-infiltrated, n=2 normal-appearing). Abbreviations: O = oligodendroglioma, A = astrocytoma, GBM = glioblastoma multiforme

**Extended Data Fig. 8.**
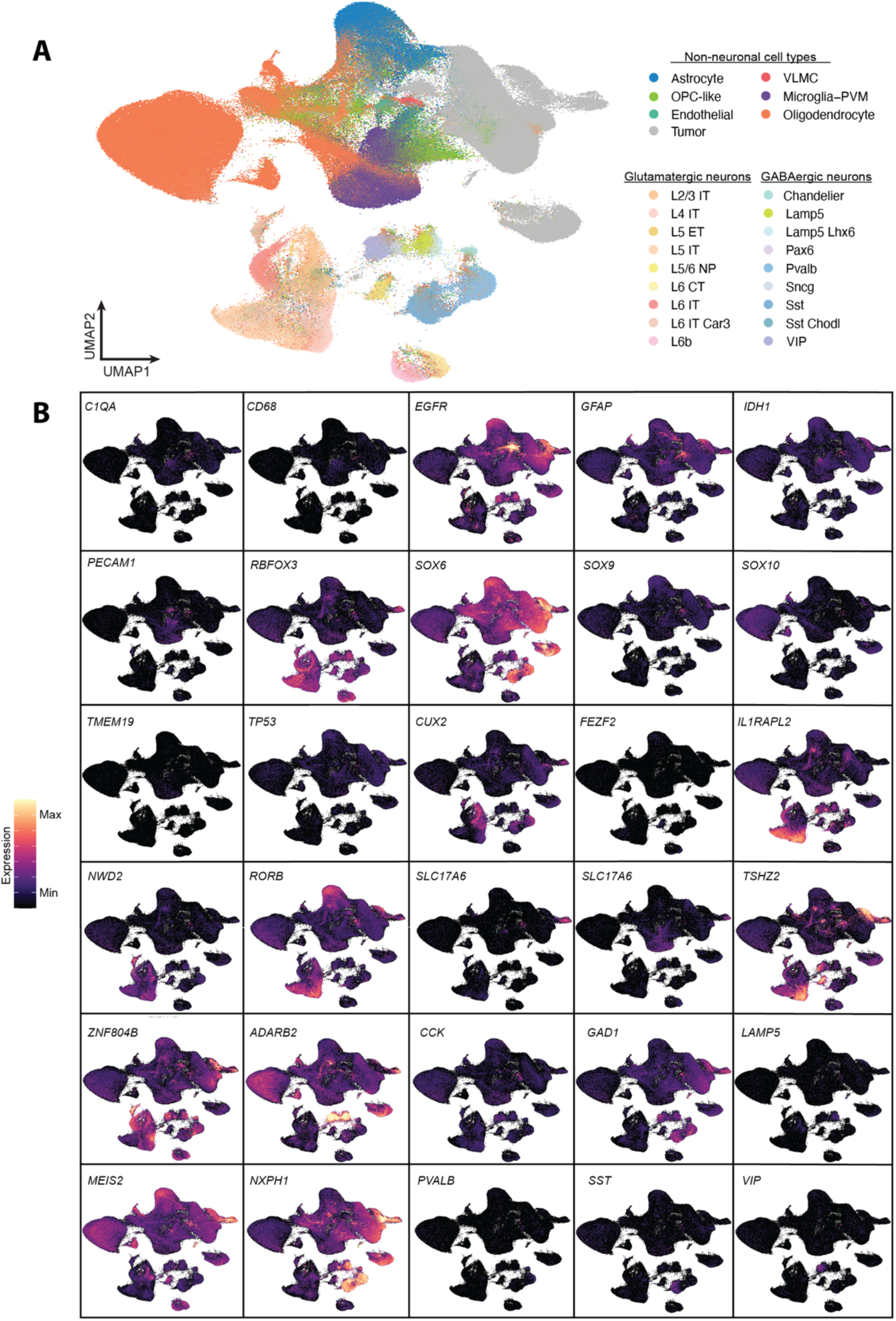
Non-neuronal and neuronal subpopulation transcriptomic cluster identities. **A,** Single-nucleus RNA sequencing UMAP of 508,899 transcriptomes from 14 human cortex samples across glioma-subtypes showing non-neuronal and neuronal population cell types. **B,** Feature plots displaying differentially expressed canonical marker genes across all clusters identified.

**Extended Data Fig. 9.**
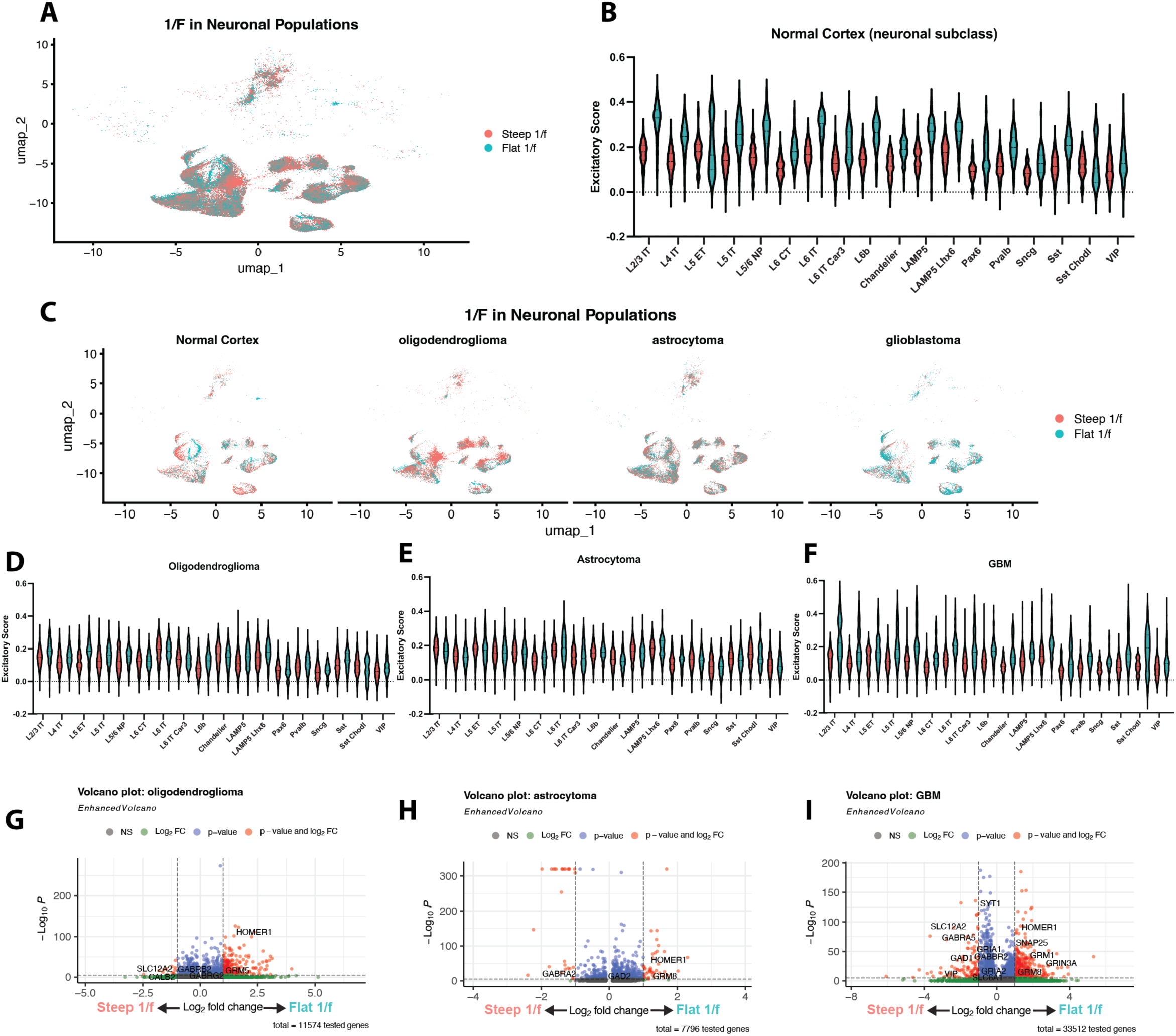
Cell type-specific transcriptomic differences between neuronal populations with steep versus flat aperiodic exponents across glioma subtypes. **A,** UMAP embedding of all neuronal populations from normal and glioma-infiltrated cortex, colored by steep (red) or flat (blue) aperiodic exponents based on matched ECoG recordings. **B,** Violin plots of excitatory module scores across neuronal subclasses in normal cortex, stratified by exponent group. Flat exponent neuronal populations (blue) demonstrate modestly higher excitatory scores across subclasses. **C,** UMAPs of neuronal populations stratified by tissue context: normal cortex, oligodendroglioma, astrocytoma, and glioblastoma. Distribution of steep and flat exponent neurons varies by glioma subtype. **D-F,** Violin plots showing excitatory score across neuronal subclasses within oligodendroglioma (D), astrocytoma (E), and glioblastoma (F) contexts, stratified by exponent group. **G-I,** Volcano plots of differential gene expression between steep and flat exponent neurons in oligodendroglioma (G), astrocytoma (H), and glioblastoma (I), highlighting key genes enriched in each group. Genes involved in synaptic signaling and excitation-inhibition balance (e.g., *SLC12A5, GABRA5, HOMER1*) are differentially expressed across neuronal populations in glioma-infiltrated cortex samples.

## Notes

### Competing Interest Statement

The authors have declared no competing interest.

