## Supplementary material for "Aperiodic neural dynamics define a novel signature of glioma-induced excitation-inhibition dysregulation": Sibih_Supplementary Figures

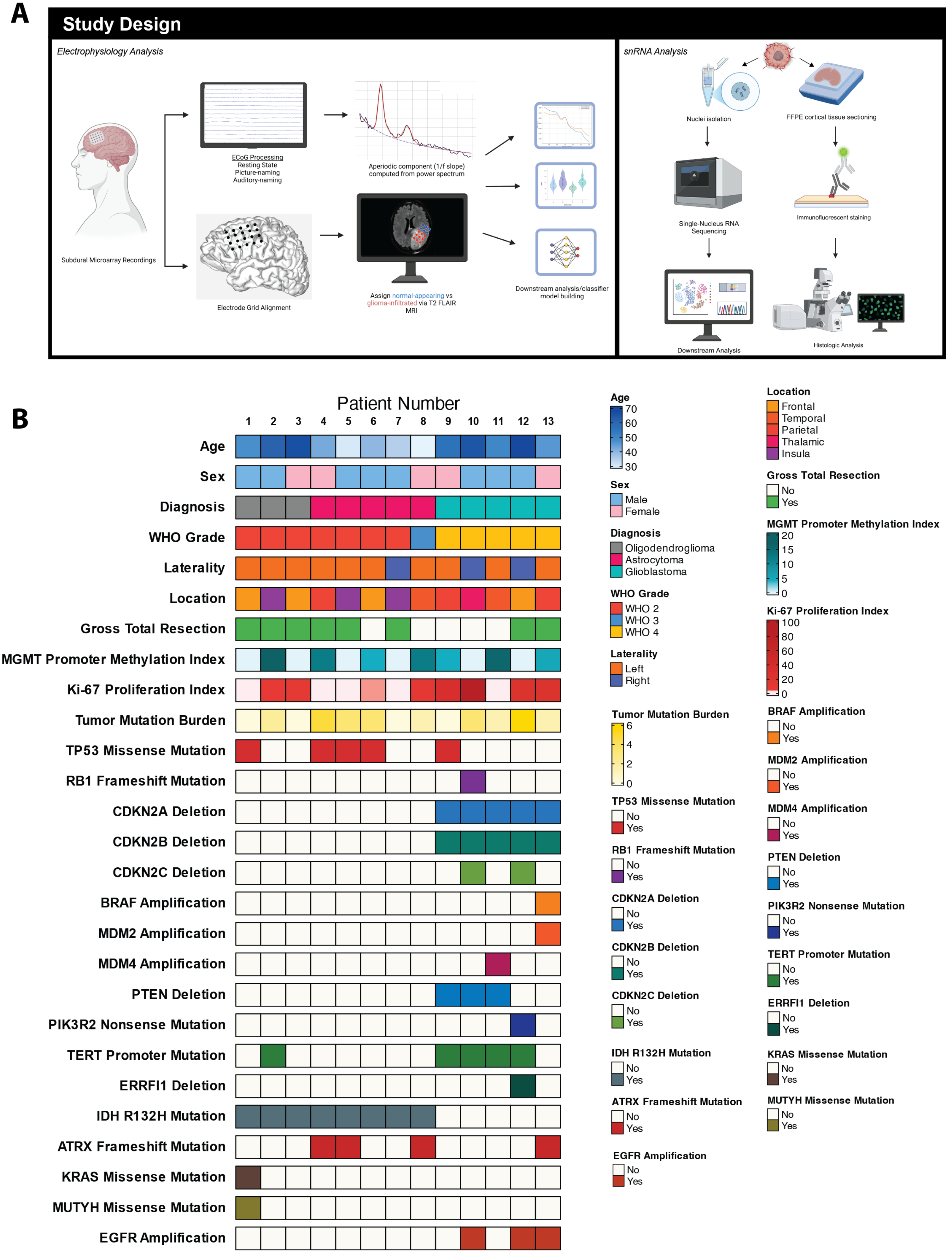
A
Study Design
Electrophysiology Analysis
Subdural Microarray Recordings
ECoG Processing
Resting State
Picture-naming
Auditory-naming
Aperiodic component (1/f slope) computed from power spectrum
Electrode Grid Alignment
Assign normal-appearing vs glioma-infiltrated via T2 FLAIR MRI
Downstream analysis/classifier model building
snRNA Analysis
Nuclei isolation
FFPE cortical tissue sectioning
Single-Nucleus RNA Sequencing
Immunofluorescent staining
Downstream Analysis
Histologic Analysis
B
Patient Number
1 2 3 4 5 6 7 8 9 10 11 12 13
Age
Sex
Diagnosis
WHO Grade
Laterality
Location
Gross Total Resection
MGMT Promoter Methylation Index
Ki-67 Proliferation Index
Tumor Mutation Burden
TP53 Missense Mutation
RB1 Frameshift Mutation
CDKN2A Deletion
CDKN2B Deletion
CDKN2C Deletion
BRAF Amplification
MDM2 Amplification
MDM4 Amplification
PTEN Deletion
PIK3R2 Nonsense Mutation
TERT Promoter Mutation
ERRFI1 Deletion
IDH R132H Mutation
ATRX Frameshift Mutation
KRAS Missense Mutation
MUTYH Missense Mutation
EGFR Amplification
Age
70
60
50
40
30
Sex
Male
Female
Diagnosis
Oligodendroglioma
Astrocytoma
Glioblastoma
WHO Grade
WHO 2
WHO 3
WHO 4
Laterality
Left
Right
Tumor Mutation Burden
6
4
2
0
TP53 Missense Mutation
No
Yes
RB1 Frameshift Mutation
No
Yes
CDKN2A Deletion
No
Yes
CDKN2B Deletion
No
Yes
CDKN2C Deletion
No
Yes
IDH R132H Mutation
No
Yes
ATRX Frameshift Mutation
No
Yes
EGFR Amplification
No
Yes
Location
Frontal
Temporal
Parietal
Thalamic
Insula
Gross Total Resection
No
Yes
MGMT Promoter Methylation Index
20
15
10
5
0
Ki-67 Proliferation Index
100
80
60
40
20
0
BRAF Amplification
No
Yes
MDM2 Amplification
No
Yes
MDM4 Amplification
No
Yes
PTEN Deletion
No
Yes
PIK3R2 Nonsense Mutation
No
Yes
TERT Promoter Mutation
No
Yes
ERRFI1 Deletion
No
Yes
KRAS Missense Mutation
No
Yes
MUTYH Missense Mutation
No
Yes

**Extended Data Fig. 1. Over of study design and patient histologic, molecular, and clinical features with complete list of next-generation sequencing targets. A,** Left: Electrophysiology analysis workflow. Right: Single-nucleus RNA sequencing analysis pipeline. **B,** Extended oncoprint displaying full list of targeted next-generation sequencing genes identified with targeted next-generation DNA sequencing.

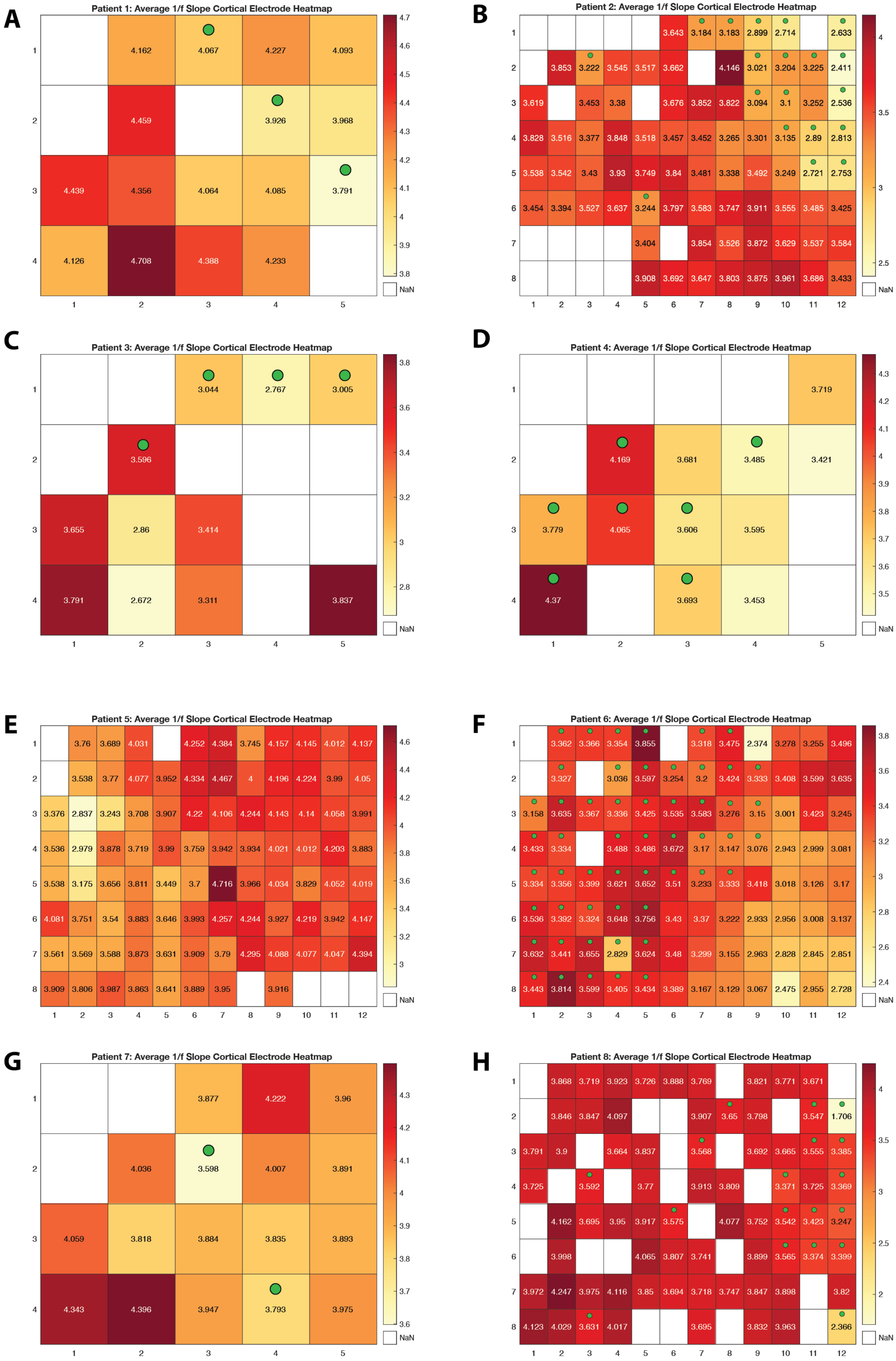
A
Patient 1: Average 1/f Slope Cortical Electrode Heatmap
B
Patient 2: Average 1/f Slope Cortical Electrode Heatmap
C
Patient 3: Average 1/f Slope Cortical Electrode Heatmap
D
Patient 4: Average 1/f Slope Cortical Electrode Heatmap
E
Patient 5: Average 1/f Slope Cortical Electrode Heatmap
F
Patient 6: Average 1/f Slope Cortical Electrode Heatmap
G
Patient 7: Average 1/f Slope Cortical Electrode Heatmap
H
Patient 8: Average 1/f Slope Cortical Electrode Heatmap
NaN

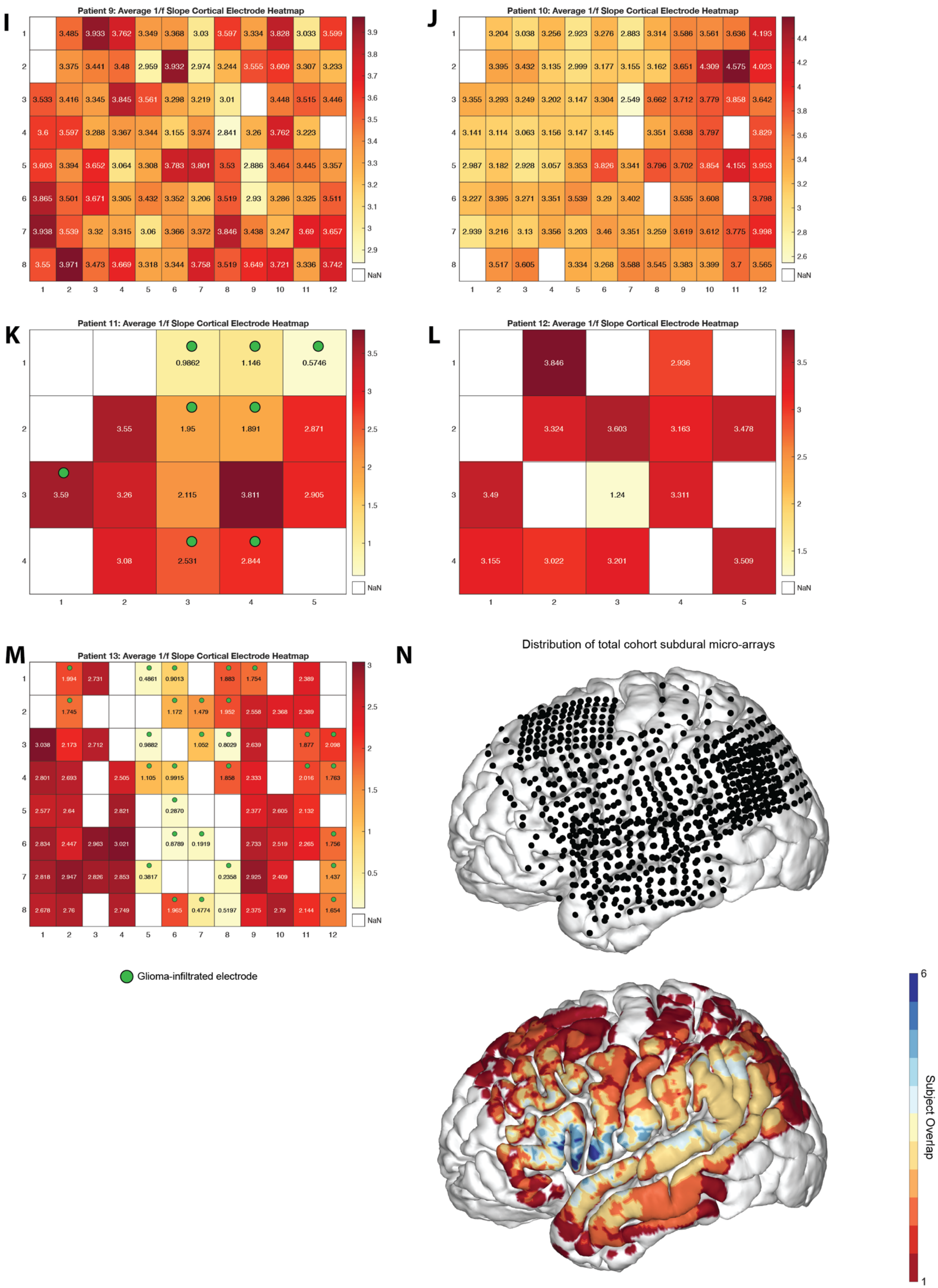
I
Patient 9: Average 1/f Slope Cortical Electrode Heatmap
3.485 3.933 3.762 3.349 3.368 3.03 3.597 3.334 3.828 3.033 3.599
3.375 3.441 3.48 2.959 3.932 2.974 3.244 3.555 3.609 3.307 3.233
3.533 3.416 3.345 3.845 3.561 3.298 3.219 3.01 3.448 3.515 3.446
3.6 3.597 3.288 3.367 3.344 3.155 3.374 2.841 3.26 3.762 3.223
3.603 3.394 3.652 3.064 3.308 3.783 3.801 3.53 2.886 3.464 3.445 3.357
3.865 3.501 3.671 3.305 3.432 3.352 3.206 3.519 2.93 3.286 3.325 3.511
3.938 3.539 3.32 3.315 3.06 3.366 3.372 3.846 3.438 3.247 3.69 3.657
3.55 3.971 3.473 3.669 3.318 3.344 3.758 3.519 3.649 3.721 3.336 3.742
NaN
J
Patient 10: Average 1/f Slope Cortical Electrode Heatmap
3.204 3.038 3.256 2.923 3.276 2.883 3.314 3.586 3.561 3.636 4.193
3.395 3.432 3.135 2.999 3.177 3.155 3.162 3.651 4.309 4.575 4.023
3.355 3.293 3.249 3.202 3.147 3.304 2.549 3.662 3.712 3.779 3.858 3.642
3.141 3.114 3.063 3.156 3.147 3.145 3.351 3.638 3.797 3.829
2.987 3.182 2.928 3.057 3.353 3.826 3.341 3.796 3.702 3.854 4.155 3.953
3.227 3.395 3.271 3.351 3.539 3.29 3.402 3.535 3.608 3.798
2.939 3.216 3.13 3.356 3.203 3.46 3.351 3.259 3.619 3.612 3.775 3.998
3.517 3.605 3.334 3.268 3.588 3.545 3.383 3.399 3.7 3.565
NaN
K
Patient 11: Average 1/f Slope Cortical Electrode Heatmap
0.9862 1.146 0.5746
3.55 1.95 1.891 2.871
3.59 3.26 2.115 3.811 2.905
3.08 2.531 2.844
NaN
L
Patient 12: Average 1/f Slope Cortical Electrode Heatmap
3.846 2.936
3.324 3.603 3.163 3.478
3.49 1.24 3.311
3.155 3.022 3.201 3.509
NaN
M
Patient 13: Average 1/f Slope Cortical Electrode Heatmap
1.994 2.731 0.4861 0.9013 1.883 1.754 2.389
1.745 1.172 1.479 1.952 2.558 2.368 2.389
3.038 2.173 2.712 0.9882 1.052 0.8029 2.639 1.877 2.098
2.801 2.693 2.505 1.105 0.9915 1.858 2.333 2.016 1.763
2.577 2.64 2.821 0.2870 2.377 2.605 2.132
2.834 2.447 2.963 3.021 0.8789 0.1919 2.733 2.519 2.265 1.756
2.818 2.947 2.826 2.853 0.3817 0.2358 2.925 2.409 1.437
2.678 2.76 2.749 1.965 0.4774 0.5197 2.375 2.79 2.144 1.654
NaN
Glioma-infiltrated electrode
N
Distribution of total cohort subdural micro-arrays
Subject Overlap
6
1

**Extended Data Fig. 2. Heatmaps of average aperiodic exponent across cortical electrodes for individual patients. A-M,** Electrode-wise average aperiodic exponent heatmaps for individual patients with high-density (8x12) and low-density (4x5) grids. Each panel represents the spatial distribution of average aperiodic exponent values across cortical electrodes for different patients, labeled by patient ID. The color scale represents the magnitude of the aperiodic exponent, with higher values shown in darker red and lower values in yellow. White squares indicate missing or excluded electrodes. The arrangement and variability of aperiodic exponents across electrodes reflect differences in spectral aperiodicity across individual patients. Green circle indicates values assigned to glioma-infiltrated electrodes. **N,** Electrode spatial locations mapped on an MNI brain. Top: Dots represent electrodes spatial distribution, Bottom: Heatmap of electrode spatial overlap across all patients.

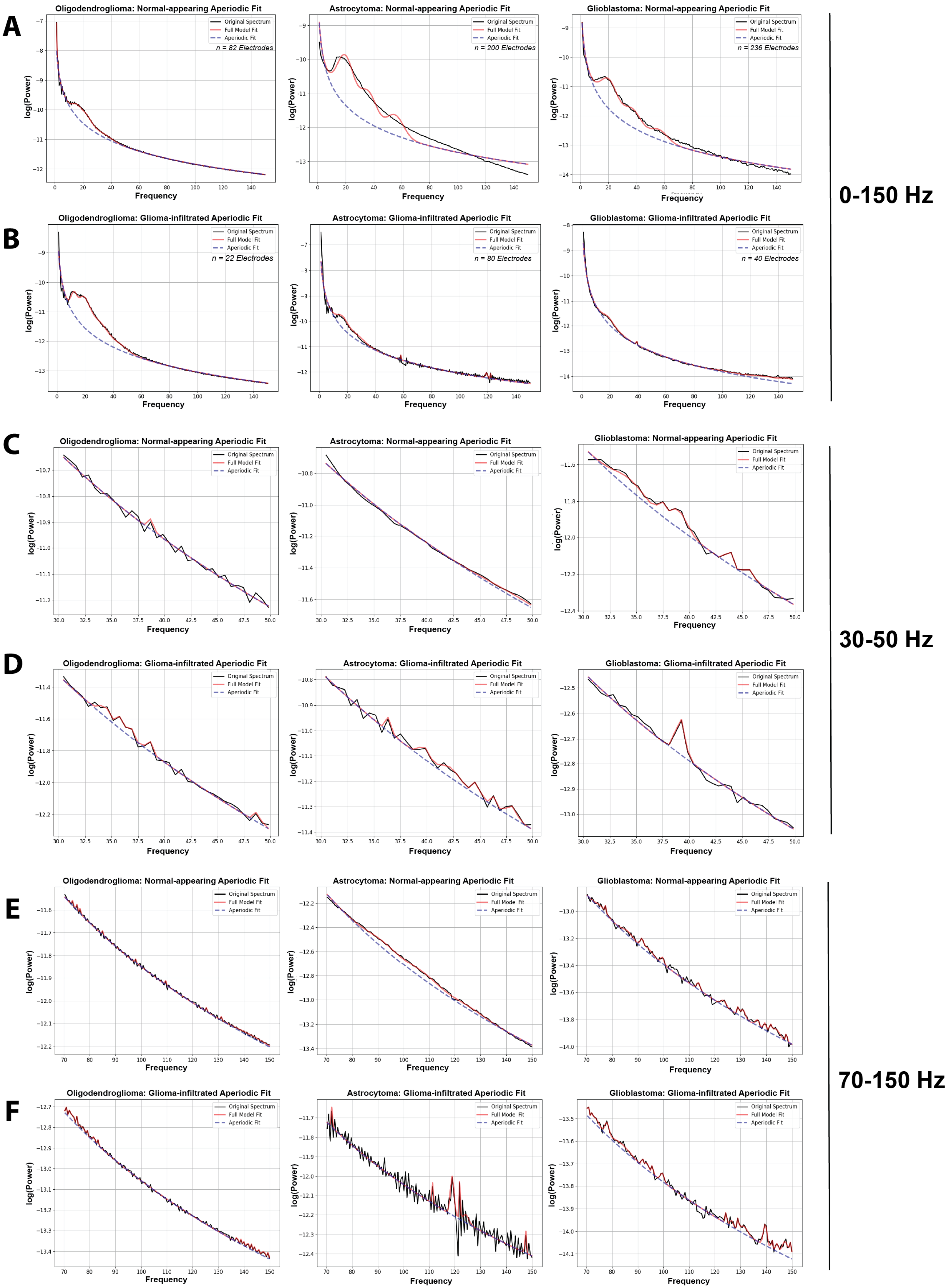
A
Oligodendroglioma: Normal-appearing Aperiodic Fit
n = 82 Electrodes
Astrocytoma: Normal-appearing Aperiodic Fit
n = 200 Electrodes
Glioblastoma: Normal-appearing Aperiodic Fit
n = 236 Electrodes
B
Oligodendroglioma: Glioma-infiltrated Aperiodic Fit
n = 22 Electrodes
Astrocytoma: Glioma-infiltrated Aperiodic Fit
n = 80 Electrodes
Glioblastoma: Glioma-infiltrated Aperiodic Fit
n = 40 Electrodes
0-150 Hz
C
Oligodendroglioma: Normal-appearing Aperiodic Fit
Astrocytoma: Normal-appearing Aperiodic Fit
Glioblastoma: Normal-appearing Aperiodic Fit
D
Oligodendroglioma: Glioma-infiltrated Aperiodic Fit
Astrocytoma: Glioma-infiltrated Aperiodic Fit
Glioblastoma: Glioma-infiltrated Aperiodic Fit
30-50 Hz
E
Oligodendroglioma: Normal-appearing Aperiodic Fit
Astrocytoma: Normal-appearing Aperiodic Fit
Glioblastoma: Normal-appearing Aperiodic Fit
F
Oligodendroglioma: Glioma-infiltrated Aperiodic Fit
Astrocytoma: Glioma-infiltrated Aperiodic Fit
Glioblastoma: Glioma-infiltrated Aperiodic Fit
70-150 Hz
Original Spectrum
Full Model Fit
Aperiodic Fit
log(Power)
Frequency

**Extended Data Fig. 3. Aperiodic exponent differences and FOOOF-fit across glioma subtypes at 0-150 Hz, 30-50 Hz, and 70-150 Hz. A-B**, Spectral analysis of normal-appearing (A) and glioma-infiltrated (B) electrode's subdural ECoG data across glioma subtypes within the broadband (0-150 Hz) range. Left: oligodendroglioma (n=82 electrodes) middle: astrocytoma (n=200 electrodes), right: GBM (n=236 electrodes). The black line illustrates the original neural power spectrum and the red line displays the full model fit. The aperiodic slope (or exponent) is represented by the blue dashed line. **C-D**, Aperiodic fit of normal-appearing (C) and glioma-infiltrated electrodes across glioma subtypes (Left to right: oligodendroglioma, astrocytoma, GBM) within the 30-50 Hz frequency range. **E-F**, Aperiodic fit of normal-appearing (E) and glioma-infiltrated (F) electrodes across glioma subtypes (Left to right: oligodendroglioma, astrocytoma, GBM) within 70-150 Hz.

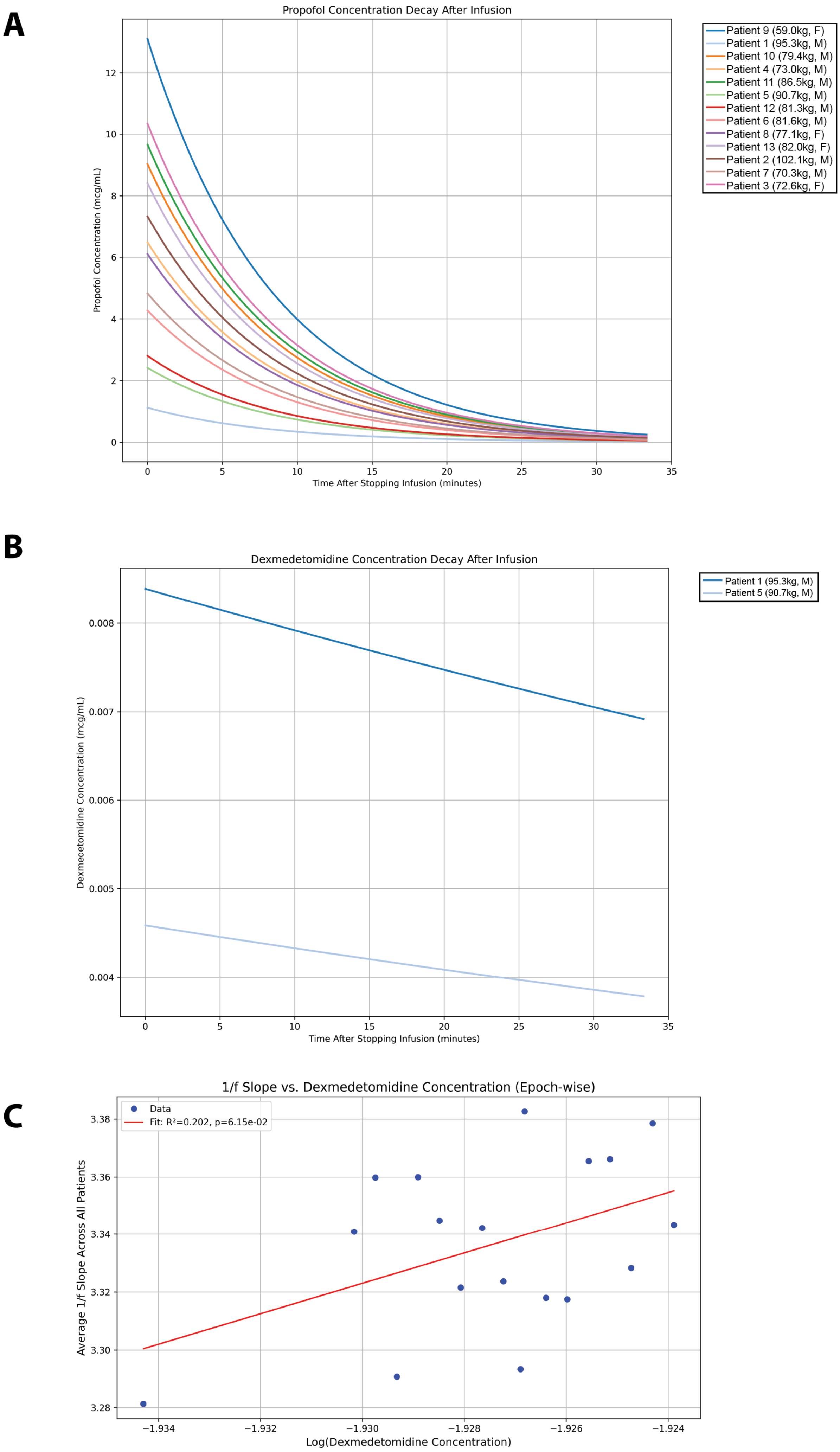
A
Propofol Concentration Decay After Infusion
Propofol Concentration (mcg/mL)
Time After Stopping Infusion (minutes)
Patient 9 (59.0kg, F)
Patient 1 (95.3kg, M)
Patient 10 (79.4kg, M)
Patient 4 (73.0kg, M)
Patient 11 (86.5kg, M)
Patient 5 (90.7kg, M)
Patient 12 (81.3kg, M)
Patient 6 (81.6kg, M)
Patient 8 (77.1kg, F)
Patient 13 (82.0kg, F)
Patient 2 (102.1kg, M)
Patient 7 (70.3kg, M)
Patient 3 (72.6kg, F)
B
Dexmedetomidine Concentration Decay After Infusion
Dexmedetomidine Concentration (mcg/mL)
Time After Stopping Infusion (minutes)
Patient 1 (95.3kg, M)
Patient 5 (90.7kg, M)
C
1/f Slope vs. Dexmedetomidine Concentration (Epoch-wise)
Data
Fit: R²=0.202, p=6.15e-02
Average 1/f Slope Across All Patients
Log(Dexmedetomidine Concentration)

**Extended Data Fig. 4. Pharmacokinetic profiles of propofol and dexmedetomidine following intravenous infusion. A,** Time course of propofol plasma concentration decay (ng/ml) after cessation of intravenous infusion across 13 subjects. Each colored line represents an individual subject (subject ID with weight in kg and sex indicated in parentheses; M, male; F, female). **B,** Time course of dexmedetomidine plasma concentration decay (ng/ml) after cessation of intravenous infusion in two male subjects (patient 1, 95.3kg; patient 5, 90.7kg). **C,** Aperiodic exponent versus Dexmedetomidine concentration. Blue dots represent average aperiodic exponent across all cohorts computed in their respective concentration bin. Red line represents linear regression fit with an $R^2$ of 0.2017, p-value = 0.0615, and 95% CI = -0.2861 to 10.8029.

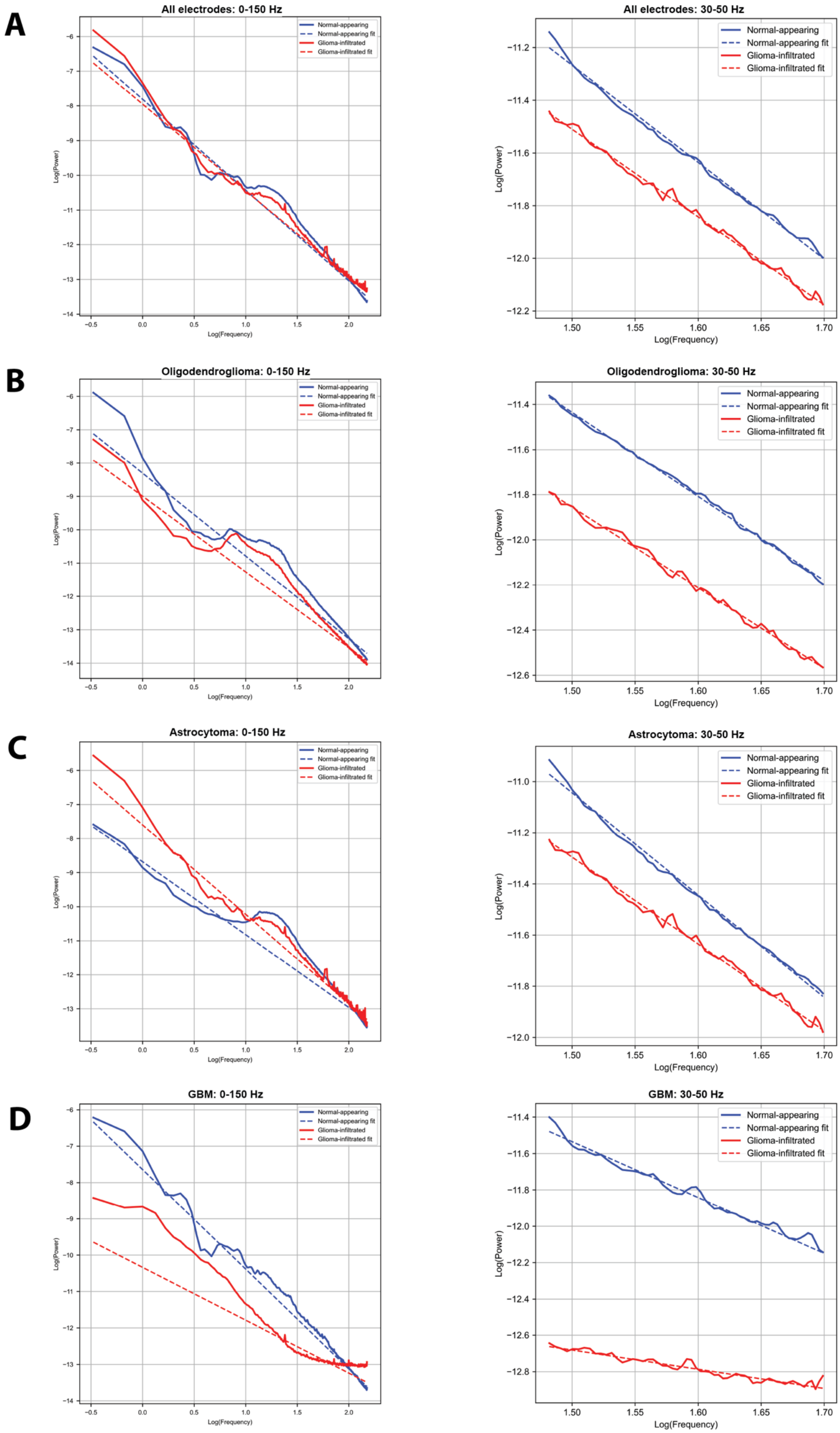

A
All electrodes: 0-150 Hz
All electrodes: 30-50 Hz
B
Oligodendroglioma: 0-150 Hz
Oligodendroglioma: 30-50 Hz
C
Astrocytoma: 0-150 Hz
Astrocytoma: 30-50 Hz
D
GBM: 0-150 Hz
GBM: 30-50 Hz
Normal-appearing
Normal-appearing fit
Glioma-infiltrated
Glioma-infiltrated fit
Log(Power)
Log(Frequency)

**Extended Data Fig. 5. Power spectral density (PSD) comparisons and aperiodic fits between across all electrodes and glioma subtypes at 0-150 Hz and 30-50 Hz. A,** Log-log aperiodic fits for all normal-appearing (blue) and all glioma-infiltrated (red) electrodes at 0-150 Hz (left) and 30-50 Hz (right). **B,** Oligodendroglioma log-log aperiodic fits for normal-appearing and glioma-infiltrated electrodes at 0-150 Hz (left) and 30-50 Hz (right). **C,** Astrocytoma log-log aperiodic fits for normal-appearing and glioma-infiltrated electrodes at 0-150 Hz (left) and 30-50 Hz (right). **D,** Glioblastoma log-log aperiodic fits for normal-appearing and glioma-infiltrated electrodes at 0-150 Hz (left) and 30-50 Hz (right).

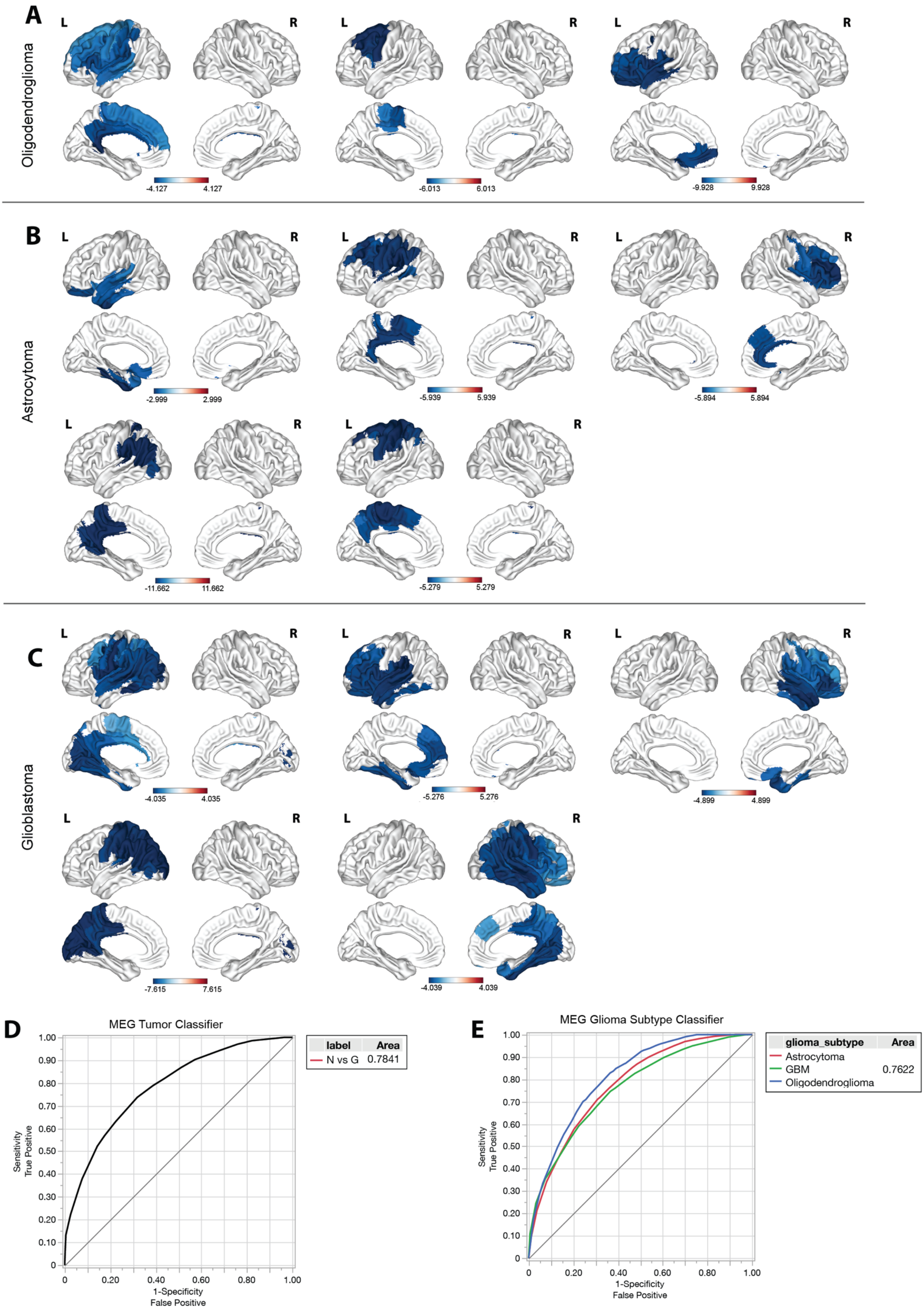

A
Oligodendroglioma
L
R
-4.127
4.127
-6.013
6.013
-9.928
9.928
B
Astrocytoma
-2.999
2.999
-5.939
5.939
-5.894
5.894
-11.662
11.662
-5.279
5.279
C
Glioblastoma
-4.035
4.035
-5.276
5.276
-4.899
4.899
-7.615
7.615
-4.039
4.039
D
MEG Tumor Classifier
Sensitivity
True Positive
1-Specificity
False Positive
label
Area
N vs G
0.7841
E
MEG Glioma Subtype Classifier
glioma_subtype
Area
Astrocytoma
GBM
0.7622
Oligodendroglioma

**Extended Data Fig. 6. Individual z-statistical maps of MEG-derived aperiodic slope deviations across all patients and glioma subtypes. A,** oligodendroglioma; **B,** astrocytoma; **C,** GBM. For each patient, aperiodic exponent values (30–50 Hz) were calculated at the voxel level and z-scored relative to anatomically homologous voxels in the contralateral hemisphere to account for individual baseline variability. Voxels with negative z-scores indicate local flattening of the aperiodic exponent in the tumor hemisphere relative to the non-tumor hemisphere, while positive z-scores reflect relative steepening. **D,** ROC curve for bootstrap random forest classifier predicting electrode tissue type. **E,** ROC curve for random forest classifier predicting glioma subtype using aperiodic exponent values. A "leave one patient out" cross-validation approach was utilized for each classifier.

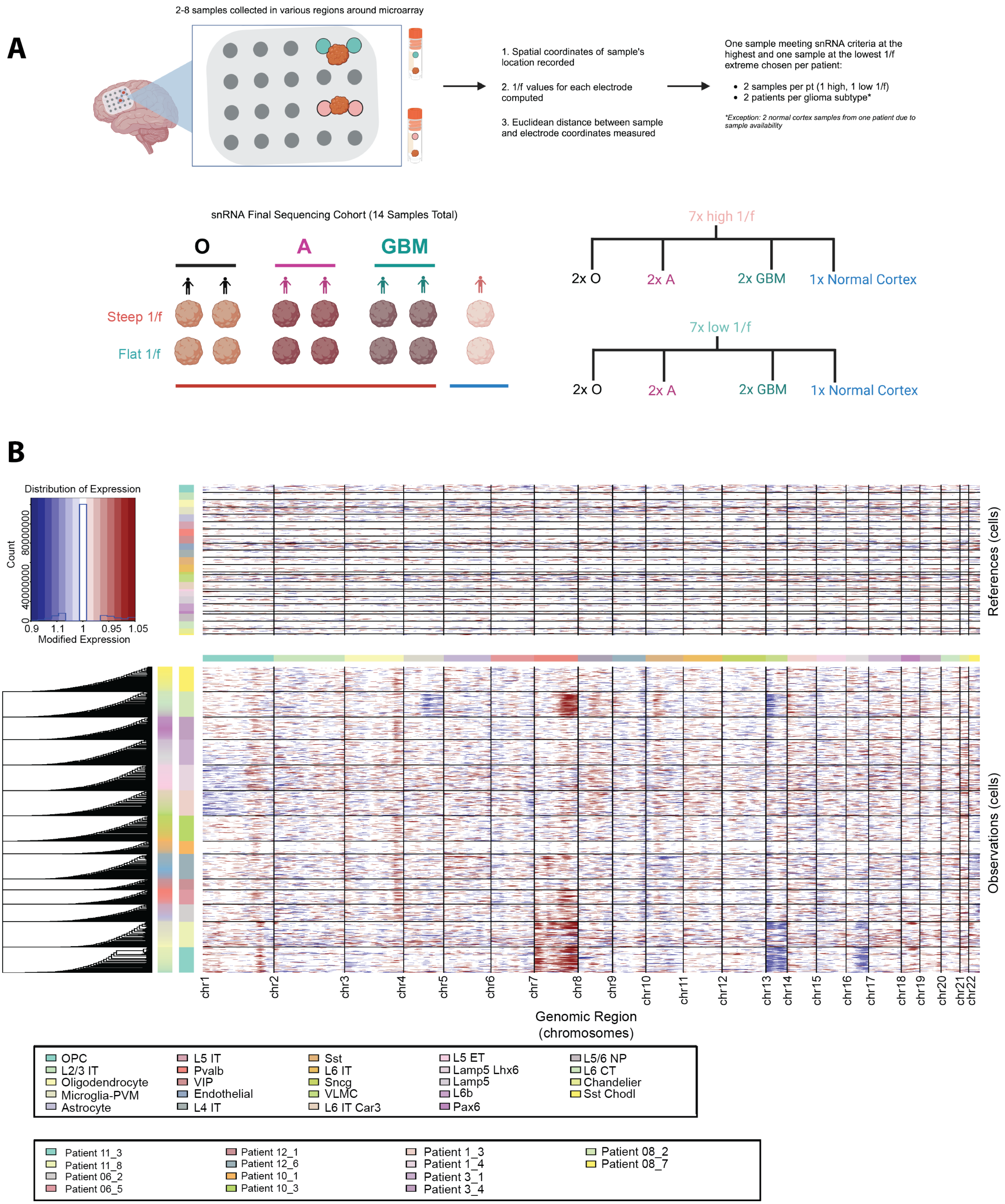
A
2-8 samples collected in various regions around microarray
1. Spatial coordinates of sample's location recorded
2. 1/f values for each electrode computed
3. Euclidean distance between sample and electrode coordinates measured
One sample meeting snRNA criteria at the highest and one sample at the lowest 1/f extreme chosen per patient:
• 2 samples per pt (1 high, 1 low 1/f)
• 2 patients per glioma subtype*
*Exception: 2 normal cortex samples from one patient due to sample availability
snRNA Final Sequencing Cohort (14 Samples Total)
O
A
GBM
Steep 1/f
Flat 1/f
7x high 1/f
2x O
2x A
2x GBM
1x Normal Cortex
7x low 1/f
2x O
2x A
2x GBM
1x Normal Cortex
B
Distribution of Expression
Count
0
40000000
80000000
0.9
1.1
1
0.95
1.05
Modified Expression
References (cells)
Observations (cells)
chr1
chr2
chr3
chr4
chr5
chr6
chr7
chr8
chr9
chr10
chr11
chr12
chr13
chr14
chr15
chr16
chr17
chr18
chr19
chr20
chr21
chr22
Genomic Region (chromosomes)
OPC
L2/3 IT
Oligodendrocyte
Microglia-PVM
Astrocyte
L5 IT
Pvalb
VIP
Endothelial
L4 IT
Sst
L6 IT
Sncg
VLMC
L6 IT Car3
L5 ET
Lamp5 Lhx6
Lamp5
L6b
Pax6
L5/6 NP
L6 CT
Chandelier
Sst Chodl
Patient 11_3
Patient 11_8
Patient 06_2
Patient 06_5
Patient 12_1
Patient 12_6
Patient 10_1
Patient 10_3
Patient 1_3
Patient 1_4
Patient 3_1
Patient 3_4
Patient 08_2
Patient 08_7

**Extended Data Fig. 7. Tissue selection and snRNA-seq analysis of normal-appearing and glioma-infiltrated cortex. A,** Schematic depicting the selection process for single-nucleus RNA sequencing (snRNA-seq) samples from glioma-infiltrated and normal cortex. Spatial coordinates of each sample were recorded, and aperiodic exponent values were computed for nearby electrodes. Euclidean distance mapping between samples and electrodes was measured to select one steep and one flat aperiodic exponent sample (at each extreme) per patient, ensuring equal representation across glioma subtypes. **B,** Heatmap of copy number variation (CNV) clusters across all 14 cortex samples (n=12 glioma-infiltrated, n=2 normal-appearing).

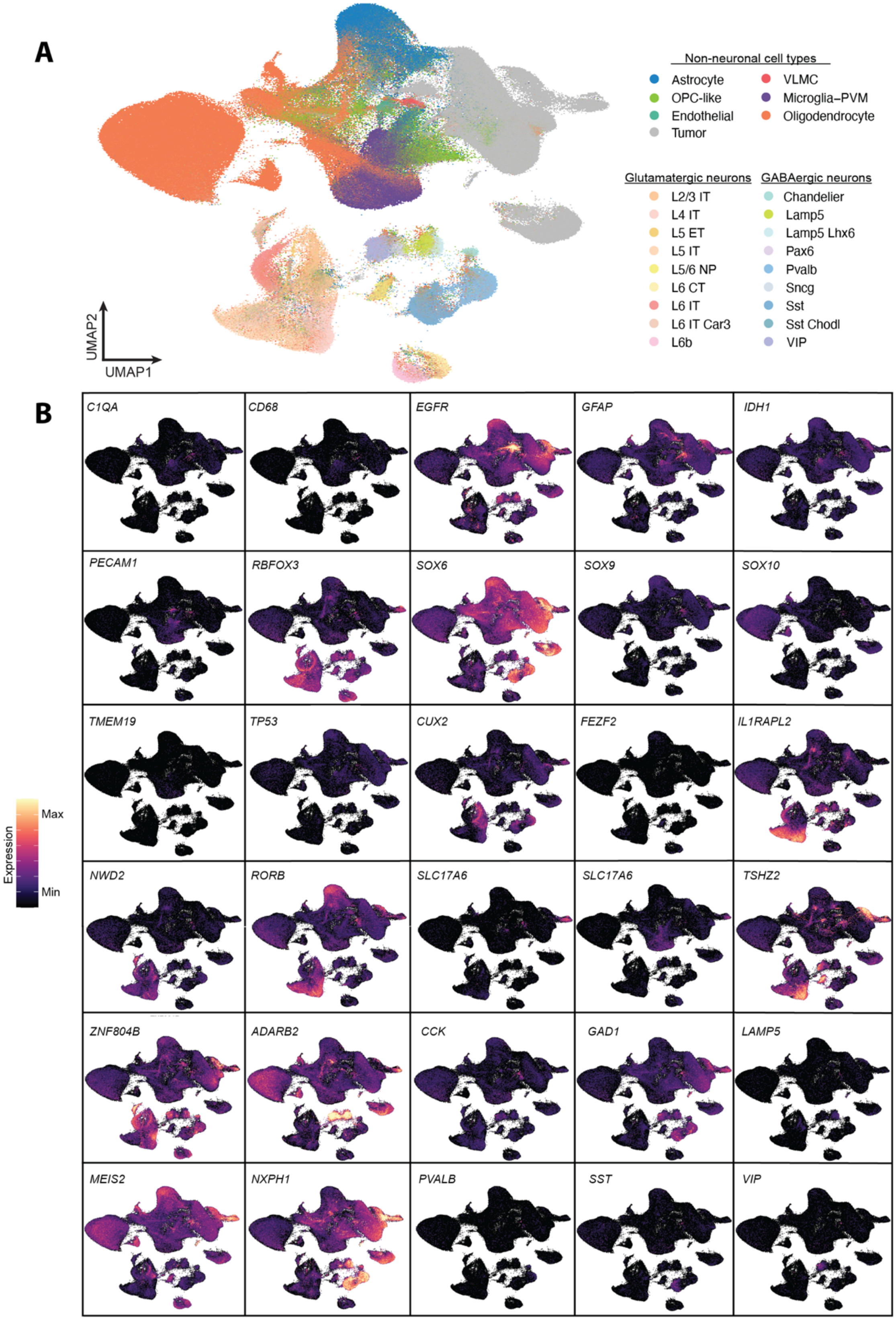


**Extended Data Fig. 8. Non-neuronal and neuronal subpopulation transcriptomic cluster identities.** **A,** Single-nucleus RNA sequencing UMAP of 508,899 transcriptomes from 14 human cortex samples across glioma-subtypes showing non-neuronal and neuronal population cell types. **B,** Feature plots displaying differentially expressed canonical marker genes across all clusters identified.

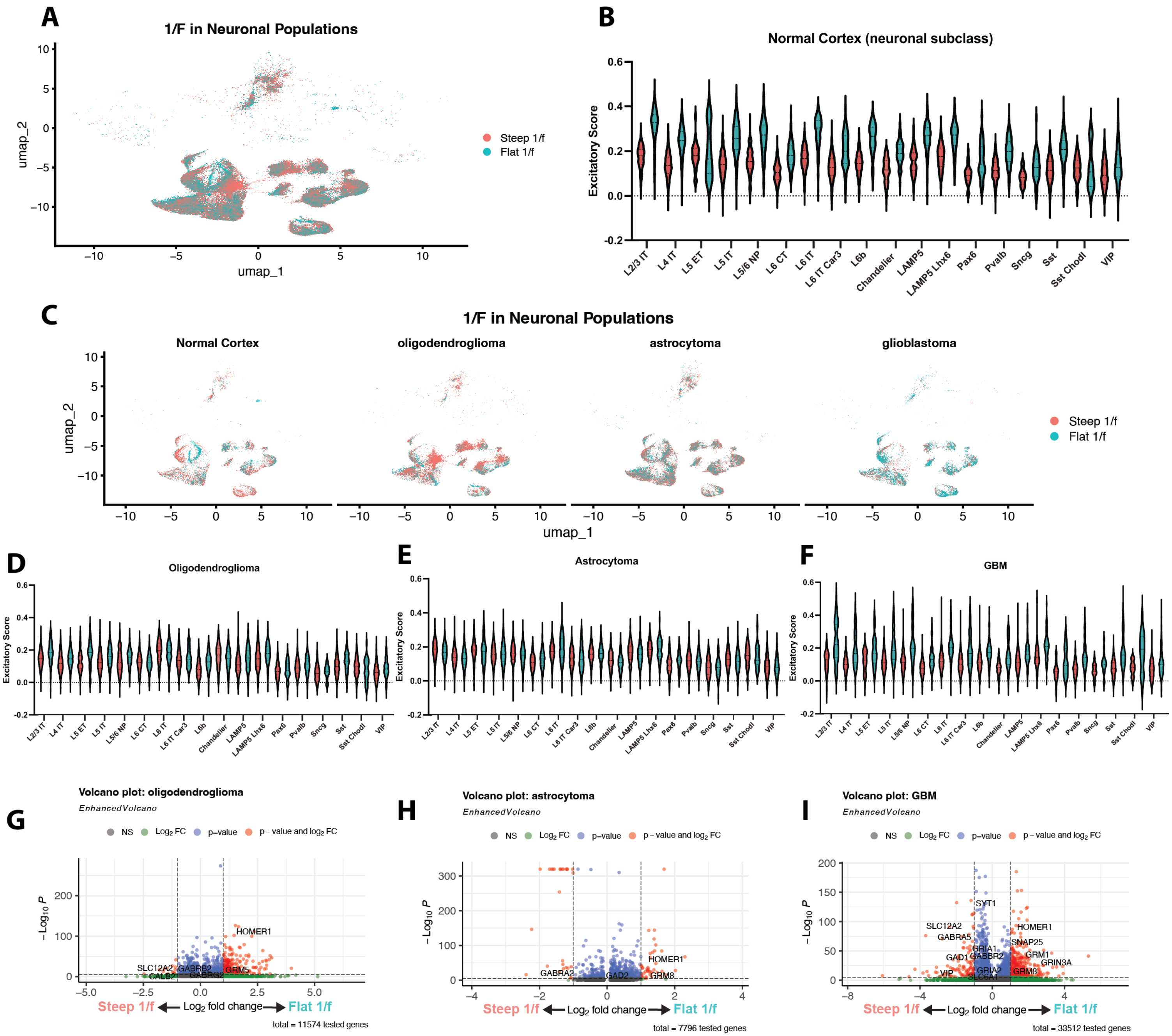


**Extended Data Fig. 9. Cell type-specific transcriptomic differences between neuronal populations with steep versus flat aperiodic exponents across glioma subtypes. A,** UMAP embedding of all neuronal populations from normal and glioma-infiltrated cortex, colored by steep (red) or flat (blue) aperiodic exponents based on matched ECoG recordings. **B,** Violin plots of excitatory module scores across neuronal subclasses in normal cortex, stratified by exponent group. Flat exponent neuronal populations (blue) demonstrate modestly higher excitatory scores across subclasses. **C,** UMAPs of neuronal populations stratified by tissue context: normal cortex, oligodendroglioma, astrocytoma, and glioblastoma. Distribution of steep and flat exponent neurons varies by glioma subtype. **D-F,** Violin plots showing excitatory score across neuronal subclasses within oligodendroglioma (D), astrocytoma (E), and glioblastoma (F) contexts, stratified by exponent group. **G-I,** Volcano plots of differential gene expression between steep and flat exponent neurons in oligodendroglioma (G), astrocytoma (H), and glioblastoma (I), highlighting key genes enriched in each group. Genes involved in synaptic signaling and excitation-inhibition balance (e.g., *SLC12A5, GABRA5, HOMER1*) are differentially expressed across neuronal populations in glioma-infiltrated cortex samples.
